# Triple-effect correction for Cell Painting data with contrastive and domain-adversarial learning

**DOI:** 10.1101/2025.01.13.632874

**Authors:** Chengwei Yan, Yu Zhang, Jiuxin Feng, Heyang Hua, Zhihan Ruan, Zhen Li, Siyu Li, Chaoyang Yan, Pingjing Li, Jian Liu, Shengquan Chen

## Abstract

Cell Painting (CP), as a high-throughput imaging technology, generates extensive cell-stained imaging data, providing unique morphological insights for biological research. However, CP data contains three types of technical effects, referred to as triple effects, including batch effects, gradient-influenced row and column effects (well position effects). The interaction of various technical effects can obscure true biological signals and complicate the characterization of CP data, making correction essential for reliable analysis. Here, we propose cpDistiller, a triple-effect correction method specially designed for CP data, which leverages a pre-trained segmentation model coupled with a semi-supervised Gaussian mixture variational autoencoder utilizing contrastive and domain-adversarial learning. Through extensive qualitative and quantitative experiments across various CP profiles, we demonstrate that cpDistiller effectively corrects triple effects, especially well position effects, a challenge that no current methods address, while preserving cellular heterogeneity. Moreover, cpDistiller effectively captures system-level phenotypic responses to genetic perturbations and reliably infers gene functions and interactions both when combined with scRNA-seq data and independently. cpDistiller also excels at identifying gene and compound targets, which is a critical step in drug discovery and broader biological research.

---

Advanced high-dimensional assay technologies, such as transcriptomics and epigenomics profiling, offer remarkable depth and breadth in molecular-level biological research^1^. Despite their strengths, these technologies often focus exclusively on specific molecular changes, lacking the capability to observe changes at the system level of cell state which involves many complex and unknown processes. To obtain information at the cellular system level, high-throughput imaging technologies have been developed to produce useful profiles of cell phenotypes by imaging stained cells^2–4^. However, these image-based technologies also have their limitations, as they typically focus on biological processes with known associations or assumptions, thereby constraining the discovery in existing knowledge^5^. Moreover, traditional methods that include both high-dimensional assay and image-based technologies are often constrained by their complexity and high costs. To overcome these issues, the technology, known as Cell Painting (CP), has been proposed as a solution. Specifically, CP technology involves staining eight cellular components with six remarkably cheap and easy dyes and imaging them in five channels on a fluorescence microscope^6^, which is simple to operate and less costly^7^. Beyond ease of use, CP technology operates on a new paradigm by collecting data on a large scale without an initial focus on specific hypotheses or known knowledge to reveal unanticipated biology at play. With these advantages, CP technology has shed light on novel phenotypes and cellular phenotypic heterogeneity, providing a valuable complement to genomics^8^. It also has been successfully used to characterize genes^9–11^ and compounds^2,11,12^ in several steps of the drug discovery process.

As CP technology has developed and been increasingly applied, a large amount of data has been accumulated^13^. In 2023, scientists established the Joint Undertaking for Morphological Profiling (JUMP) dataset, a standardized collection of cell-stained images featuring over 116,000 unique compound perturbations and more than 15,000 unique genetic perturbations^14^. Despite the dataset’s substantial analytical potential, technical effects caused by non-biological factors pose significant challenges in the analysis process. Some of these technical effects, resulting from variations across different laboratories and different batches in a laboratory have been observed in the JUMP dataset^15^. In addition to these well-recognized technical effects observed in most data collection techniques, previous studies have found that CP features extracted using the conventional tool, CellProfiler^16^, exhibit distinctive well position effects^14^. Concretely, well position effects arise from the unique design of the CP experiment. In CP technology, experimental plates are organized into 16 rows and 24 columns, totaling 384 wells, with each well influenced by both row effects and column effects. We collectively refer to row effects, column effects, and the effects of different batches as triple effects. The complex and combined triple effects can lead to deviations from accurate biological profiles, and thus need to be corrected urgently.

However, no methods have been specifically designed to correct triple effects especially well position effects in CP profiles. Although there exist some batch correction methods designed for single-cell data, directly applying them to CP data remains challenging for several reasons. First, the characteristics of CP data differ significantly from single-cell data, as CP data is denser and exhibits lower variability compared to single-cell data^17^. Second, well position effects in CP data contrast with batch effects in single-cell data, as row or column effects show a gradient-influenced pattern, where greater differences in row or column numbers leads to more pronounced effects. Third, the triple effects, especially row and column effects, are complexly interactive and need to be corrected simultaneously. Some methods, such as scVI^18^, can correct only one type of technical effects and are constrained to correct multiple technical effects. Although methods like Harmony^19^ model one type of technical effects at a time and can correct all effects one-by-one, they are unable to simultaneously model triple effects in CP data.

Besides the challenges of correcting triple effects, existing studies that rely on features extracted by the CellProfiler tool still encounter several unresolved disputes and limitations^14^. First, it is currently unknown whether the well position effects are inherent to the CP data or arise from the feature extraction process using CellProfiler. Second, the CellProfiler software, as a non-end-to-end feature extraction tool, is overly reliant on traditional computer vision features and requires expert selection, which may overlook certain relevant phenotypic variation^4^. Third, the CellProfiler software is a non-data-driven feature extraction method, where the effectiveness of its predefined features is heavily reliant on the characteristics of the input images and domain knowledge.

Here, we demonstrated a one-stop method named cpDistiller for correcting triple effects and extracting latent patterns in CP data. cpDistiller mainly comprises three modules: the extractor module for deriving more comprehensive image information, the joint training module for integrating dual-source features, and the technical correction module for simultaneously correcting batch, row, and column effects. Specifically, the extractor module, inspired by transfer learning, employs a pre-trained segmentation model in an end-to-end and data-driven manner, which is adjusted to extract features from nearly 30 terabytes of raw images. The joint training module aligns the features extracted by both CellProfiler and the extractor module, improving the model’s ability to better characterize cell-to-cell variation. The technical correction module employs a semi-supervised Gaussian mixture variational autoencoder (GMVAE), incorporating contrastive and domain-adversarial learning strategies, to simultaneously correct technical effects. Based on comprehensive experiments across various CP profiles, we demonstrated that cpDistiller exceled in both qualitative visualizations and quantitative metrics, outperforming five baseline methods in single-batch well position effect correction as well as simultaneous triple-effect correction, all while preserving biological heterogeneity. Besides, we showcased the extensive capabilities of cpDistiller, including the ability to integrate more information-rich image features, the support for incremental learning, and the robustness to various feature selection strategies. Moreover, we emphasized that cpDistiller effectively captures system-level phenotypic responses to genetic and chemical perturbations, serving as a powerful tool to complement single-cell RNA sequencing (scRNA-seq) data for uncovering gene functions and relationships. In addition to combination with scRNA-seq, cpDistiller has the potential to provide unbiased insights into gene associations independently. Furthermore, by improving the matching of genetic perturbations with their target genes and enhancing gene-compound similarity assessments, cpDistiller shows strong promise for accelerating the identification of targets, which is quite valuable in facilitating drug discovery and various fields of biological research.

## Results

### The overall architecture of cpDistiller

cpDistiller maps the input data to the low-dimensional embedding space that aims to correct triple effects while capturing true biological signals. Specifically, cpDistiller is composed of three main modules: the extractor module, the joint training module and the technical correction module (Fig. 1).

**Fig. 1.**
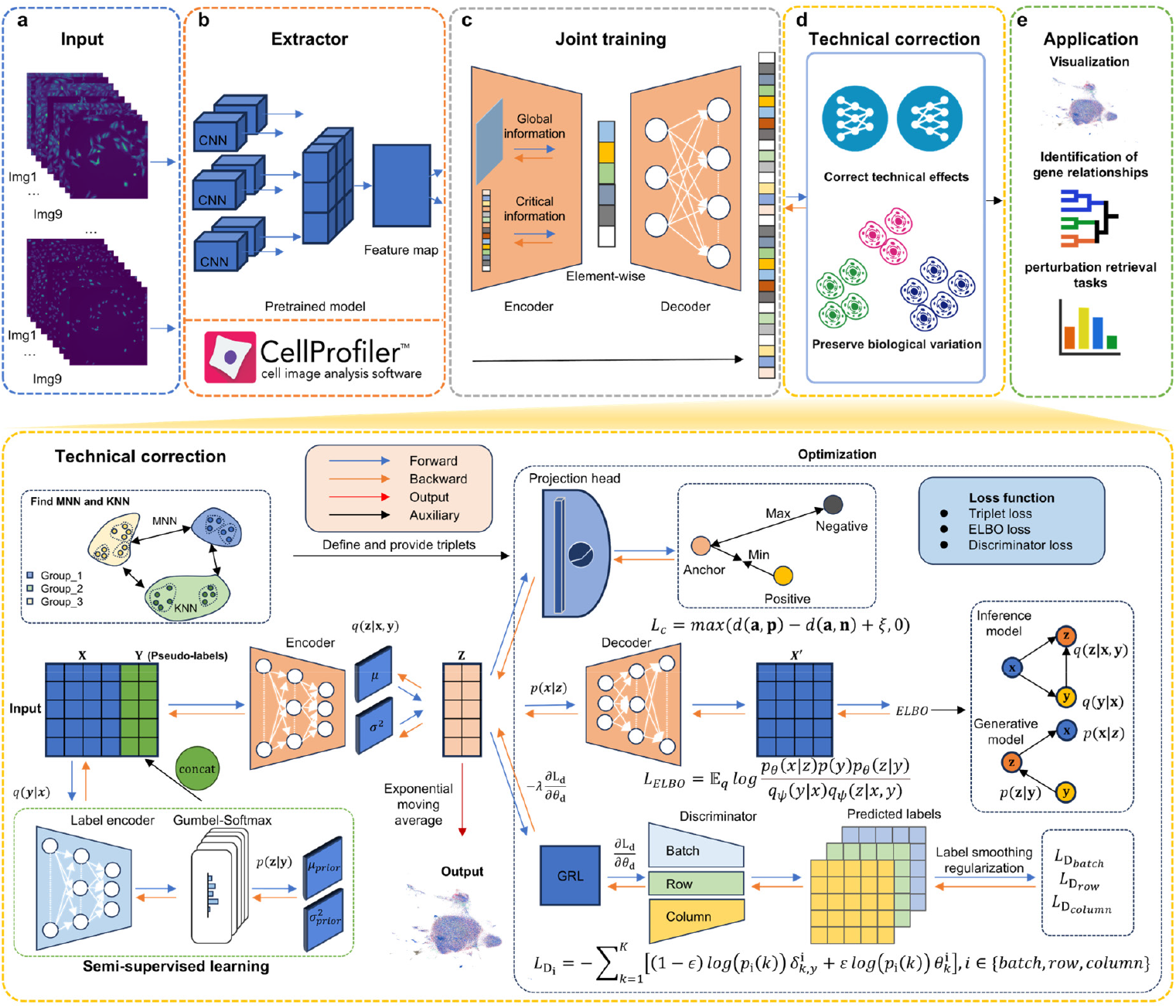
Overview of cpDistiller. **a**,**b**, cpDistiller takes CP images as input (**a**) and extracts features by CellProfiler and the extractor module (**b**). **c**, The joint training module integrates features from both cpDistiller-extractor module and CellProfiler. The CellProfiler-based features retain unprocessed while the cpDistiller-extractor-based features undergo the encoder-decoder architecture with an attention mechanism. These refined features are then input into the technical correction module. **d**, The technical correction module, built on a GMVAE, infers pseudo-labels and obtains low-dimensional representations. In the low-dimensional space derived from the GMVAE, representations pass through a projection head before applying the triplet loss to restore more accurate nearest-neighbor relationships. Additionally, three discriminators are designed to predict batch, row, and column labels, and the technical correction module use the GRL to enable adversarial learning to correct triple effects. Moreover, the exponential moving average (EMA) is applied during training, and the averaged parameters are used as the final model parameters. **e**, The low-dimensional embeddings obtained by cpDistiller facilitate a range of applications, including data visualization, identification of gene relationships, and various perturbation retrieval tasks.

cpDistiller processes CP images at a resolution of 1080 × 1080 pixels and extracts comprehensive features from each well. These features form a matrix, where rows represent samples (cells, wells or perturbations) and columns correspond to extracted features. We first extract features from the raw images using CellProfiler, a widely-used but not data-driven approach, and refer to these features as CellProfiler-based features. Drawing inspiration from transfer learning, we further develop the extractor module based on an end-to-end pre-trained segmentation model to automatically extract features, and refer to these features as cpDistiller-extractor-based features (Methods).

Then, the two sets of features are introduced into the joint training module for integration. The CellProfiler-based features retain unprocessed, while the cpDistiller-extractor-based features are transformed through an attention mechanism-based encoder, reducing them to a latent space, and then reconstructed via a decoder. This encoder-decoder structure, applied exclusively to the cpDistiller-extractor-based features, ensures feature refinement by reducing noise and enhancing the quality of the representations. The attention mechanism emphasizes key features across dimensions, facilitating a more effective combination with the CellProfiler-based features while filtering out irrelevant information (Methods).

The technical correction module, with a GMVAE at its core, infers pseudo-labels for each well in a semi-supervised mode using the features integrated by the joint training module. These pseudo-labels represent the Gaussian components that characterize the underlying patterns of each well, capturing the differences among these patterns. By combining these pseudo-labels along with the integrated features, the technical correction module derives the latent low-dimensional representations for each well. Besides, we employ both contrastive and domain-adversarial learning strategies. In contrastive learning, we use *k*-nearest neighbors (KNN) and mutual nearest neighbors (MNN) to construct triplets, applying triplet loss^20^ to restore more accurate nearest-neighbor relationships for each well. To facilitate domain-adversarial learning and avoid stage-wise training in generative adversarial networks (GANs)^21^, we further apply the gradient reversal layer (GRL)^22^ in an end-to-end training approach, making it harder for discriminators to distinguish data sources and thus removing technical effects. Additionally, for calculating the discriminator loss, we design a soft label distribution tailored to the gradient-influenced pattern of the CP data, where each entry in the distribution vector reflects its proximity to the actual data source for a more accurate representation.

Currently, no specialized methods are available for correcting triple effects, especially well position effects in CP data. However, a recent benchmark study has explored the application of single-cell batch correction methods as a potential solution^15^. In light of this, we compared cpDistiller with methods that excel at removing batch effects in single-cell data, including Seurat v5^23^, Harmony^19^, Scanorama^24^, scVI^18^ and scDML^25^. Among them, Seurat v5, Harmony, and Scanorama can iteratively correct triple effects by sequentially incorporating different technical labels (removal of batch, row, and column effects one after another), as they are based on low-dimensional representations, which allow for flexible, stepwise corrections. In contrast, scVI and scDML, directly model the original high-dimensional inputs to obtain low-dimensional representations, making them less suitable for iterative correction and limiting them to address only one type of technical effects (one of batch, row, or column effects). Due to the lack of metrics for evaluating the correction of technical effects in CP profiles, we use single-cell analysis metrics to assess the capacity of different methods to remove technical effects while preserving biological variation. Metrics such as average silhouette width (ASW)^26^, technic average silhouette width (tASW)^27^ and graph connectivity^27^ are used to measure the model’s ability to remove technical effects, while isolated label silhouette^27^, isolated label F1^27^, perturbation average silhouette width (pASW)^27^, and normalized mutual information (NMI)^27^ are used to evaluate the characterization of heterogeneity (Methods).

### The features derived from CellProfiler and cpDistiller-extractor module confirm triple effects

Previous studies have identified three types of technical effects in CP data: batch, row, and column effects^28^. Notably, row and column effects suggest the novel well position effects. Following the prior study^14^, we employed CellProfiler to obtain 1,446-dimensional features (CellProfiler-based features) from open reading frame (ORF) overexpression dataset in cpg0016, and then performed uniform manifold approximation and projection (UMAP)^29^ for these features. From the UMAP visualization, we observed distinct clustering patterns corresponding to different batches, rows, and columns (Fig. 2a). Batch effects are evident as patterns from different batches cluster into separate groups, while row and column effects are displayed as color gradients in the visualization, highlighting the presence of triple effects (Fig. 2a). To investigate whether row and column effects are evident within a single batch, we further observed the CellProfiler-based features from Batch_7. The UMAP visualizations revealed color gradients, indicating a gradient-influenced pattern where more distant rows or columns exhibit more pronounced effects, a phenomenon consistently observed across 12 batches (Fig. 2a and Supplementary Figs. 1-3).

**Fig. 2.**
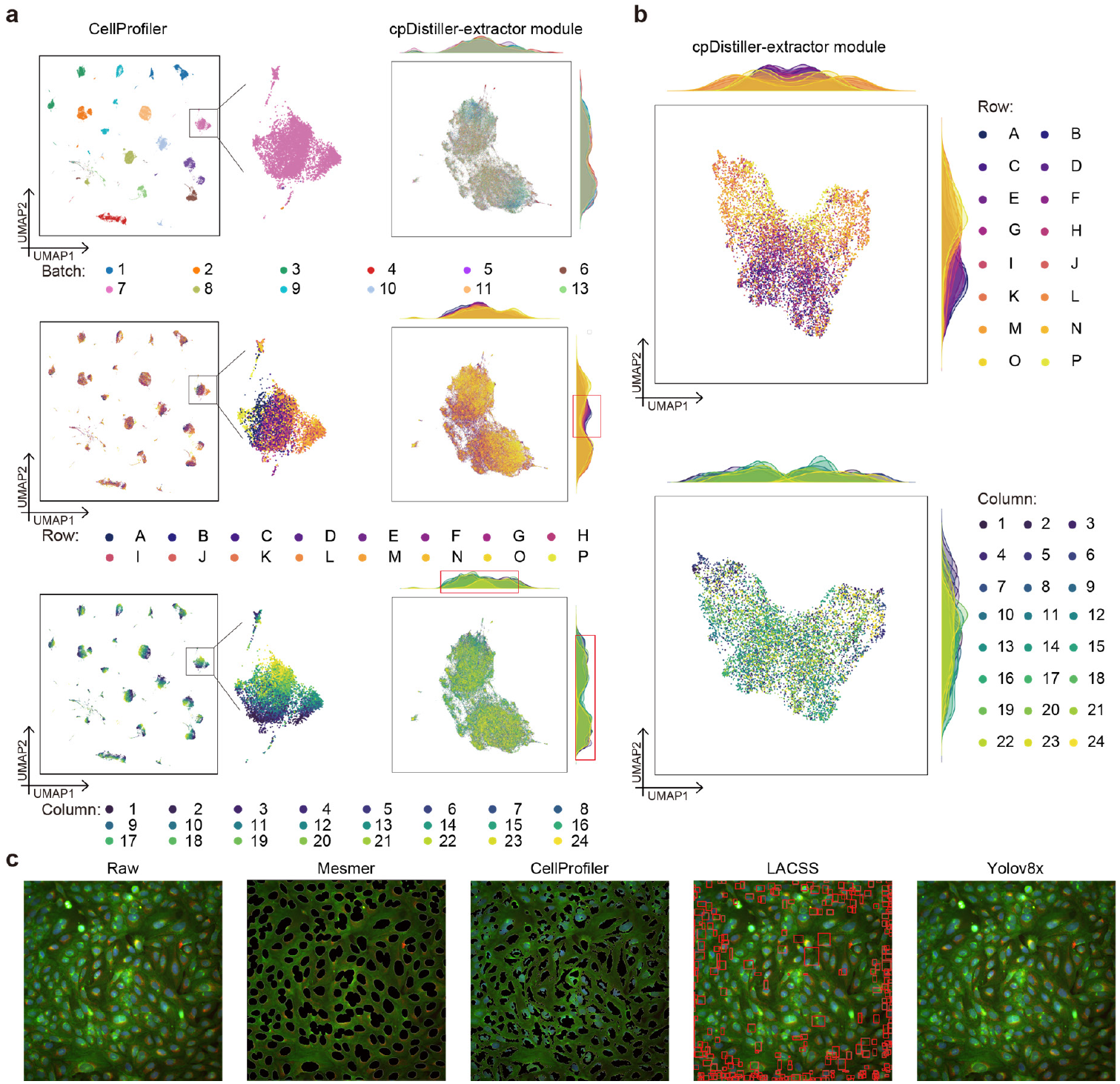
CellProfiler and cpDistiller-extractor module confirm triple effects. **a**, The UMAP visualizations of embeddings obtained by the CellProfiler (left) and cpDistiller-extractor module (right), colored by batch, row, and column, respectively. **b**, The UMAP visualizations of embeddings obtained by cpDistiller-extractor module in Batch_7, colored by row and column, respectively. **c**, Images show segmentation or detection results of different models. The raw image is derived from one of the nine sub-images in a well from CP images. Segmentation results are shown as black areas for Mesmer and CellProfiler, while detection results in the LACSS are highlighted with red boxes. YOLOv8 fail to perform cell segmentation from CP images.

To evaluate whether the triple effects are due to inherent characteristics of CP data or biases in the feature extraction process with CellProfiler, we developed the cpDistiller-extractor module, based on the pre-trained segmentation model Mesmer^30^, to extract deep-learning features (cpDistiller-extractor-based features). As illustrated in Fig. 2a, UMAP visualizations of the cpDistiller-extractor-based features from 12 batches display slight batch effects and clear, consistent effects related to row and column (Fig. 2a). We further plotted and observed the density distributions of the scatter points along the **x** and **y** axes in the UMAP visualizations. These distributions revealed distinct peaks corresponding to different batches, indicating the presence of batch effects (Fig. 2a). Additionally, we observed that the distributions for rows and columns were non-overlapping, with row-specific and column-specific peaks highlighted in red boxes, suggesting the existence of row and column effects (Fig. 2a). Furthermore, UMAP visualizations of the cpDistiller-extractor-based features from Batch_7 demonstrated clear evidence of row and column effects (Fig. 2b), a pattern that was consistently observed across all batches (Supplementary Figs. 4 and 5).

To investigate whether the cpDistiller-extractor module can capture more precise information regarding cell positions and shapes from CP images, we compared its segmentation results with those obtained from CellProfiler. As illustrated in Fig. 2c, the Mesmer model, which serves as the core of cpDistiller-extractor module, outperformed CellProfiler in segmentation capabilities, particularly in capturing details of cell morphology (Supplementary Text 1 and Supplementary Fig. 6). Before adopting Mesmer, we also assessed the widely used LACSS^31^ and YOLOv8^32^ models as alternatives. However, neither was able to accurately locate cell positions (Fig. 2c and Supplementary Fig. 7). We further tested the YOLOv8 model with various parameters, but the results confirmed that it remained ineffective for cell detection and segmentation in CP data (Supplementary Figs. 8-13). Details regarding the pre-trained model settings can be found in Supplementary Text 2.

In conclusion, features from both CellProfiler and cpDistiller-extractor module captured information about cell positions and shapes in CP images and revealed the inherent batch, row and column effects of CP data.

### cpDistiller effectively corrects well position effects and preserves cellular phenotypic heterogeneity

We first conducted experiments on the cpg0016 ORF profiles derived from the U2OS cell type in the JUMP dataset to demonstrate the effectiveness of cpDistiller. The cpg0016 ORF profiles consisted of 12 batches, and we used Batch_1 as an example. In the raw data, UMAP visualization reveals a gradient-influenced pattern, with more distant rows or columns exhibiting more pronounced effects (Fig. 3a,b and results from other methods and batches in Supplementary Figs. 14-25). We then visualized the low-dimensional embeddings obtained by cpDistiller and observed that the gradient-influenced pattern was significantly reduced. The data distribution across different rows was more uniform, and the previously pronounced column effects, particularly the large differences between low-numbered columns (e.g., Column_1) and high-numbered columns (e.g., Column_24), were well-mixed and greatly corrected (Fig. 3a,b). This demonstrated that cpDistiller can effectively correct both row and column effects. Moreover, overexpression reagents and compounds are theoretically expected to induce significant differences in cell phenotypes between negative and positive controls. Specifically, by utilizing UMAP to visualize the impact of various perturbations, cpDistiller successfully differentiated positive controls, including JCP2022_037716, JCP2022_035095, and JCP2022_012818 from negative controls across various batches (Supplementary Figs. 14-25). In addition to analyzing the controls, we also explored the ORF perturbations in treatment. cpDistiller successfully identified perturbations caused by 12 overexpression reagents in treatment and preserved their unique stimulatory effects, which were obscured in the raw data due to well position effects (Fig. 3c). In contrast, UMAP visualizations and hierarchical clustering results suggest that methods like scDML and scVI, which are limited to correcting only one type of technical effects, struggle to preserve biological variation while also failing to effectively correct well position effects (Supplementary Figs. 14-25 and Supplementary Fig. 26a). Although these methods may achieve partial mixing across rows and columns, they still fail to remove most non-biological noise, such as the clear striping pattern, particularly noticeable in Batch_5 and Batch_8 (Supplementary Figs. 18 and 21). While methods like Harmony, Seurat v5, and Scanorama can iteratively correct well position effects, they also fall short in preserving the true biological variation of ORF perturbations (Supplementary Fig. 26a).

**Fig. 3.**
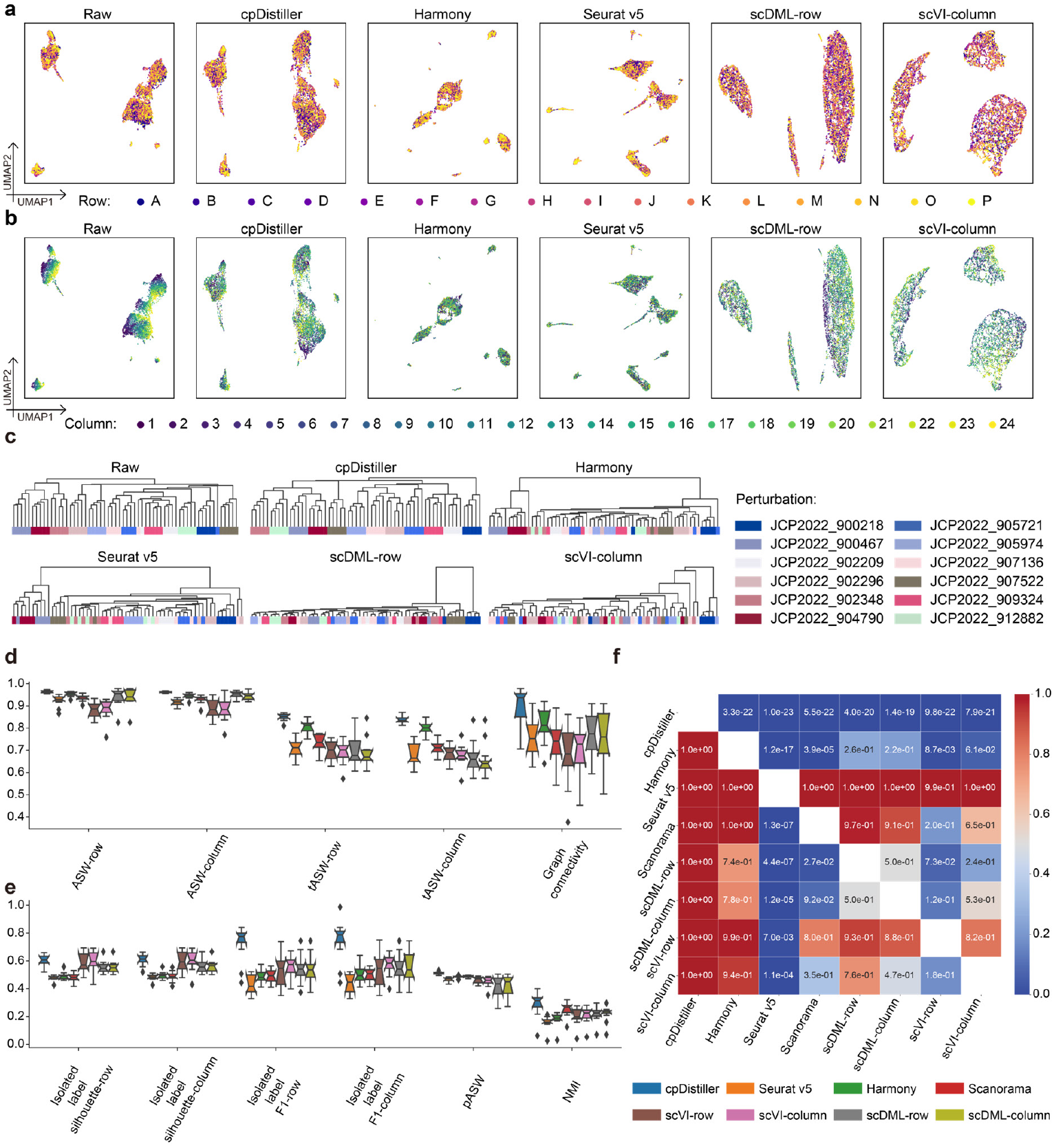
cpDistiller can effectively correct well position effects while preserving biological variation for ORF profiles across 12 batches. **a**,**b**, UMAP visualizations of embeddings obtained by different methods in Batch_1 colored by row (**a**) and column (**b**). **c**, Dendrograms illustrate the clustering of ORF perturbations induced by 12 reagents in treatment, graphically rendered based on low-dimensional representations generated by different methods. **d**,**e**, Quantitative evaluation of the performance of different methods on ORF profiles across 12 batches for correcting well position effects while preserving biological variation. The top plot focuses on technical correction capabilities (**d**), while the bottom plot highlights biological preservation abilities (**e**). Since the well position effects encompass both row and column effects, the metrics of ASW, tASW, isolated label scores yield two results, respectively. In the boxplots, center lines indicate the medians, box limits show upper and lower quartiles, whiskers represent the 1.5× interquartile range, and notches reflect 95% confidence intervals via Gaussian-based asymptotic approximation. **f**, Heatmap shows *P*-values from one-sided paired Wilcoxon signed-rank tests. Each cell in the heatmap reflects the statistical significance of one method’s (row) superiority over another (column), derived from 132 evaluations across 12 batches using 11 metrics.

To quantitatively demonstrate the advantages of cpDistiller in correcting well position effects, we further conducted experiments in 12 batches and assessed the performance by ASW, tASW, and graph connectivity as suggested in refs.^15,26,27^. ASW and tASW were used to assess the extent of data mixing across different technical labels. Graph connectivity assumed that, once technical effects were corrected, data with the same biological labels, specifically perturbation labels, should cluster more tightly. cpDistiller outperformed other baseline methods in all metrics and consistently demonstrated stable performance across 12 batches (Fig. 3d), indicating superior performance in both uniformly mixing CP profiles and accurately reflecting cell phenotype differences induced by various perturbations.

We further assessed the preservation of biological variation via metrics such as isolated label scores (including isolated label F1 and isolated label silhouette), pASW, and NMI. Isolated labels refer to perturbation labels with the least number of the technical effects. Here, the perturbation labels computed from isolated labels all belonged to positive controls. Higher isolated label scores reflect a greater impact of positive controls on cell phenotypes compared to others. cpDistiller excelled in isolated label scores, particularly achieving the highest isolated label F1 score among all baseline methods (Fig. 3e). Additionally, for the clustering metrics of pASW and NMI, which were used to assess the clustering quality of replicated experiments involving the same types of perturbations as suggested in refs.^15,27^, cpDistiller also surpassed other methods (Fig. 3e). Harmony has demonstrated excellent performance in removing batch effects for CP data in recent benchmark analysis^15^ and also emerges as the most effective baseline method for correcting row and column effects in our study. To further demonstrate the significant advantages of cpDistiller, we used one-sided paired Wilcoxon signed-rank tests to compare the performance of different methods across 12 batches. The *P*-values highlighted cpDistiller’s significant advantages in removing well position effects and preserving biological variation across batches, outperforming all baseline methods (Fig. 3f).

Besides, in our previous experiments with baseline methods like Seurat v5, Harmony, and Scanorama, which can iteratively remove well position effects, we initially corrected rows before columns. To investigate whether the performance of these methods was influenced by the order of correction, we also applied the reverse order, namely correcting columns before rows. The box plots across 12 batches showed no significant difference in performance when the same method was applied with different correction orders (Supplementary Fig. 27a). However, when we used one-sided paired Wilcoxon signed-rank tests for a more detailed quantification of performance differences, we observed that, although the differences were not statistically significant, the *P*-values indicated a trend where correcting rows before columns generally yielded better results than the reverse order for all baseline methods (Supplementary Fig. 27b).

In summary, cpDistiller demonstrated superior performance on 12 batches of ORF data spanning diverse CP profiles, achieving a satisfactory balance between correcting well position effects and preserving cellular phenotypic heterogeneity.

### cpDistiller enables effective and simultaneous correction of triple effects

After integrating ORF profiles from 12 batches and visualizing them using UMAP, we observed batch effects in addition to well position effects (Fig. 4a). To study biological-process-related mechanisms of action based on feature similarity^33^, it is necessary to remove batch effects to ensure reliable comparisons across batches. For competently removing batch effects, in addition to the discriminators for row and column labels, we also created an additional discriminator to identify batch labels, encouraging cpDistiller to learn low-dimensional representations that are indistinguishable with respect to the sources of batch labels, thereby removing batch effects (Methods). At the same time, we additionally considered KNN intra batches and MNN inter batches to construct triplets, using the contrastive learning technique to restore more accurate nearest-neighbor relationships for each well (Methods). We next verified that cpDistiller can satisfactorily correct triple effects.

**Fig. 4.**
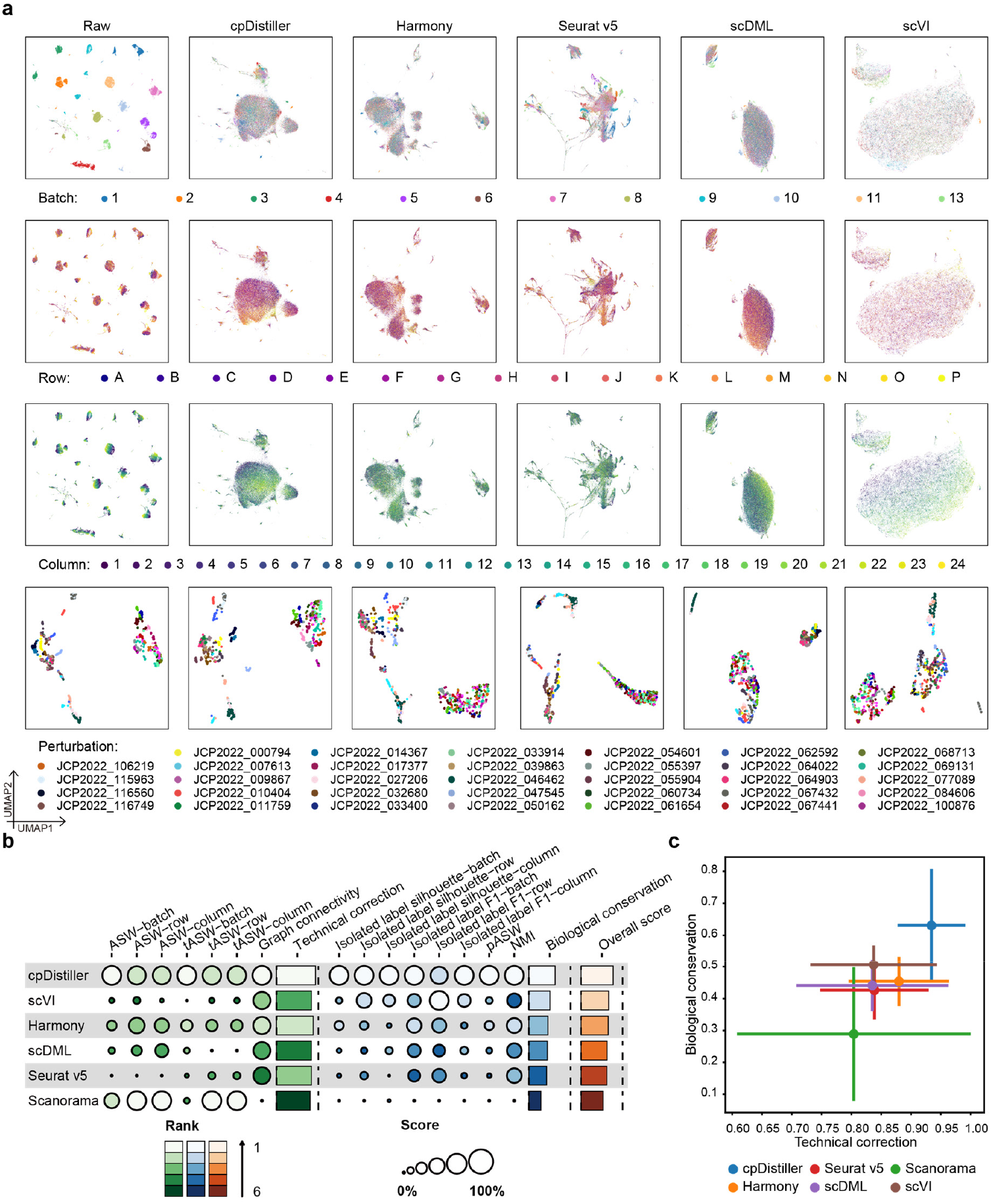
cpDistiller can simultaneously correct triple effects while preserving cellular phenotypic heterogeneity. **a**, UMAP visualizations of embeddings obtained by different methods across 12 batches in ORF profiles, colored by batch, row, column, and perturbation, respectively. The embeddings are consistent with all visualizations, and the selected 34 perturbations are shown. **b**, Overview of benchmarking outcomes for different methods. Technical correction and biological conservation scores refer to the average performance in these two aspects, whereas the overall score represents the aggregate performance across all metrics for different methods. Due to triple effects, which encompass batch, row and column effects, the metrics of ASW, tASW, isolated label scores yield three results, respectively. **c**, Scatter plot summarizes overall performance on triple-effect correction in ORF profiles. The x-axis represents the technical correction score, while the y-axis shows the biological conservation score. Each point corresponds to the average value per method, with error bars indicating the standard deviation.

First, we qualitatively compared the performance of different methods using UMAP visualizations, which showed that cpDistiller achieved successful mixing of data across batches, rows, and columns simultaneously (Fig. 4a and Supplementary Fig. 28). Moreover, to qualitatively assess how well different methods preserve the specificity of perturbations, compound perturbations shared across 12 batches on the target plates provided a reliable measure^15^. UMAP visualizations showed that cpDistiller ensured perturbations caused by the same compounds had a tendency to be clustered (Fig. 4a). To be specific, cpDistiller successfully identified multiple perturbations in target plates, including JCP2022_010404, JCP2022_000794, and JCP2022_047545. However, methods that are limited to correcting only one type of triple effects, such as scDML and scVI, failed to correct well position effects, resulting in the persistence of striping pattern for rows and columns (Fig. 4a). On the other hand, methods capable of iteratively correcting triple effects, including Scanorama and Seurat v5, tended to overcorrect, leading to fail to preserve true biological signals (Supplementary Fig. 28). To further evaluate if cpDistiller effectively maintained biological variation during triple-effect correction, we reanalyzed the ORF perturbations that were assessed during the correction of well position effects. As shown in Supplementary Fig. 26b, cpDistiller can still identify these differential perturbations, whereas other methods cannot.

Moreover, we quantitatively evaluated the performance of different methods. cpDistiller generally exhibited superior performance in correcting triple effects compared to baseline methods (Fig. 4b). Although Scanorama performed better than cpDistiller in tASW, it scored lowest in graph connectivity. It indicated that while the data was well mixed across batches, rows, and columns, it lost the specificity of the perturbations. Additionally, when assessing the preservation of biological variation through clustering metrics, cpDistiller consistently performed best across all metrics (Fig. 4b). Besides, we noted the balance between technical effect removal and biological conservation. We found that all baseline methods struggled to achieve the equilibrium (Fig. 4c). Specifically, scVI and Harmony both performed comparably well among the baseline methods. However, scVI tended to favor the preservation of biological variation, while Harmony focused more on technical correction (Fig. 4c). In contrast, cpDistiller balanced both aspects, offering superior overall performance in both removing triple effects and preserving cellular phenotypic heterogeneity (Fig. 4c).

Furthermore, to comprehensively evaluate the advantages of cpDistiller, we conducted a series of experiments to demonstrate its ability to leverage more information-rich image features for enhanced performance, support incremental learning for ongoing studies, and maintain robustness to feature selection. To investigate the impact of cpDistiller-extractor-based features on the performance of cpDistiller, we conducted experiments using only CellProfiler-based features. We found that cpDistiller still outperformed baseline methods in correcting technical effects while preserving biological variation, even when utilizing only CellProfiler-based features, as the baseline methods did (Supplementary Text 3 and Supplementary Fig. 29a-c). Moreover, with leveraging cpDistiller-extractor-based features, cpDistiller can exploit a broader range of image characteristics from raw images and lead to further performance improvements (Supplementary Text 4 and Supplementary Fig. 30).

Besides, cpDistiller supports incremental learning (Supplementary Text 5). Methods like Harmony and Scanorama require reprocessing and realigning the entire dataset to integrate new data, which can be cumbersome, especially when working with large public datasets^15^. These processes often involve modifying existing representations, which can disrupt the continuity of prior analytical results. In contrast, cpDistiller can leverage the model parameters learned from previous tasks, enabling the direct integration of new data without the necessity of realignment (Supplementary Fig. 31a). These flexibilities are particularly beneficial for ongoing studies, enabling seamless integration while preserving the integrity of previous analyses.

Additionally, cpDistiller demonstrates robustness to feature selection (Supplementary Text 6). We first extracted features with dimensions of 4,752 from fluorescence images and 7,638 when combined with brightfield images by using the CellProfiler software (Methods). We then utilized these two types of features to carry out two key tasks: correcting well position effects in single batch and simultaneously correcting triple effects. The results showed that even when dealing with high-dimensional features that may contain substantial redundant features and noise, cpDistiller consistently outperformed baseline methods in correcting technical effects and preserving biological variation (Supplementary Fig. 31b,c).

Overall, cpDistiller achieved a satisfactory balance between correcting triple effects and preserving biological variation, and demonstrated strong capabilities in information-rich feature extraction, incremental learning, and robust feature selection.

### cpDistiller can combine scRNA-seq data to reveal gene functions and relationships

Inferring gene functions and interactions, as a fundamental step in many areas of biological research, often relies on various types of sequencing data such as scRNA-seq data and chromatin immunoprecipitation sequencing (ChIP-seq) data^34^. This inferring task is associated with numerous unknown and highly intricate biological processes, but the sequencing methods tend to focus on only a limited number of interesting molecular-level changes, making obtaining thorough and accurate inferences challenging^35^. In contrast, we demonstrated that cpDistiller offers comprehensive system-level phenotypic characteristics under genetic perturbations and has the potential to integrate molecular-level RNA data for revealing gene functions and relationships.

To demonstrate the capability of the embeddings learned by cpDistiller in preserving system-level phenotypic characteristics of perturbation heterogeneity, we applied cpDistiller to the controls in the ORF data from the cpg0016 dataset. Specifically, we focus on controls, including the positive controls JCP2022_037716, JCP2022_012818, and JCP2022_035095, and the negative control JCP2022_915131. The positive controls are expected to produce noticeable phenotypic changes, while the negative control should induce minimal changes^14^. Theoretically, phenotypes under identical genetic perturbation in different wells or different plates should yield consistent patterns. Therefore, we evaluated the similarity of identical positive and negative controls duplicated across different wells and plates within a batch, using embeddings generated by cpDistiller and baseline methods, respectively. Taking Batch_4 for example, after performing hierarchical clustering based on cosine similarity (Fig. 5a), we found that the baseline methods failed to cluster identical treatment sets. In contrast, cpDistiller successfully grouped duplicated positive and negative controls from different wells, as well as revealing treatment-specific embeddings. These results illustrated that cpDistiller can effectively capture phenotypic signals across various genetic perturbations.

**Fig. 5.**
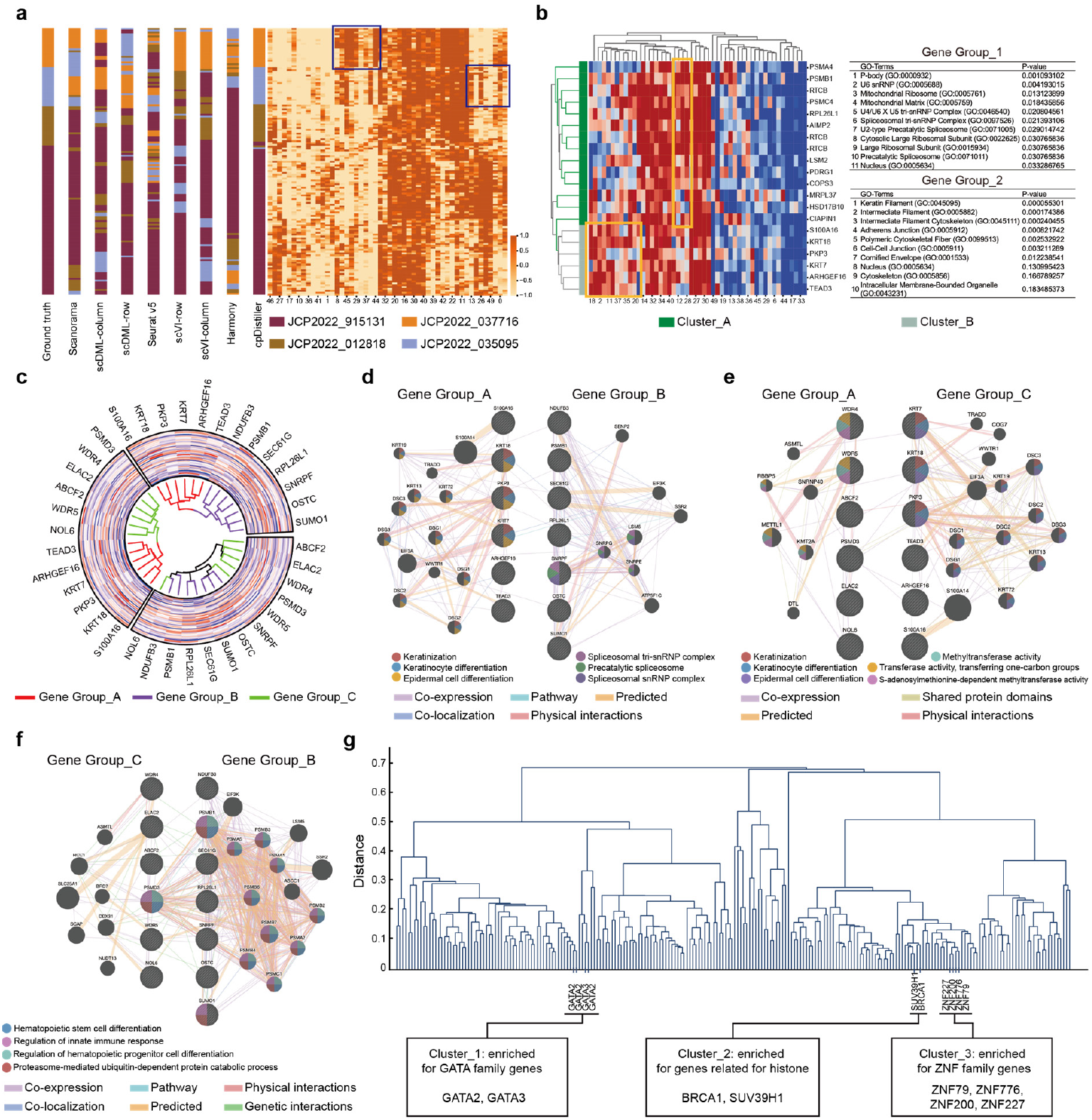
Analyses of gene functions and gene relationships. **a**, Dendrograms illustrate the clustering of duplicates of controls, graphically rendered based on low-dimensional representations generated by different methods. The additional heatmap visualizes the 50-dimensional embeddings obtained by cpDistiller, highlighting the clustering patterns of controls. **b**, Hierarchical clustering and heatmap of the 50-dimensional embeddings obtained by cpDistiller for gene Group_1 and gene Group_2. Detailed views show the cellular compound analysis of GO term analysis for gene Group_1 and gene Group_2. **c**, Circular dendrograms are performed for genes in different gen groups, graphically rendered based on embedding obtained by cpDistiller. **d-f**, The networks of gene relationships are provided by the GeneMANIA tool for gene Group_A and gene Group_B (**d**), gene Group_A and gene Group_C (**e**), and gene Group_B and gene Group_C (**f**). Each dot represents a gene, with the dot’s color indicating its related enrichment functions from GeneMANIA. **g**, Dendrogram of hierarchical clustering for genes using the embeddings learned by cpDistiller.

To elucidate cpDistiller’s capability to integrate with scRNA-seq data for inferring gene functions, we used the embeddings obtained by cpDistiller to analyze genetic perturbation data from the ORF dataset. Given that the ORF dataset encompasses large-scale perturbations targeting 12,602 genes, we first employed the ARCHS4 tool^36^ to establish gene groups based on the similarities of their scRNA-seq data, where genes with highly similar expression patterns were grouped together (Methods). Next, we calculated the Euclidean distance between the embeddings from cpDistiller for the individual genes in gene groups and then performed hierarchical clustering based on these distances. For example, we found gene Group_1 and gene Group_2 are highly separated into distinct clusters (Fig. 5b). Gene Group_1 is enriched in Cluster_A, while gene Group_2 is enriched in Cluster_B, indicating that cpDistiller has learned group-specific embeddings, resulting in significantly distinct morphological embeddings for genes in Group_1 and Group_2. To elucidate that the morphological embeddings from cpDistiller have the capacity to reveal gene functions, we conducted cellular component (CC) analyses via gene ontology (GO) enrichment for the genes in Group_1 and Group_2, respectively (Methods). We found that the top three significantly enriched cellular components for the genes in Group_1 were all linked with the function of RNA processing, including ‘P-bodies’, ‘U6 snRNP’, and ‘Mitochondrial Ribosome’^37–39^ (Fig. 5b). By contrast, the top three significantly enriched cellular components for the genes in Group_2 were all involved in the function of forming structural elements, including ‘keratin filament’, ‘intermediate filament’, and ‘intermediate filament cytoskeleton’^40^. The CC analysis results revealed that the genes in Group_1 and the genes in Group_2 are associated with distinct functions, which is consistent with the cluster categorization where genes in Group_1 and Group_2 have distinct embeddings learned by cpDistiller.

To demonstrate cpDistiller’s effectiveness in elucidating gene relationships when combined with scRNA-seq data, we further analyzed the embeddings obtained by cpDistiller for genes in gene groups through multiple types of biological analyses. We conducted hierarchical clustering analysis for individual genes in gene groups using embeddings obtained by cpDistiller and found that some gene groups formed distinct clusters. For example, as illustrated in Fig. 5c, gene Group_A and Group_B, as well as gene Group_A and Group_C, are distinguishable into different clusters. However, gene Groups_B and Group_C are not easily separated. These results indicated that although genes within different groups have dissimilar scRNA-seq expression patterns, they may have indistinguishable phenotypic embeddings obtained from cpDistiller. To elucidate that the embeddings from cpDistiller can illustrate gene relationships, we utilized the GeneMANIA tool^41^ to construct gene networks to visualize gene relationships based on large and diverse databases^41^ (Methods). We found that the correlation in the network for genes in Group_A and Group_B is minimal (Fig. 5d), as well as the correlation for genes in Group_A and Group_C (Fig. 5e). By contrast, genes in Group_B and Group_C exhibit closer relationships (Fig. 5f). These findings aligned with the clustering results from cpDistiller, where genes in Group_A and Group_C, as well as those in Group_A and Group_B, have significantly different embeddings, while those in Group_B and Group_C share similar embeddings. These results indicate that genes with similar embeddings by cpDistiller tend to have closer relationships, illuminating that cpDistiller can uncover gene relationships. Moreover, we used GeneMANIA to predict the significantly enriched gene functions within different groups and found that functional enrichment results consistently aligned with the clustering results (Fig. 5d-f). These confirmed that analyzing the embeddings obtained by cpDistiller has the potential to infer gene functions. In conclusion, these results demonstrated the phenotypic embeddings obtained by cpDistiller effectively capture gene functions and relationships, highlighting its potential as a valuable tool for exploring gene interactions.

In addition to combination with scRNA-seq data, we further demonstrated that cpDistiller alone is also capable of uncovering gene relationships. We first applied cpDistiller to each batch in the ORF dataset, with each batch containing approximately 2,000 genetic perturbations. Since many genetic perturbations did not induce significant phenotypic changes, we then screened for active genes that exhibited substantial phenotypic changes compared to controls in each batch (Methods). We calculated the similarity of the embeddings from cpDistiller for these active genes, and then performed hierarchical clustering based on the similarity to identify significantly clustered genes (Methods). As illustrated in Fig. 5g, taking Batch_1 as an example, we found that *SUV39H1* and *BRCA1* are grouped together. Both *SUV39H1* and *BRCA1* exhibit co-expression as well as various connections^42^ and they are associated with condensed chromosomes, histone H3-K9 methylation, and related chromosomal modifications^43^. Additionally, genes from the zinc finger protein family which share protein domains^44,45^, including *ZNF227, ZNF200, ZNF776*, and *ZNF79*, are successfully clustered together based on the embeddings obtained by cpDistiller. Similarly, *GATA2* and *GATA3*, members of the *GATA* transcription factor family crucial for cell differentiation and development, which share protein domains^46^ and exhibit co-expression^44,45^, are also found to be clustered together. These results confirmed that analyzing the embeddings learned by cpDistiller alone also has the potential to reveal gene relationships.

### cpDistiller can facilitate the search for gene and compound targets

The JUMP dataset is primarily generated using U2OS cells, with all ORF experimental data in the cpg0016 dataset conducted on this cell type. However, since biological research often extends beyond U2OS cells, the JUMP dataset includes a pilot dataset, cpg0000, conducted on A549 cells in a single batch. The cpg0000 dataset provides paired genetic and compound perturbations targeting the same gene, offering substantial potential for uncovering biological targets^47^. Leveraging this design, researchers have established simulated tasks to retrieve gene and compound targets using features extracted by CellProfiler^47^. However, as shown in Fig. 6a,b, when we performed UMAP visualization using CellProfiler-based features for A549 cells, we observed noticeable row and column effects, which could obscure real biological signals. To address this, we applied cpDistiller to correct both row and column effects. As illustrated in Fig. 6a,b, the data distributions across different rows and columns were more uniform.

**Fig. 6.**
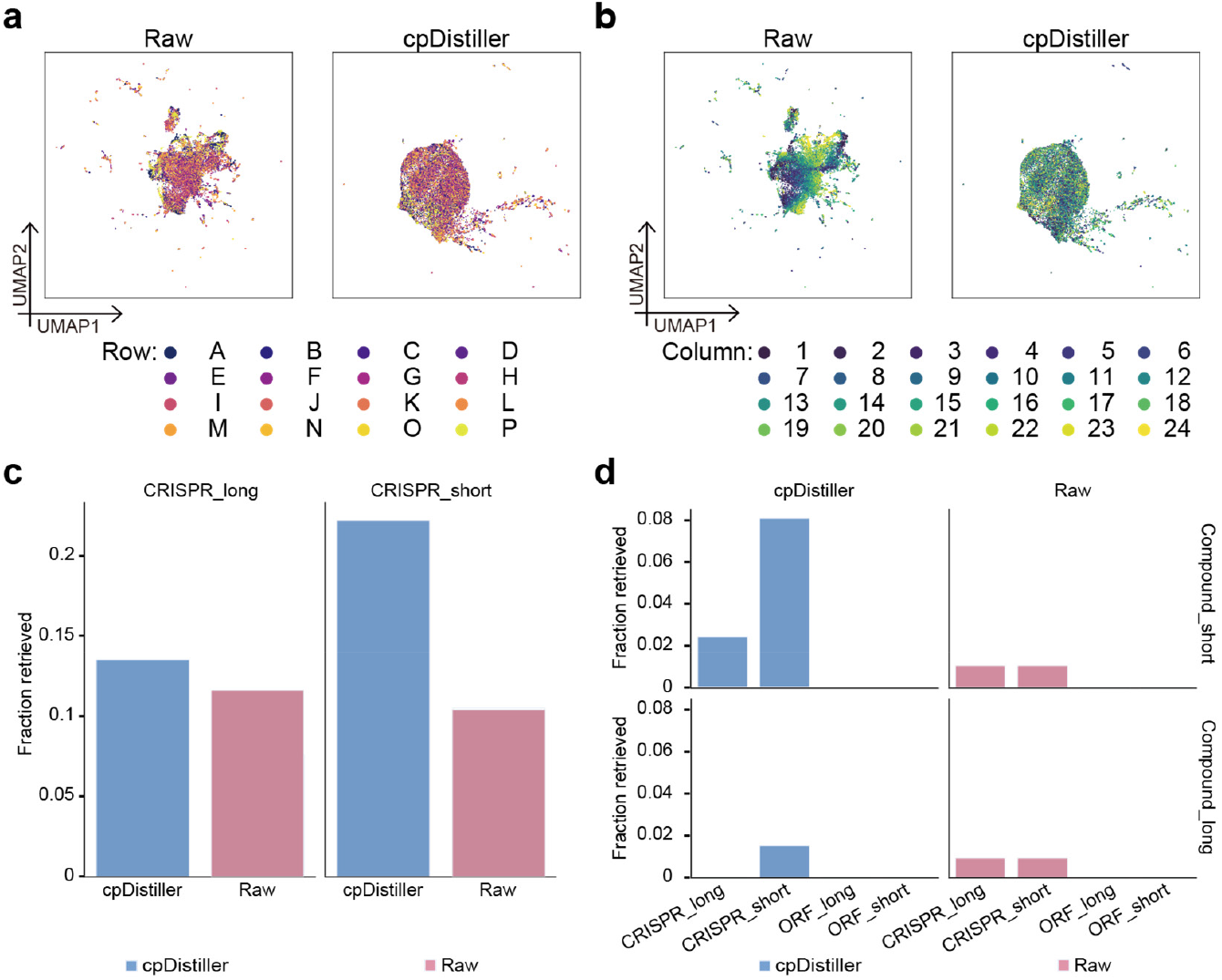
The performance of cpDistiller in retrieval tasks. **a**,**b**, UMAP visualization for the A549 cells using the features extracted by CellProfiler and embeddings learned by cpDistiller, colored by row (**a**) and column (**b**). **c**, Fraction retrieved scores for retrieving sister CRISPR guides in A549 cells using embeddings of cpDistiller and CellProfiler-based features, respectively. Short (_short) and long (_long) time points mean the experimental conditions. **d**, Fraction retrieved scores for retrieving gene-compound pairs in A549 cells using the embeddings learned by cpDistiller and CellProfiler-based features, respectively. Short (_short) and long (_long) time points mean the experimental conditions.

We further demonstrated that the cpDistiller-derived embeddings facilitate the identification of gene and compound targets. Specifically, the cpg0000 dataset includes genetic and compound perturbations for 160 genes in A549 cells, conducted at both long and short time points. Each gene is perturbed through one ORF treatment, two gene knockouts by Clustered Regularly Interspaced Short Palindromic Repeats (CRISPR) guides, and two compound experiments^47^. Using sister CRISPR guides targeting the same gene, researchers designed retrieval tasks to simulate gene target identification, calculating the fraction retrieved score to evaluate retrieval performance^47^. Following their workflows, we calculated the fraction retrieved scores using the embeddings from cpDistiller and compared the scores to those generated using CellProfiler-based features (Methods). As shown in Fig. 6c, using CellProfiler-based features, the fraction retrieved score for retrieving sister CRISPR guides at long time points (long-time CRISPR retrieval) is 0.123 and the score for retrieving sister CRISPR guides at short time points (short-time CRISPR retrieval) is 0.11. In contrast, with embeddings obtained by cpDistiller, the fraction retrieved scores improve to 0.139 for long-time CRISPR retrieval and 0.235 for short-time CRISPR retrieval. This represented a notable increase, particularly for short-time retrieval tasks, where cpDistiller achieves over a two-fold increase compared to CellProfiler-based features. These results highlighted the pivotal role of cpDistiller in enhancing the accuracy of gene retrieval tasks, making it a valuable tool for identifying genes involved in similar processes and uncovering critical gene relationships.

In addition to retrieving sister perturbations, researchers also conducted cross-modality gene-compound retrieval tasks to simulate the identification of compound targets^47^. Concretely, they searched for compounds that produced similar effects on cell morphology as the query gene using CellProfiler-based features. They calculated the fraction retrieved scores to evaluate retrieval performance. Following their methodology, we calculated the fraction retrieved scores using embeddings generated by cpDistiller to demonstrate that cpDistiller can improve retrieval results. As shown in Fig. 6d, when using CellProfiler-based features for compound perturbations, the fraction retrieved scores are nearly 0.008 for retrieving all compound-CRISPR pairs. In contrast, when using embeddings extracted by cpDistiller for compound perturbations at both long and short time points, the fraction retrieved scores for retrieving most compound-CRISPR pairs over a two-fold increase (Fig. 6d). Given the complexity and importance of gene-compound retrieval tasks, even slight improvements hold substantial value^47^. These results demonstrated that cpDistiller achieved notable improvements in gene-compound retrieval tasks compared to uncorrected CellProfiler-based features, highlighting its potential to advance the discovery and evaluation of novel chemical compounds.

In conclusion, cpDistiller satisfactorily corrected well position effects and obtained embeddings that offered clear advantages in exploring gene and compound targets, providing valuable insights for biological research and drug discovery.

## Discussion

We developed cpDistiller, a method specifically designed to correct triple effects, particularly well position effects in Cell Painting (CP) data. cpDistiller leverages raw CP images by integrating high-level features via pre-trained segmentation model with handcrafted features from traditional methods, followed by a semi-supervised GMVAE utilizing contrastive and domain-adversarial learning to correct triple effects. We first conducted systematic experiments to demonstrate that CP data inherently exhibits triple effects, including batch, row, and column effects. Through comprehensive experiments on multiple batches varying in cell types, plate designs, perturbation types, we then validated cpDistiller’s superior performance in correcting triple effects and preserving cellular phenotypic heterogeneity. Moreover, we highlighted the extensive advantages of cpDistiller, including its ability to integrate information-rich details from raw images, its support for incremental learning, and its consistent robustness across various feature selection strategies. Additionally, we conducted various downstream experiments to illustrate the application of cpDistiller in revealing gene functions and gene relationships, highlighting its ability to uncover gene associations both when combined with scRNA-seq data and independently. Scientists have found that target-based drug discovery can be limited in certain situations, and phenotypic drug discovery is sometimes more likely to succeed^33^. In recent years, research related to CP has expanded significantly, offering a better perspective on studying drug targets and cellular phenotypic heterogeneity due to its simpler procedures and lower costs. Therefore, we also explored the potential of cpDistiller in identifying gene and compound targets, which could enhance drug discovery efforts.

While cpDistiller demonstrated excellent performance, there are several areas that could be explored for future improvement. First, compared to CellProfiler-based features, which has specific and interpretable for each dimension, the low-dimensional representations learned from cpDistiller may lack interpretability. Enhancing the interpretability of deep learning models remains a challenge^33^. Second, we can utilize Segment Anything Model^48^ along with expert annotation and refinement to generate accurate annotations for a subset of CP images. These annotations will serve as a basis for supervised training of the cpDistiller-extractor module, allowing it to effectively capture more detailed cellular information. Third, to gain a more comprehensive understanding of cellular behavior, cpDistiller could be extended to integrate with multi-omics profiling, connecting morphological and molecular phenotypes at single-cell resolution, thereby providing a more detailed insight into genetic perturbations and their effects^49^.

## Methods

### Overview of cpDistiller

cpDistiller consists of three main modules: the extractor module, the joint training module and the technical correction module. We first used the cpDistiller-extractor module to extract features from Cell Painting (CP) images. Then we integrated cpDistiller-extractor-based and CellProfiler-based features via the joint training module and processed the combined features through the technical correction module to remove batch, row and column effects.

### The extractor module

We develop the extractor module to address the limitations of non-data-driven approaches, such as CellProfiler, while minimizing biases from human interference. By modeling our extraction task as the segmentation task in computer vision, we transform raw images into low-dimensional representations using an end-to-end process. We select Mesmer^30^, pre-trained on the TissueNet datasets^30^, as the base model for transfer learning due to its strong performance in cell segmentation tasks on cellular images^50^. By experimenting with various intermediate layers of Mesmer and considering factors such as overall effectiveness, number of parameters, and processing speed, we select the backbone of Mesmer and its pre-trained parameters.

In the CP assay, each well contains 9 sub-images sized at 1080 × 1080 pixels, captured across five channels: mitochondria (Mito), nucleus (DNA), nucleoli and cytoplasmic RNA (RNA), endoplasmic reticulum (ER), and Golgi and plasma membrane and the actin cytoskeleton (AGP)^14^. To meet Mesmer’s dual-channel input requirements and enhance the visibility of cell contours, we discard the AGP and Mito channels, focusing instead on the DNA channel and the cell channel created by averaging the ER and RNA channels, as the information in the AGP and Mito channels is difficult to predict^33^. Besides, to align with the Mesmer’s specifications for pixel density and reduce computational resources, we apply a tiling operation, adjusting the stride and overlap ratio to crop each large sub-image into multiple small images with dimensions of 256 × 256 pixels. The number of small images generated from each well’s sub-image is calculated using the following formula:

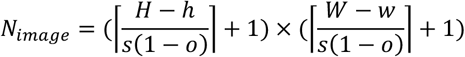

where *H* and *W* represent the height and width of each large sub-image, while *h* and *w* denote the height and width of the cropped images. *s* denotes the sliding stride, which refers to the number of pixels moved per step, while *o* represents the overlap ratio, indicating the degree of overlap between adjacent cropped images. To calculate the starting position of each crop, we determine the top-left corner coordinates (*x*_*start,i*_, *y*_*start,j*_) for each small image based on its index (*i, j*):

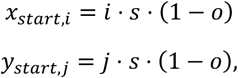

where *i* and *j* represent the crop index along the height and width. Specifically, *i* ranges from 0 to *r*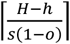 and *j* ranges from 0 to 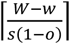.

After processing each cropped image through Mesmer, we reverse the tiling operation to reconstruct the large feature map, then apply 2D max pooling operation followed by flattening to produce the embedding for each sub-image. To aggregate the embeddings of the nine sub-images per well, we sequentially concatenate them to form a single overall embedding for each well.

Finally, we obtain the feature matrix 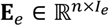 extracted by the extractor module, where *n* denotes the number of wells and *I*_*e*_ denotes the feature dimensions. We also refer to these as cpDistiller-extractor-based features.

### The joint training module

We design the joint training module to integrate CellProfiler-based features and cpDistiller-extractor-based features. To reduce potential noise and redundancy in the high-dimensional cpDistiller-extractor-based features, we initially use average pooling to reduce the feature dimensionality and smooth out irrelevant variation, resulting in the feature matrix **E**_*pooled*_ for further processing. Besides, we further employ two approaches to extract valuable information. For the first part, we use principal component analysis (PCA) to obtain the low-dimensional representation for each well, which we refer to as critical information:

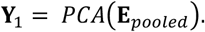

In the second part, we reshape **E**_*pooled*_ back into 2D feature maps and pass them through an attention module to get the global information **Y**_*2*_:

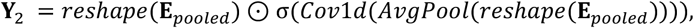

where the *AvgPool* represents a 2D average pooling operation, while *Cov*1*d* indicates a 1D convolution to capture inter-channel dependencies. *σ* denotes the Sigmoid function, which is used to produce a channel-wise attention map. ⊙ represents element-wise multiplication between the attention map and the reshaped **E**_*pooled*_.

To fusing critical and global information in the low-dimensional space, we apply element-wise addition as the encoding process for **E**_*pooled*_. The latent representations **z**_*e*_ for **E**_*pooled*_ are computed as:

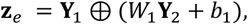

where ⊕ represents element-wise addition. *W*_*l*_ and *b*_*l*_ denote the weight and bias parameters of the encoder. Once the latent representations **z**_*e*_ are obtained, the decoder reconstructs the data back to the same dimensionality as **E**_*pooled*_.

Ultimately, we combine the CellProfiler-based features with the cpDistiller-extractor-based features transformed by the attention mechanism-based encoder-decoder architecture and feed the combined features into the technical correction module for further refinement.

### The technical correction module

The technical correction module removes triple effects from CP data and generates low-dimensional representations that maintain biological variation. Specifically, it consists of three parts: the Gaussian mixture variational autoencoder (GMVAE) as the core component, along with the contrastive learning module and the gradient reversal module.

### The Gaussian mixture variational autoencoder

Given the feature matrix **X**_*p*_ ∈ ℝ^*nxI*^*p* integrated by the joint training module, where *n* denotes the number of wells and *I*_*p*_ denotes the feature dimensions, the GMVAE takes the input **X**_*p*_ to obtain low-dimensional representations **Z**_*p*_. To illustrate the workflow of the GMVAE, we consider a sample **x** from **X**_*p*_. Since CP data encompasses numerous perturbations that may conform to distinct underlying Gaussian distributions, we use the GMVAE to identify these categorical distributions (pseudo-labels, denoted as **y**), which helps the model capture biological variation and ultimately contribute to biologically meaningful low-dimensional representations (denoted as **z**). The categorical distributions are inferred from the posterior distribution *q*_*ψ*_(**y**|**x**), which follow semi-supervised training pattern. Here, *q*_*ψ*_(**y**|**x**) is represented by a feedforward neural network as: *q*_*ψ*_(**y**|**x**) *= Cat*(*y*|*π*_*ψ*_(**x**)) and *π*_*ψ*_(**x**) is a probability vector. Since the categorical distribution cannot be backpropagated in the neural network, we use the Gumbel-Softmax distribution^51^ to facilitate gradient backpropagation, which allows the categorical distribution to be approximated using a continuous distribution.

In GMVAE, the objective is to optimize the Evidence Lower Bound (ELBO), which is expressed as follows:

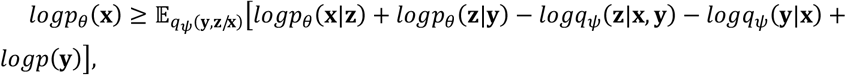

where these components can be broadly divided into three main optimization directions. The first optimization direction focuses on the reconstruction loss, represented by the expectation 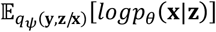. This loss is measured using mean squared error (MSE), which ensures that the low-dimensional representations **z** effectively capture the key information of the original input **x**.

The term 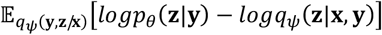 represents the Kullback-Leibler (KL) divergence between the variational posterior distribution *q*_*ψ*_(**z**|**x**, **y**) and the conditional prior distribution *p*_*0*_(**z**|**y**). This divergence ensures that the variational posterior distribution aligns with the prior, meaning that the learned latent representations **z** conform to the expected Gaussian distribution. Specifically, 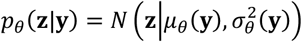 represents the prior Gaussian distribution conditioned on the category **y**, while 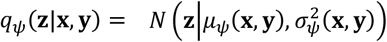 represents the variational posterior distribution conditioned on the input **x** and category **y**.

*q*_*ψ*_(**y**|x) provides the probability that **x** originates from various Gaussian distributions, Satisfying 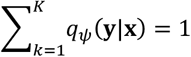, where *K* is the predefined number of Gaussian distributions. We use cross-entropy loss to refine the probabilities, guiding them towards high confidence regions. We treated the prior *p*(**y**) as a constant during loss backpropagation, as it does not influence the updates to the model’s parameters.

### The gradient reversal module

Within the low-dimensional space derived from the GMVAE, discriminators are employed to identify the source of each well’s representation **z**, specifically the batch, row, and column labels of the corresponding well. To implement adversarial learning similar to generative adversarial networks (GANs)^21^, different batches, rows, and columns can be treated as distinct domains. We then employ domain-adversarial learning, specifically the gradient reversal layer (GRL)^22^, to remove triple effects across these domains. Here, discriminators are denoted as: *D*_*batch*_, *D*_*row*_, *D*_*column*_, which are used to predict the batch, row, and column labels of **z**.

Specifically, the GRL reverses the gradient during backpropagation, causing the parameters of the GMVAE’s encoder, which acts as the generator, to be updated in the opposite direction of the discriminators, thereby achieving the adversarial objective. The GRL can be described as a pseudo-function: *GRL*_*λ*_(*x*). During the forward pass, the GRL functions as an identity operation, leaving the input parameters unchanged. However, during the backward pass, it scales the gradients from the following layers by −*λ* before sending them back to the preceding layers. The forward and backward passes are described by the following two equations:

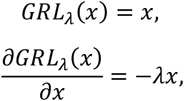

where *λ* is a hyperparameter that undergoes a non-linear transformation, varying from 0 to 1. In the early stages of training, its value is kept small to allow the discriminators to train sufficiently and develop discriminative capabilities. As training progresses, *λ* gradually increases to strengthen adversarial interactions between the encoder part of the GMVAE and the discriminators. The calculation formula for *λ* is as follows:

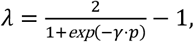

where *γ* is a hyperparameter, and *p* represents the percentage of the total iteration progress during training. To be specific, we denote the encoder part of the GMVAE as follows:

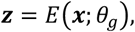

where *θ*_*g*_ denotes the learnable parameters of the GMVAE’s encoder. Subsequently, the low-dimensional representations **z** are passed through the GRL and discriminators, followed by the Softmax function to obtain the probability distribution for batch, row, and column predictions. We further describe the discriminators in detail as:

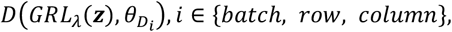

where 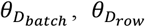 and 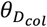 denote the learnable parameters of the batch, row, and column discriminators, which are updated to minimize the discriminator loss. Meanwhile, *θ*_*g*_ is updated through the GRL to maximize the discriminator loss, ensuring that the discriminators cannot distinguish the source of the low-dimensional representations, thereby obtaining representations free of technical effects.

Using the column discriminator loss as an example, the loss function is inspired by the label smoothing cross-entropy loss^52^. This approach is similarly applied to the batch and row discriminators for avoiding overconfidence:

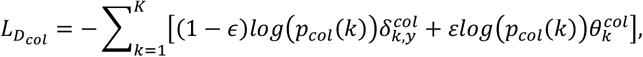

where **y** represents the index of the true label, and *ε* is a hyperparameter representing the proportion of soft labels considered when calculating the loss. 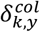 is a one-hot encoding, where the position corresponding to the true label is 1, with others set to 0. The probability that the **z** originates from *k* -th column, as predicted by the discriminator, is represented by *p*_*col*_(*k*). 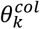 represents the soft labels, which indicates the weight assigned to the *k* -th column.

To be specific, standard cross-entropy loss focuses solely on optimizing the predicted probability of the true label, which can lead to overconfidence and neglect the uncertainty in the model’s predictions. Label smoothing cross-entropy loss addresses this limitation by redistributing a portion of the probability mass from the true label to the other classes, applying a uniform distribution across them to reduce overconfidence. However, this approach assigns equal importance to all non-true labels, which may not be appropriate for tasks where the predicted probabilities for non-true labels carry varying degrees of significance. In the context of our study, UMAP visualizations of the raw CP data show a gradient-influenced pattern that more distant column indexes exhibit more apparent column effects. This suggests that predicted probabilities closer to the true label index carry more meaning. To better capture this, we define a soft label distribution 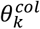, that reflects the differences among predicted probabilities, assigning varying significance to them based on their distance from the true label, while still avoiding overconfidence:

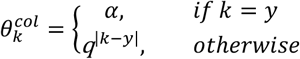

where the hyperparameter *α* represents the weight assigned to the true label, and *q* is the common ratio that determines the weights for the other labels based on their distance from the true label *y*. The value of *q* is determined by solving the higher-degree equation:

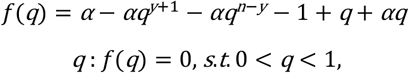

where *n* represents the number of categories for the technical effects (in this case, the number of column labels). The solution to this equation ensures that the soft label distribution is normalized, meaning that the sum of all probabilities equals 1, while also reflecting a geometric decay in significance as the distance from the true label index increases.

### The contrastive learning module

We first establish nearest neighbor relationships by utilizing *k*-nearest neighbors (KNN) intra technical effects and mutual nearest neighbors (MNN) inter technical effects, based on CellProfiler-based features and cpDistiller-extractor-based features processed with average pooling, using cosine distance as the similarity metric. This approach is applied to batch, row, and column effects, respectively, constructing the adjacency matrix that captures nearest neighbor relationships between wells for each effect. We then leverage the relationships identified across multiple technical effects to form triplets. We next use triplet loss^20^ to restore more accurate nearest-neighbor relationships for each well.

Specifically, we need to select a triplet (**a, p, n**) to act as the anchor, positive and negative samples. Each data point serves as the anchor in turn. For each anchor, data points with nearest neighbor relationships to that anchor are considered positive, while those without such relationships are considered negative. Specifically, (**a, p**) pairs are either in the *k*-nearest neighbors set 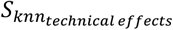 or in the mutual nearest neighbors set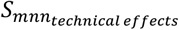. Conversely, (**a, n**) pairs are not found in either of two set. The specific formula is as follows:

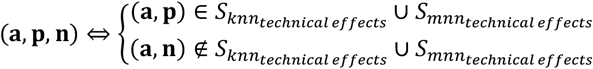

where 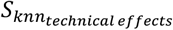 represents the set of all pairs of data points that originate from the same type of technical effects and are *k*-nearest neighbors within the same category. 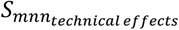 represents the set of all pairs of data points that originate from the same type of technical effects but are mutual nearest neighbors in different categories.

The triplet loss function is not directly applied to the latent space of the GMVAE. Instead, the representations **z** need to pass through a nonlinear projection head, as previous research has demonstrated that adding such a layer can improve the quality of the learned representations^53^. This architecture can be represented as *ph*(**z**):

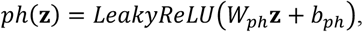

where the parameters *W*_*ph*_ and *b*_*ph*_ represent the learnable weights and biases of a linear layer.

Then we use the triplet loss to remove triple effects:

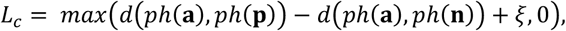

where *d*(*ph*(**a**), *ph*(**p**)) represents the Euclidean distance between the anchor and positive samples after passing through the projection head, and the *ξ* is a hyperparameter.

If we aim to correct row and column effects, the overall loss can be written as follows:

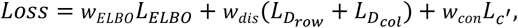

where *L*_*c*′_ represents the triplet loss calculated by considering well position effects.

If we need to correct batch, row and column effects, the overall loss can be expressed as follows:

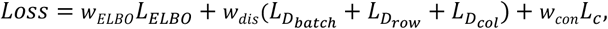

where *L*_*c*_ represents the triplet loss calculated by considering triple effects. In the above loss functions, the weights of *w*_*ELBO*_, *w*_*dis*_ and *w*_*con*_ are the weighting coefficients assigned to different components of the loss function. These coefficients control the relative importance of the ELBO, discriminator losses, and the triplet loss in the overall optimization process.

The parameters 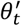 of the final trained model are obtained through exponential moving average (EMA)^54^, which can be given by:

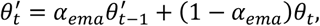

where *θ*_*t*_ represents the weighted model parameters obtained at round *t*, and *α*_*ema*_ is a hyperparameter. All training hyperparameters are available in the training details.

### Training details

For the extractor module, the overlap ratio *o* is set to 0.25 and the sliding stride *s* is set to 256. Max pooling with kernel size and step size both set to 16, is applied to merge feature maps from nine sub-images, yielding the output dimension *I*_*e*_ to 11,664, corresponding to the 108 × 108 feature map. For the joint training module, average pooling with kernel size and step size both set to 9, is applied to smooth out irrelevant variation. For the technical correction module, the hidden dimensions of the encoder and decoder are set to 512, while the hidden dimensions of discriminators are set to 128. The latent space dimensionality of cpDistiller is set to 50. The technical correction module uses LeakyReLU with default parameters as the activation function throughout, except for the variance inference component, which utilizes the Softplus activation function. The projection head consists of a linear layer with an output dimension of 50 and a LeakyReLU activation function. The optimizer used is AdamW, with the learning rate set to 3e-3 for the discriminators and 1e-3 for the other parts. Although we optimized the adversarial learning, mode collapse is still a common issue. When considering 7,638 CellProfiler-based features, the default learning rate is unsuitable for training and can lead to collapse. Therefore, we adjusted the initial learning rate to half of the default value. The *γ* is set to 10 in GRL. The soft label hyperparameter *α* is set to 0.75, and the hyperparameter *ϵ* in label smoothing cross-entropy loss is set to 0.1. For the contrastive learning, the *ξ* is set to 10, and the nearest neighbor hyperparameters for MNN and KNN are set to 5 and 10, respectively. The default number of epochs is 50, with *α*_*ema*_ set to 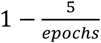. In the loss functions, the weights of *w*_*ELBO*_, *w*_*dis*_ and *w*_*con*_ are set to 1, 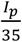and 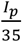, respectively. The experimental environment includes two 24GB Nvidia 4090 graphics cards and 96 Intel(R) Xeon(R) Gold 5318N CPUs @ 2.10GHz.

### Implementation details of downstream analyses

The establishment of gene groups based on scRNA-seq data: we used the gget tool, a Python package available at https://github.com/pachterlab/gget, which enables efficient querying of the top 100 similar genes of the input gene calculated by the ARCHS4^36^ based on scRNA-seq data. Concretely, we input the 12,602 genes experimented in the cpg0016 dataset and retrieved the top 100 most similar genes for each input gene, forming 12,602 gene sets. Since a similarity score between 0.6 and 0.8 was considered significant^55^, we selected 0.6 as the threshold to include more genes and got more applicable gene sets. Finally, we intersected each gene set with the genes present in the cpg0016 dataset to obtain the final gene groups of similar genes.

Gene Ontology (GO) term analysis: we utilized the Enrichr^56^ platform to conduct GO term enrichment analysis^57^ for each gene group, assigning relevant GO terms to the genes. We specifically focused on Cellular Component (CC) analysis to identify the organelles associated with each gene and ranked the results by their statistical significance.

Gene relationship analysis: we used the GeneMANIA tool^41^ to evaluate the relationship of genes in different groups. We submitted the genes in gene groups to GeneMANIA and used the default network selections. Specifically, GeneMANIA first identifies genes that are functionally similar or share properties with the submitted genes, then builds the network connecting both the submitted and similar genes. It then displays the gene network regarding co-expression networks, physical interaction, genetic interaction, co-localization, pathways, and predicted and shared protein domain information.

Screening for active genes: following the approach established by ref.^14^, we calculated the average of repetitions for each perturbation at the same position across five different plates to obtain the average representation for each perturbation. Considering that negative treatments generally do not induce significant phenotypic changes, we calculated the Euclidean distances between negative treatments and set the 95th percentile of these distances as the threshold for identifying active genes. Overexpression treatments were classified as active if their Euclidean distance from the negative treatments exceeded this threshold. In Batch_1, 226 genes were identified as active genes.

Ranking the significance of the clusters in single-link hierarchical clustering: we applied a perturbation approach to rank the significance of clusters in single-link hierarchical clustering. For each cluster, we randomly selected the same number of data points as contained in that cluster. We then calculated the minimum clustering distance among these randomly selected points. This perturbation process was repeated 1000 times for each cluster, generating 1000 distance scores. By comparing the actual cluster distance with the distribution of the 1000 perturbed distance scores, we ranked the clusters and got the ranking of the actual cluster, which provides an indication of the significance level of data point aggregation within each cluster.

The calculation of fraction retrieved in gene and compound retrieval tasks: we calculated and compared the fraction retrieved scores for gene-gene and gene-compound simulated retrieval tasks using cpDistiller-derived representations and CellProfiler-based features in the cpg0000 dataset, following the workflow in ref.^47^.

### Pre-processing for CellProfiler-based features

We utilized features pre-extracted with CellProfiler from the prior study^14^, which included up to 7,638 features for images with both brightfield and fluorescence images and 4,752 features for only fluorescence images, after removing features containing NaN values. Additionally, we followed the feature selection process described in the study^14^ to select 1,446 features. Due to data quality issues reported in the original dataset^14^, we excluded data from Batch_12 and the BR00123528A plate. For pre-computed feature transformation, we followed the Pycytominer^58^ and applied its default operations, including z-score normalization on the entire dataset.

### Data pre-processing for raw images

To match the dual-channel input format required by Mesmer of the extractor module, we extracted data from three channels: DNA, RNA, and ER. The DNA channel primarily pertains to the nucleus, while the RNA and ER channels are related to the cytoplasm. After computing these with their respective illumination files, we combined them into two separate channels. The processed data was then stored in NPZ format to prepare the image data for the extractor module. For image pre-processing, we utilized the procedure from Mesmer, which included using Contrast Limited Adaptive Histogram Equalization (CLAHE) to enhance local contrast, followed by logarithmic smoothing of the data in the first channel.

### Evaluation metrics

To assess the effectiveness of different methods in removing technical effects, we used three technical correction metrics: average silhouette width (ASW)^26^, technic average silhouette width (tASW)^27^, and graph connectivity^27^. To evaluate biological preservation, four metrics were used: isolated label F1^27^, isolated label silhouette^27^, perturbation average silhouette width (pASW)^27^, and normalized mutual information (NMI)^27^. For a more intuitive and clear comparison, we normalized all metrics to a range of 0 to 1, where higher values indicated better results. Metrics involving perturbation labels, excluding ASW and tASW, which reflected the clustering of same perturbations and the dispersion of different ones, were calculated on the selected subset. The reasons for these calculation approaches are based on two main challenges. First, the perturbation labels in the JUMP dataset are not well-defined, making it difficult to accurately cluster similar perturbations. Specifically, the ORF perturbations induced by different reagents may target functionally similar genes, making some perturbations inherently difficult to distinguish. Besides, with over 10,000 unique perturbation labels in ORF data, it is difficult to separate functionally overlapping or densely labeled perturbations. Second, category imbalance and numerous categories pose a challenge for accurately calculating clustering metrics. To be specific, perturbations in controls replicated substantially more frequently than those in the treatment, often by several to dozens of times. Metrics like pASW, which do not account for category imbalance, result in biased scores. Metrics like NMI typically require unsupervised algorithms, such as Leiden^59^ to generate predicted labels. However, the numerous and less well-defined perturbation labels in the JUMP dataset pose additional challenges, leading to inaccuracies in the metric calculations.

To address the issues mentioned above, we focused on certain perturbations. For correcting well position effects, we used controls, including five positive controls (JCP2022_037716, JCP2022_035095, JCP2022_050797, JCP2022_012818, and JCP2022_915132) and four negative controls (JCP2022_915128, JCP2022_915129, JCP2022_915130, and JCP2022_915131). For removing triple effects, we selected perturbations induced by 34 different compounds that showed better clustering on target plates. First, these selections reduce the impact of unwell-defined perturbation labels, allowing for more reliable clustering of similar perturbations. In particular, positive and negative controls theoretically should have significant or negligible effects on cell phenotypes, resulting in marked differences in their impacts^14^. Furthermore, different positive controls are expected to exhibit distinct effects on cell phenotypes. Besides, for target plates, perturbations are shared across different batches, which can be used for aligning data^15^. Additionally, perturbations caused by the same compounds in target plates that clustered well in the raw data should remain well-clustered after technical correction. Otherwise, it may indicate overcorrection, as true biological differences should persist even after correction. Second, these selections help address category imbalance and reduce categories, allowing for more accurate calculation of clustering metrics. Specifically, the number of positive and negative controls are generally consistent, and each perturbation in target plates was duplicated approximately 20 times, ensuring a consistent number of tests.

The specific calculation of above metrics, which follow the approach of previous works by scArches^26^ and scIB^27^ for analyzing single-cell data, are further detailed in Supplementary Text 7.

### Baseline methods

To evaluate technical correction in CP data, we compared several widely used single-cell analysis methods: Seurat v5 (v5.0.1)^23^, Harmony (v0.0.9)^19^, Scanorama (v1.7.4)^24^, scVI (v0.14.6)^18^, and scDML (v0.0.1)^25^. UMAP plots, used to compare embeddings generated by different methods, were created with the following parameters: *n_neighbors=15, min_dist=0*.*1*, and *random_state=9000*. For removing well position effects, data points within each batch were standardized using z-score. When correcting triple effects, z-score standardization was applied within each batch, followed by integration across batches and an additional round of normalization to mitigate batch effects. All methods, including cpDistiller, were applied to CP data that had undergone the above preprocessing steps to ensure consistency.

For Harmony, Seurat v5 and Scanorama, which can correct multiple technical effects, the workflow involved sequentially passing the row and column labels if considering removing well position effects. When aiming to remove triple effects, these methods required first passing the batch labels, followed sequentially by the row and column labels. For methods that are limited to correcting only one type of technical effects, if considering well position effects, we selected row and column effects separately as correction targets, resulting in two results, respectively. Considering triple effects, we focused solely on batch effects as they represent the most significant influence. Specific implementation details are as follows:

Seurat v5: we followed the example pipeline provided by the Seurat v5 for integrative analysis. The labels for technical effects were passed sequentially into the *split* function. We skipped the *NormalizeData* function and *FindVariableFeatures* function, as these are specifically tailored for scRNA-seq data. During the correction phase, the *CCA* method was employed to remove technical effects. All parameters were used with default settings.

Harmony: dimensionality reduction was performed using *scanpy*.*tl*.*pca* to the default size of 50. The labels for technical effects were passed sequentially into the *scanpy*.*external*.*pp*.*harmony_integrate* function. All parameters were used with default settings.

Scanorama: dimensionality reduction was performed using *scanpy*.*tl*.*pca* to the default size of 50. The labels for technical effects were passed sequentially into the *scanpy*.*external*.*pp*.*scanorama_integrate* function. All parameters were used with default settings.

scVI: we followed the example tutorial provided by the scVI on GitHub for processing scRNA-seq data. We did not use the *scanpy*.*pp*.*highly_variable_genes* function, which is specifically tailored for scRNA-seq data. In the preprocessing stage, we followed the preprocessing operations of previous work to process each feature as follows: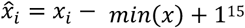. Other operations were used with default settings.

scDML: we followed the example tutorial on GitHub provided by scDML. We did not use the *scanpy*.*pp*.*log1p* function and the *scanpy*.*pp*.*highly_variable_genes* function, which are specifically tailored for scRNA-seq data. Other operations were used with default settings.

## Data availability

We downloaded well-level images and the features extracted by CellProfiler, following the recommended workflow^14^. For the JUMP datasets, raw stained images and the features extracted by CellProfiler are accessible via AWS at https://registry.opendata.aws/cellpainting-gallery/.

## Code availability

The cpDistiller software, along with detailed documentation and tutorials, is freely available at https://github.com/Cell-Painting/cpDistiller.

## Acknowledgements

This work was supported by the National Key Research and Development Program of China grants no. 2020YFA0908700 (J.L.), 2020YFA0908702 (J.L.) and 2024YFA1307703 (S.C.), the National Natural Science Foundation of China grants no. 62473212 (S.C.), 62203236 (S.C.) and 62272246 (J.L.), and the Young Elite Scientists Sponsorship Program by CAST grant no. 2023QNRC001 (S.C.).

