## supplemental file for "Triple-effect correction for Cell Painting data with contrastive and domain-adversarial learning"

### Contents

**Supplementary Texts** ... **4**

**Supplementary Text 1** ... **4**

**Supplementary Text 2** ... **5**

**Supplementary Text 3** ... **6**

**Supplementary Text 4** ... **7**

**Supplementary Text 5** ... **8**

**Supplementary Text 6** ... **9**

**Supplementary Text 7** ... **10**

**Supplementary Figures** ... **13**

**Supplementary Figure 1** ... **13**

**Supplementary Figure 2** ... **14**

**Supplementary Figure 3** ... **15**

**Supplementary Figure 4** ... **16**

**Supplementary Figure 5** ... **17**

**Supplementary Figure 6** ... **18**

**Supplementary Figure 7** ... **19**

**Supplementary Figure 8** ... **20**

**Supplementary Figure 9** ... **21**

**Supplementary Figure 10** ... **22**

**Supplementary Figure 11** ... **23**

**Supplementary Figure 12** ... **24**

**Supplementary Figure 13** ... **25**

**Supplementary Figure 14** ... **26**

**Supplementary Figure 15** ... **27**

**Supplementary Figure 16** ... **28**

**Supplementary Figure 17** ... **29**

**Supplementary Figure 18** ... **30**

**Supplementary Figure 19** ... **31**

**Supplementary Figure 20** ... **32**

**Supplementary Figure 21** ... **33**

**Supplementary Figure 22** ... **34**

**Supplementary Figure 23** ... **35**

**Supplementary Figure 24** ... **36**

**Supplementary Figure 25** ... **37**

### Supplementary Texts

#### Supplementary Text 1: Discussion on the insights and value of features extracted by the cpDistiller-extractor module

To investigate if the well position effects in the Cell Painting (CP) data are due to the inherent biases of the CellProfiler[1], we implemented the extractor module based on the pre-trained segmentation model Mesmer[2]. After validating that these effects were not due to the biases, we focused on evaluating the quality and utility of the features extracted by cpDistiller-extractor module. We applied Mesmer to open reading frame (ORF) overexpression dataset in cpg0016 from the terabyte-scale Joint Undertaking for Morphological Profiling (JUMP) dataset[3] and found that Mesmer produced highly accurate cell segmentation results, clearly delineating cell boundaries and nuclei (Supplementary Fig. 6a,b). We also explored the capabilities of several widely used pre-trained models, as detailed in Supplementary Figs. 7-13. Notably, Mesmer exceled in accurately segmenting cell nuclei in CP images compared to other models.

Mesmer was originally trained on TissueNet[2], which focused on cell nuclei and the whole cell segmentation in tissue samples. Deep learning-based feature extraction methods, like Mesmer, rely heavily on large amounts of high-quality, annotated data to learn valuable patterns. Although there were currently no annotated CP image datasets available for further fine-tuning, Mesmer was still able to effectively extract essential information from CP images, such as cell nuclei, cell placement, clustering and variations in lighting, as shown in Supplementary Fig. 6b. These findings confirmed that the cpDistiller-extractor module, built on Mesmer, was capable of deriving meaningful insights from CP images.

**Supplementary Text 2: Settings used in pre-trained models**

For the YOLO segmentation task, we selected a range of models with varying parameter sizes implemented by Ultralytics[4], including YOLOv8m-seg, YOLOv8n-seg, and YOLOv8x-seg, all used with default settings. Similarly, for the YOLO detection task, we utilized models with different parameter sizes, including YOLOv8m, YOLOv8n, and YOLOv8x, all applied with default configurations. In the LACSS framework[5], we employed the pre-trained cnsp4_bf weights using the default settings. For image creation within Mesmer, the first channel was set to green and the second channel was set to blue. Mesmer provides two default parameter sets tailored for post-processing: one for whole-cell segmentation and the other for nuclear segmentation. For whole-cell segmentation, the default settings include a maximum threshold of 0.075, no smoothing for maxima detection, an interior threshold of 0.2, an interior smoothing value of 2, a threshold of 15 for removing small objects, a hole-filling threshold of 15, and a radius parameter of 2. In contrast, the nuclear segmentation defaults set a slightly higher maxima threshold of 0.1, while other parameters such as smoothing, interior threshold, interior smoothing, thresholds for small objects and hole filling, and the radius value remain the same as in whole-cell segmentation. These carefully chosen parameters are essential for defining how the segmentation algorithm identifies and processes different cellular components, ensuring precise and consistent segmentation results.

**Supplementary Text 3: cpDistiller can effectively correct technical effects just using CellProfiler-based features**

cpDistiller introduces the extractor module to obtain richer information from raw CP images and uses the joint training module to integrate cpDistiller-extractor-based features with CellProfiler-based features to perform technical correction. However, the capability of technical correction may arise from the introduction of cpDistiller-extractor-based features, rather than the cpDistiller's inherent ability. To validate whether cpDistiller itself is effective, we conducted experiments on ORF data in 12 batches using only CellProfiler-based features. By using quantitative metrics and performing one-sided paired Wilcoxon signed-rank tests to quantitatively evaluate the effectiveness of cpDistiller in correcting well position effects across 12 batches, we confirmed that cpDistiller significantly outperformed all baseline methods (Supplementary Fig. 29a). Similarly, when correcting triple effects, cpDistiller consistently surpassed other baseline methods (Supplementary Fig. 29c). The results demonstrated the effectiveness of cpDistiller in correcting technical effects and preserving biological variation when processing CellProfiler-based features independently.

Additionally, we measured the time and memory usage of cpDistiller compared to other baseline methods just using CellProfiler-based features on Batch_1. Although cpDistiller ranked fourth overall, it's important to note that Harmony[6] and Scanorama[7] perform technical correction on low-dimensional representations, which enhances the algorithm's efficiency but may also lead to information loss (Supplementary Fig. 29b). Moreover, scDML[8] is designed to correct only one type of technical effects, giving it inherent advantages in terms of time and memory consumption.

#### Supplementary Text 4: Ablation experiments on cpDistiller

To demonstrate the impact of cpDistiller-extractor-based features and validate the effectiveness of the extractor module and joint training module, we conducted the ablation experiments across 12 batches. In these experiments, the model that excluded both the extractor and joint training modules, thus relying solely on CellProfiler-based features, was referred to as cpDistiller-C.

We observed that cpDistiller outperformed cpDistiller-C in metrics assessing the removal of well position effects across 12 batches, specifically graph connectivity[9] (Supplementary Fig. 30). Similarly, cpDistiller showed better performance in preserving biological information across all biological metrics (Supplementary Fig. 30). These results indicated that cpDistiller ensured a more stable removal of both row and column effects while better preserving biological variation with cpDistiller-extractor-based features. Furthermore, we used one-sided paired Wilcoxon signed-rank tests for a more detailed quantification of performance differences across the 12 batches. The results showed that incorporating cpDistiller-extractor-based features, along with joint training, led to a significant improvement in performance compared to the model using only CellProfiler-based features, with a $P$-value of 0.098.

#### Supplementary Text 5: The ability of cpDistiller to achieve incremental learning

Harmony is the most effective methods for correcting both row and column effects in baseline methods for ORF data and has shown strong performance in removing batch effects for compound data in recent benchmark analyses[10]. However, it has limitations in scalability. When processing new data, Harmony must re-align the new data with the original data, modifying the low-dimensional representations and requiring reprocessing[10]. These approaches can be computationally intensive and inefficient, especially with large datasets.

In contrast, cpDistiller supports incremental learning. To validate cpDistiller's capability, we partitioned the ORF data into two groups: the original dataset (Batch_1 to Batch_6), representing existing public data, and the new dataset (remaining batches), representing newly acquired data. After training cpDistiller on the original dataset to correct triple effects simultaneously, the saved model parameters can be directly applied to the new dataset. As shown in Fig. 4, there were significant batch effects between the original and new datasets, along with well position effects within both datasets. Nonetheless, uniform manifold approximation and projection (UMAP) visualizations[11] showed that cpDistiller successfully corrected batch, row, and column effects in the new dataset using the pre-trained parameters on the original dataset, without the need for re-alignment or retraining (Supplementary Fig. 31a). These results demonstrated cpDistiller's scalability and efficiency in leveraging incremental learning to seamlessly process new dataset.

**Supplementary Text 6: The robustness of cpDistiller to the feature dimensions**

CellProfiler-based features can be redundant, leading to efforts in feature selection to streamline the data[12]. The 1446-dimensional CellProfiler-based features are obtained through selection processes[3]. Incorporating additional potentially redundant or noisy features extracted by the CellProfiler raises the feature dimensionality to 4,752 for only fluorescence images and 7,638 when combining both fluorescence and brightfield images. We also conducted experiments using these high-dimensional features, demonstrating that cpDistiller maintains robustness to feature selection.

First, we conducted experiments on ORF data in 12 batches and assessed the performance by using quantitative metrics to demonstrate the ability of cpDistiller in correcting well position for a single batch. We found that cpDistiller was highly effective in correcting well position while preserving biological variation, consistently outperforming other baseline methods in terms of average performance across all feature combinations over 12 batches (Supplementary Fig. 31b). Besides, we also conducted experiments for simultaneously correcting triple effects, and cpDistiller still demonstrated remarkable stability across different feature dimensions, surpassing all baseline methods (Supplementary Fig. 31c). These results highlighted the robustness of cpDistiller to feature selection, consistently delivering superior performance in both correcting well position effects within single batch and simultaneously removing triple effects across batches.

#### Supplementary Text 7: Details of evaluation metrics

For assessing the effectiveness of different methods in removing technical effects, we used three metrics: average silhouette width (ASW)[13], technic average silhouette width (tASW)[9], and graph connectivity[9]. ASW can be used to evaluate the clarity of silhouettes of categories after clustering, reflecting the degree of closeness of sample points to elements within their class compared to elements outside their class. ASW ranges from -1 to 1, where value closer to 1 indicates well-separated categories, and value closer to 0 suggests categories are mixed and difficult to distinguish. When using ASW to measure the effectiveness of technical correction, it can be calculated as follows:

$$ASW_{technical\ correction} = 1 - abs(ASW).$$

In our study, ASW can be calculated separately for batch, row, and column, with higher values indicating more effective technical correction.

tASW is used to evaluate the effectiveness of removing technical effects. Unlike ASW, tASW is calculated by selecting different types of perturbations and computing the average ASW value for each perturbation type. The $tASW_i$ for a given perturbation type $P_i$, calculated across all data points $j$ belonging to that type, is as follows:

$$tASW_i = \frac{1}{|P_i|}\sum_{j \in P_i} 1 - |ASW(j)|.$$

The final score is determined by averaging calculated tASW for each perturbation type $T$:

$$tASW = \frac{1}{|T|}\sum_{j \in T} tASW_i.$$

tASW takes perturbation labels into account, expecting the same perturbations across different batches, rows, and columns to be mixed. In our study, tASW can be calculated separately based on batch, row, and column, with higher values indicating more effective technical correction.

We calculate the $k$-nearest neighbors (KNN) subgraph for each perturbation type, hoping that the perturbation labels on each KNN subgraph contain only the expected type. The high graph connectivity score indicates that the same perturbations can cluster well after removing technical effects. Graph connectivity is calculated by averaging the connectivity of each perturbation's KNN subgraph:

$$Graph\ connectivity = \frac{1}{|D|}\sum_{j=1}^{D}\frac{|LCC_j|}{N_j},$$

where $\left|LCC_j\right|$ represents the number of cells in the largest connected component of perturbation $j$'s $k$-nearest neighbor graph, and $N_j$ represents the number of the perturbation $j$. Graph connectivity ranges from 0 to 1, with higher value indicating better technical correction.

For assessing the effectiveness of different methods in preserving biological variation, we used four metrics: perturbation average silhouette width (pASW)[9], isolated label F1[9], isolated label silhouette[9], and normalized mutual information (NMI)[9].

pASW measures the clustering of same perturbations and is calculated as follows:

$$pASW = \frac{1 + ASW}{2},$$

where value closer to 1 indicates a stronger clustering effect of the same perturbations, demonstrating the model's ability to maintain biological variation.

Isolated labels represent perturbation types present in the fewest technical labels, which can be calculated separately for batch, row and column. If there are multiple isolated labels, the result is calculated as the average of all scores. To determine how well isolated labels are separated from others in the latent representation, we optimize their clustering assignment through the F1 score of isolated labels at different resolutions. The optimal F1 score for isolated labels is used as the final score, defined as:

$$F1 = 2\frac{precision \times recall}{precision + recall}.$$

Besides, we calculate the pASW between isolated and non-isolated labels on the latent representations, following the same procedure as the definition of the isolated label F1 score. If multiple isolated labels are present, the final score is calculated as the average of all scores. Isolated label scores can be calculated separately for batch, row, and column, with higher values indicating clearer separation between isolated labels and other categories in the latent representation.

The normalized mutual information (NMI) metric measures the similarity between two clusters. Here, clustering labels are generated by the Leiden algorithm[14], with resolutions set from 0.1 to 2 in increments of 0.1. Each time, the labels generated by the Leiden algorithm are compared to the true labels, specifically perturbation labels, to calculate the NMI score, with the highest result used as the final score. The specific calculation for NMI is:

$$NMI(Y, C) = \frac{2 \times I(Y; C)}{H(Y) + H(C)},$$

where $Y$ represents the true labels and $C$ represents the predicted labels generated by the Leiden algorithm. $I(Y; C)$ represents the mutual information between the true labels and predicted labels, and $H(\cdot)$ represents entropy.

### Supplementary Figures

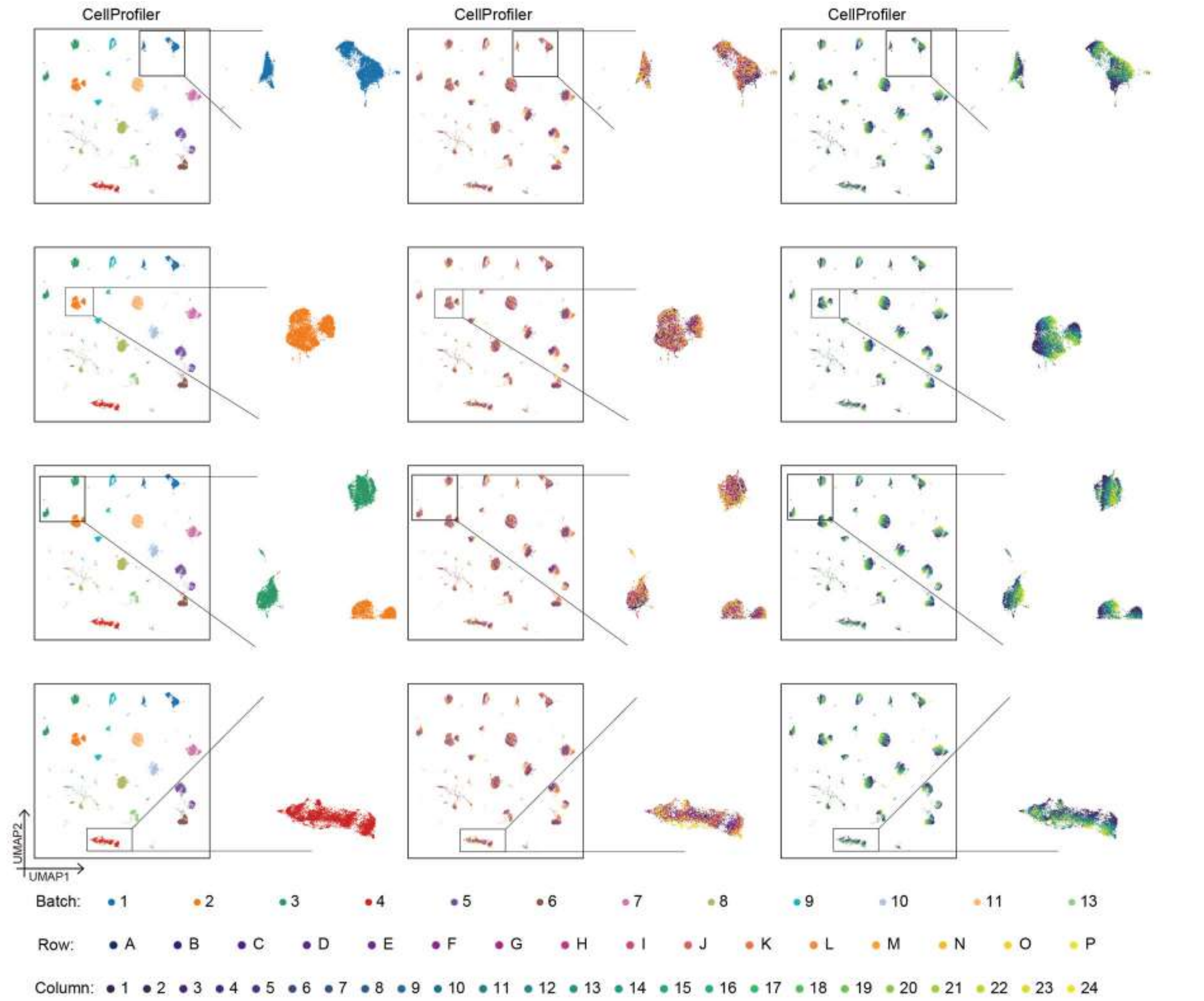

**Supplementary Figure 1. The UMAP visualization of CellProfiler-based features (Batch_1-Batch_4).** The UMAP visualizations of CellProfiler-based features, from top to bottom, represent data from Batch_1-Batch_4 of the ORF data in the JUMP dataset, each colored by batch, row, and column, respectively.

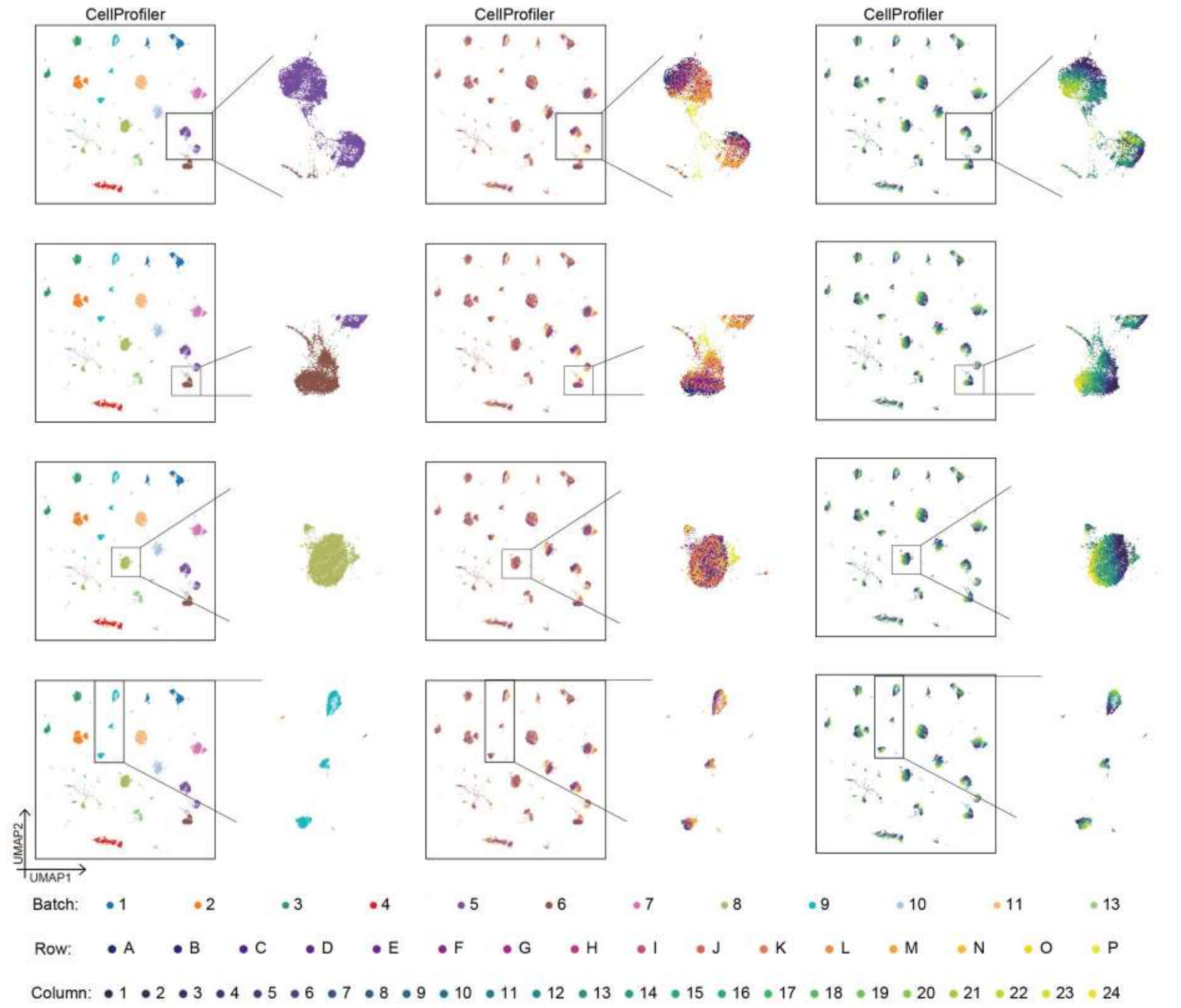

**Supplementary Figure 2. The UMAP visualization of CellProfiler-based features (Batch_5, Batch_6, Batch_8, and Batch_9).** The UMAP visualizations of CellProfiler-based features, from top to bottom, represent data from Batch_5, Batch_6, Batch_8, and Batch_9 of the ORF data in the JUMP dataset, each colored by batch, row, and column, respectively.

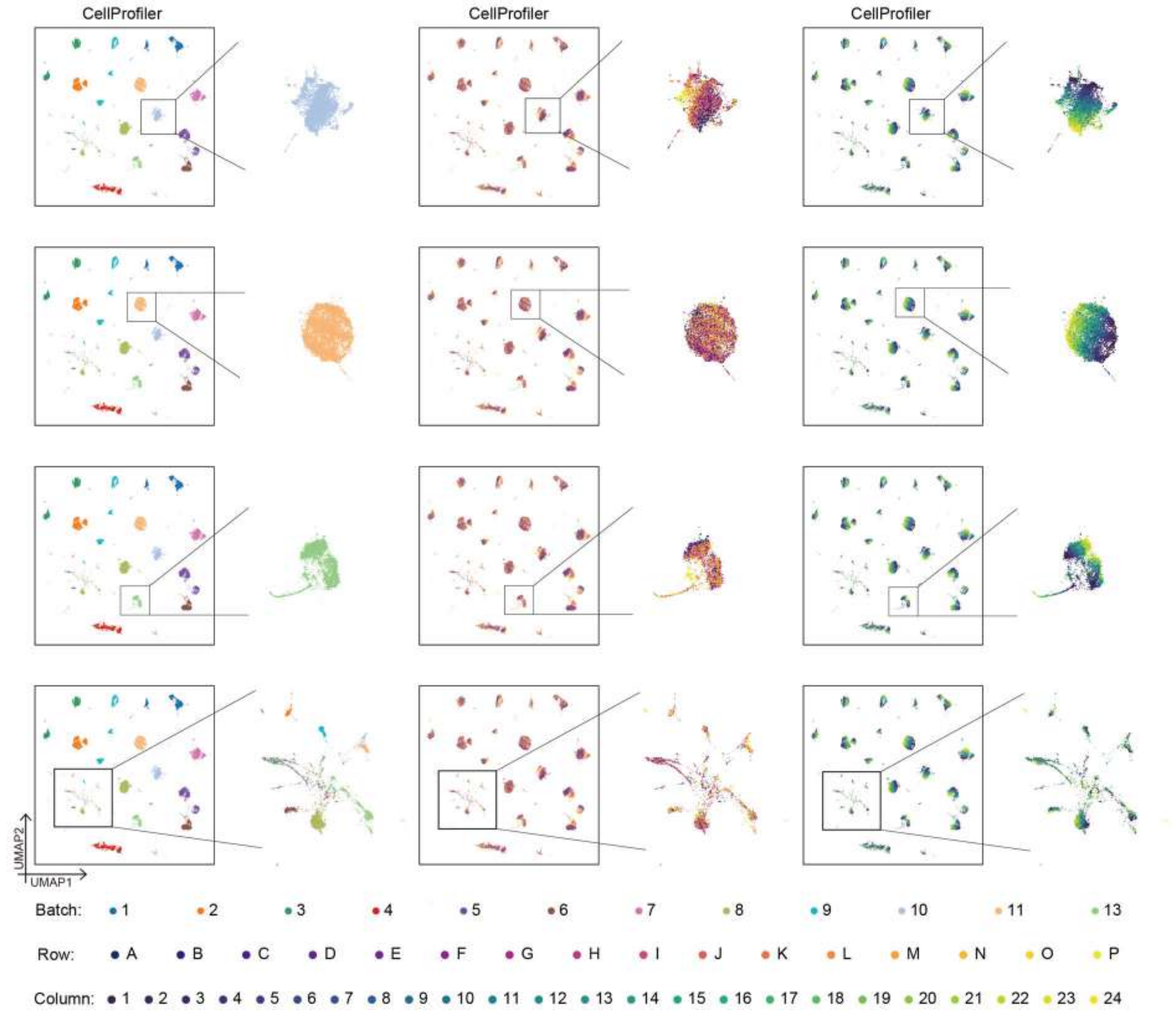

**Supplementary Figure 3. The UMAP visualization of CellProfiler-based features.** The UMAP visualizations of CellProfiler-based features, from top to bottom, represent data from Batch_10, Batch_11, Batch_13 of the ORF data in the JUMP dataset, with the final visualization showing a mixed subset of data from Batch_1–Batch_11 and Batch_13. Each plot is colored by batch, row, and column, respectively.

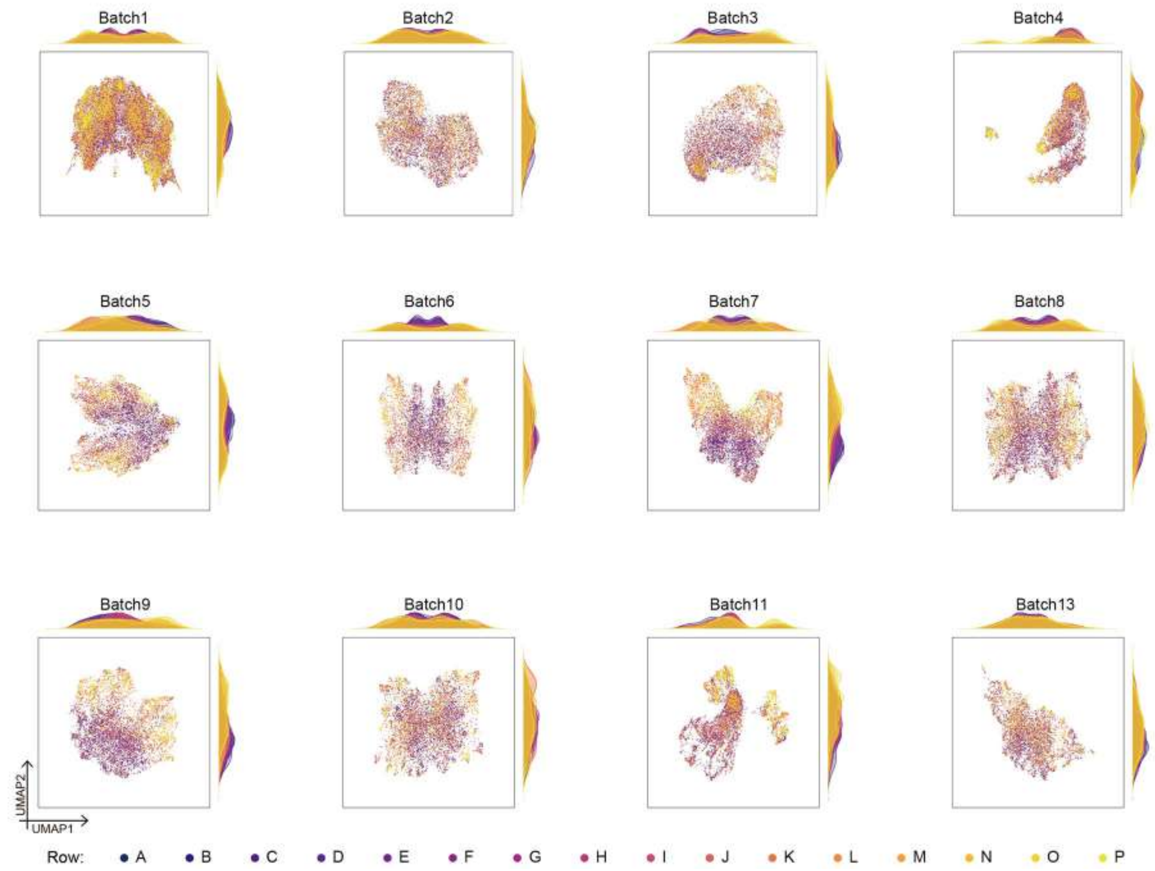

**Supplementary Figure 4. The UMAP visualization of cpDistiller-extractor-based features, colored by row.** The UMAP visualization of the cpDistiller-extractor-based from Batch_1-Batch_11 and Batch_13 of the ORF data in the JUMP dataset, colored by row.

241

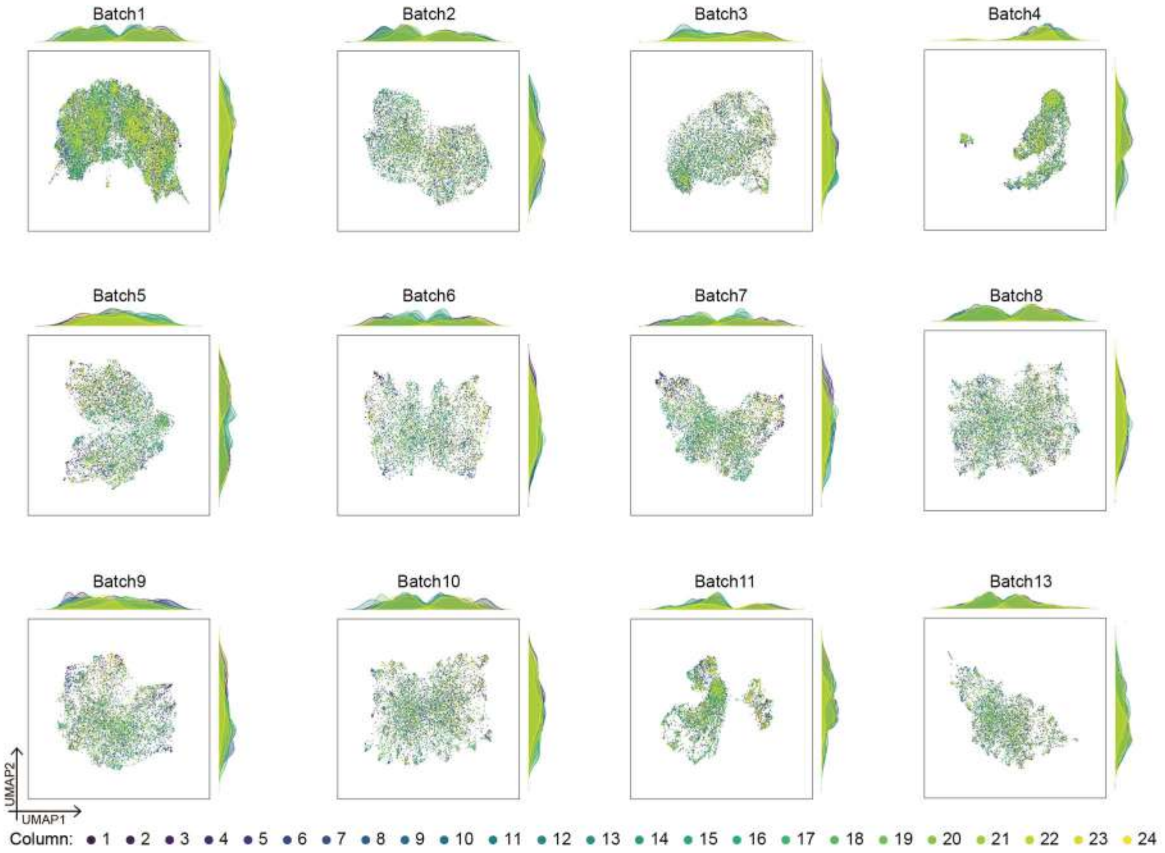

**Supplementary Figure 5. The UMAP visualization of cpDistiller-extractor-based features, colored by column.** The UMAP visualization of the cpDistiller-extractor-based from Batch_1-Batch_11 and Batch_13 of the ORF data in the JUMP dataset, colored by column.

242

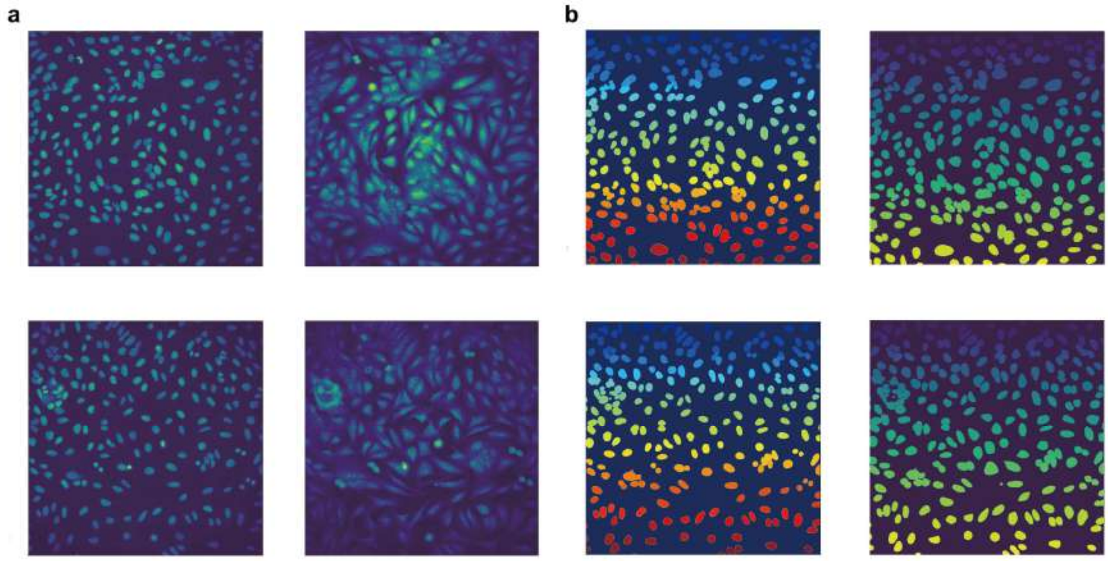

**Supplementary Figure 6. Segmentation results of the Mesmer model. a**, Cell nuclei and dual-channel cell images. **b**, Segmentation results for cell nuclei and whole cells in the JUMP dataset using the Mesmer model.

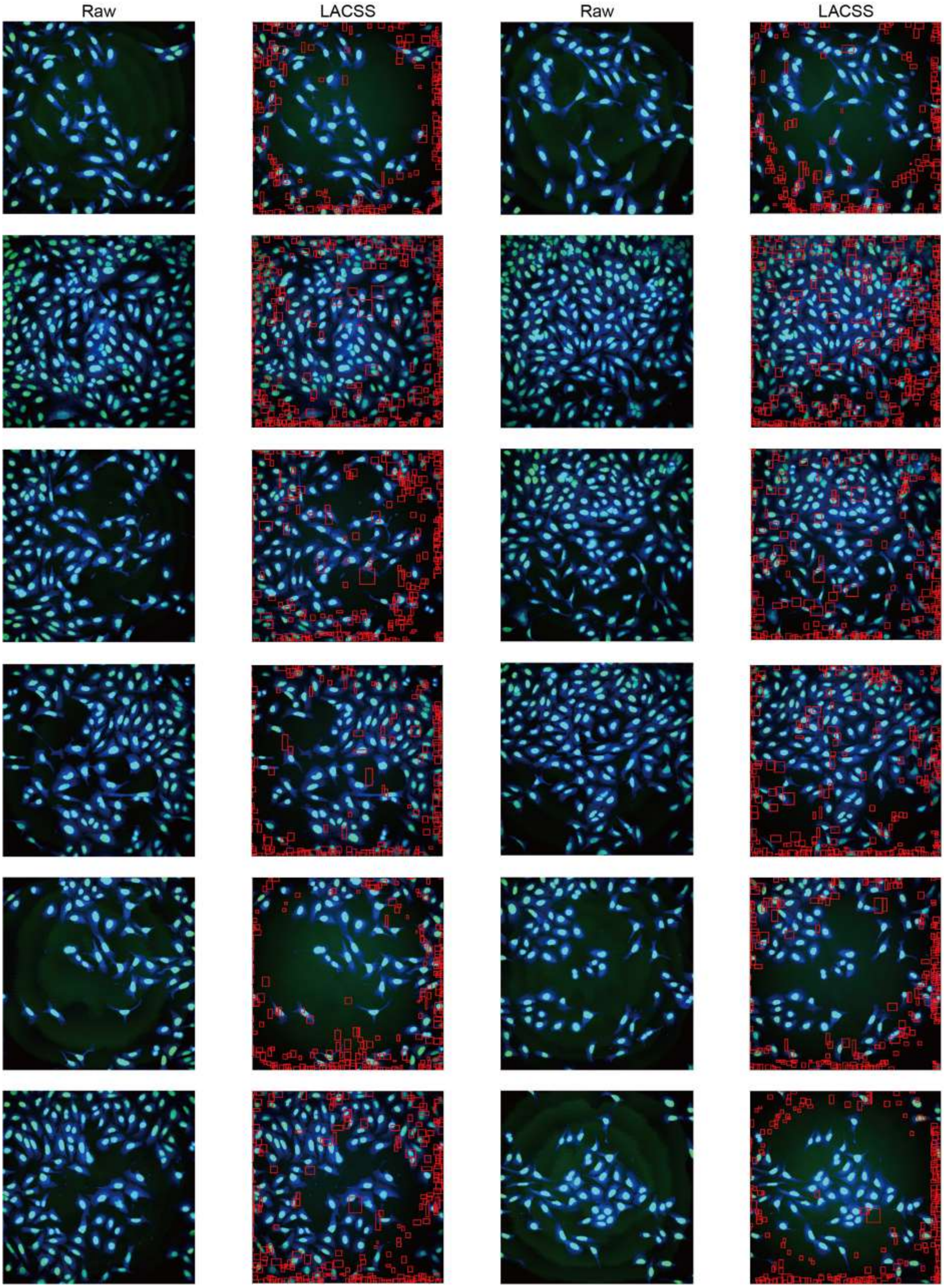

**Supplementary Figure 7. Detection results of the LACSS model.** Cell detection results using the LACSS model on Batch_1 images of the ORF dataset for wells A01 to A03 in plate BR00117035. The first and third columns display raw images, and the second and fourth columns show detection results.

244

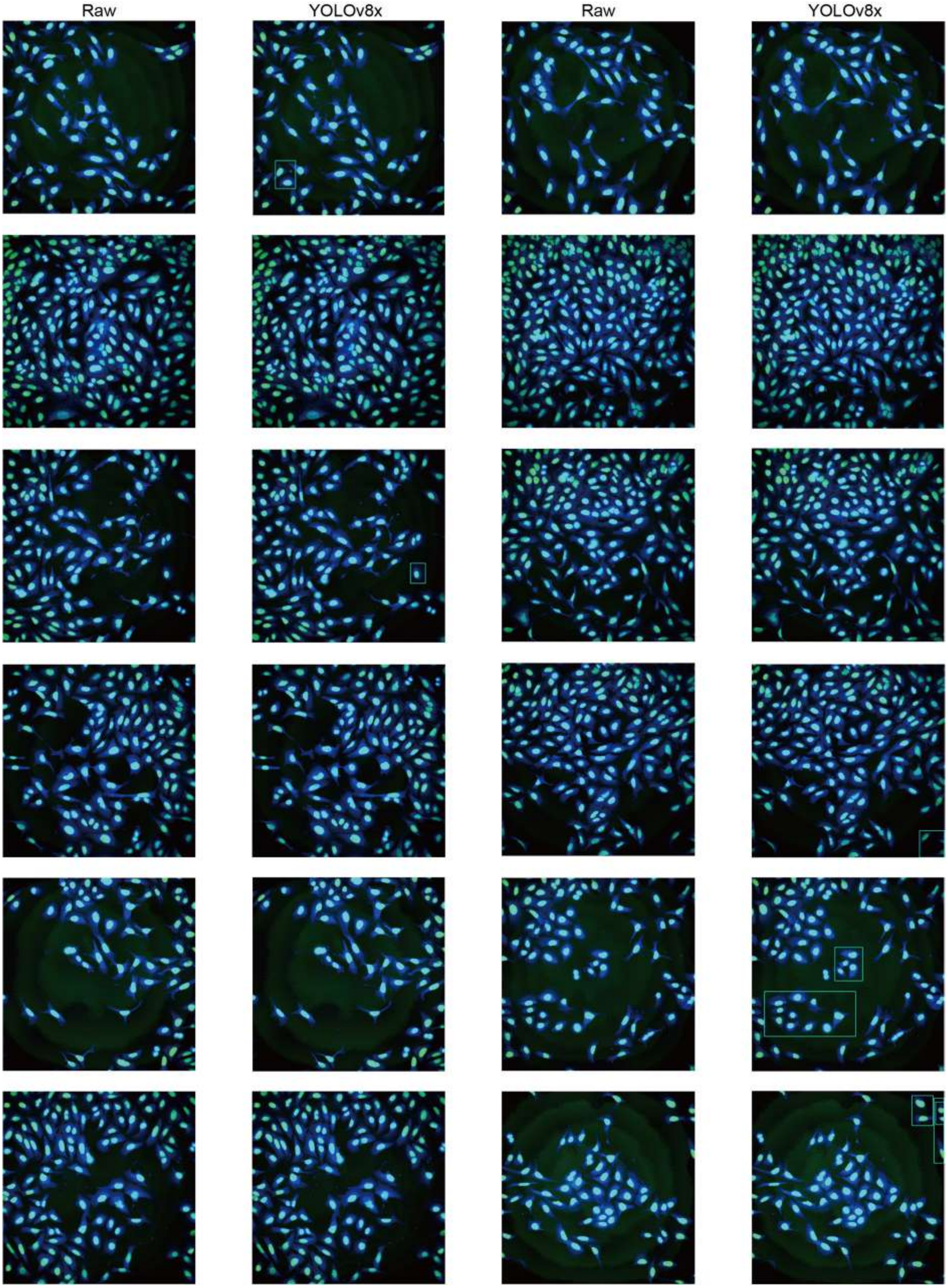

**Supplementary Figure 8. Detection results of the YOLOv8x model.** Cell detection results using the YOLOv8 model (YOLOv8x.pt) on images of the ORF dataset for wells A01 to A03 in plate BR00117035. The first and third columns display raw images, and the second and fourth columns show detection results.

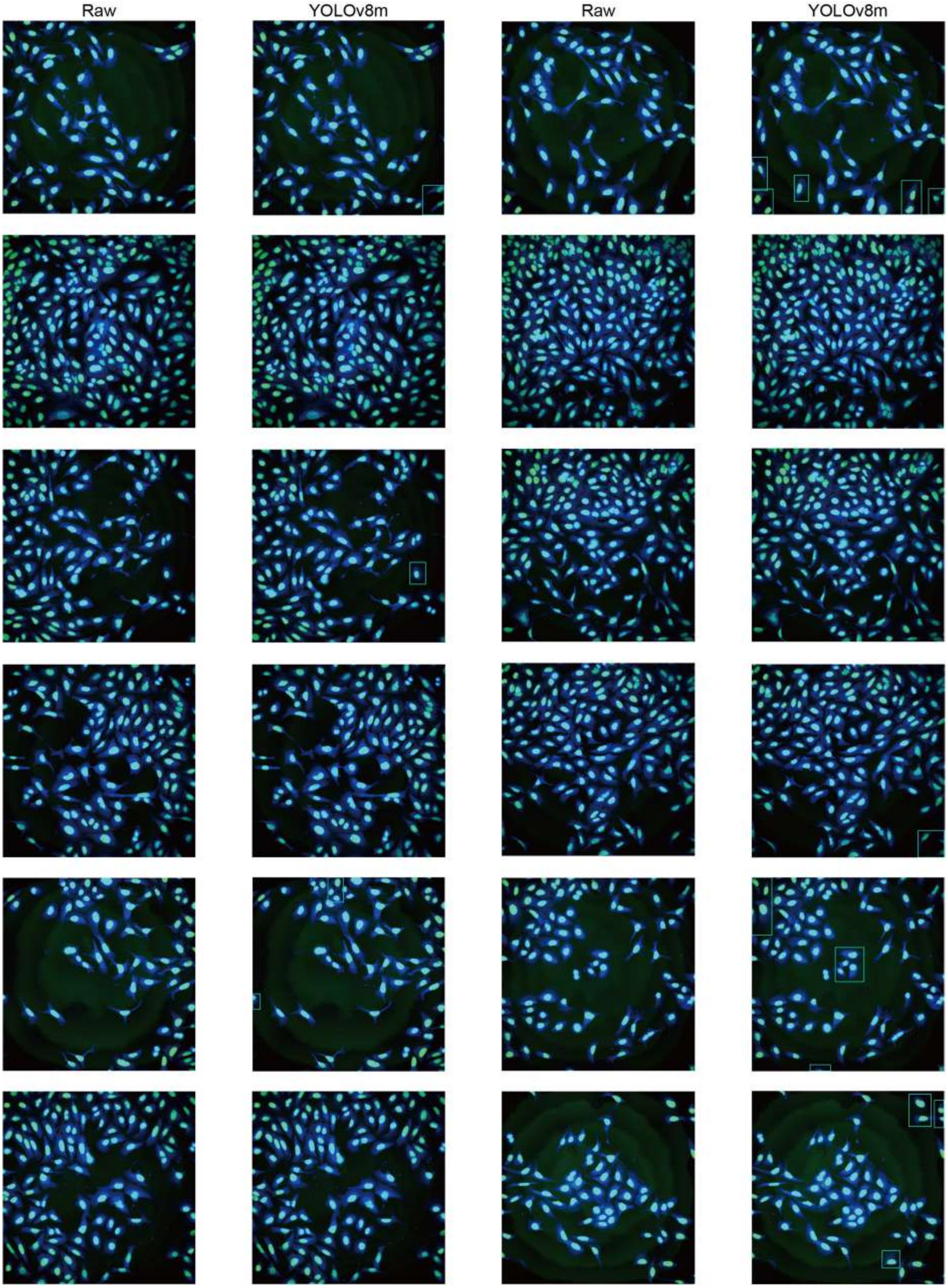

**Supplementary Figure 9. Detection results of the YOLOv8m model.** Cell detection results using the YOLOv8 model (YOLOv8m.pt) on images of the ORF dataset for wells A01 to A03 in plate BR00117035. The first and third columns display raw images, and the second and fourth columns show detection results.

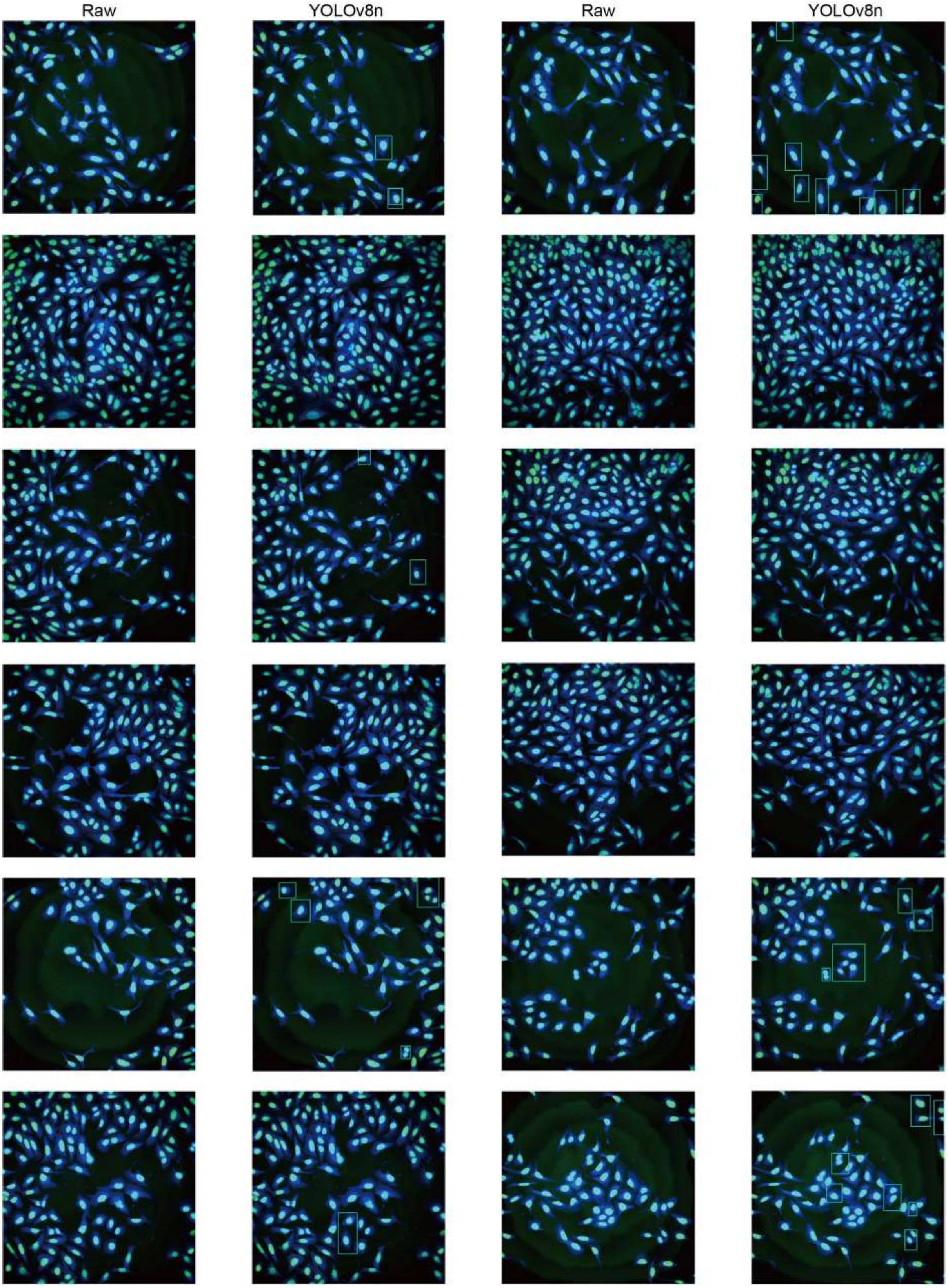

**Supplementary Figure 10. Detection results of the YOLOv8n model.** Cell detection results using the YOLOv8 model (YOLOv8n.pt) on images of the ORF dataset for wells A01 to A03 in plate BR00117035. The first and third columns display raw images, and the second and fourth columns show detection results.

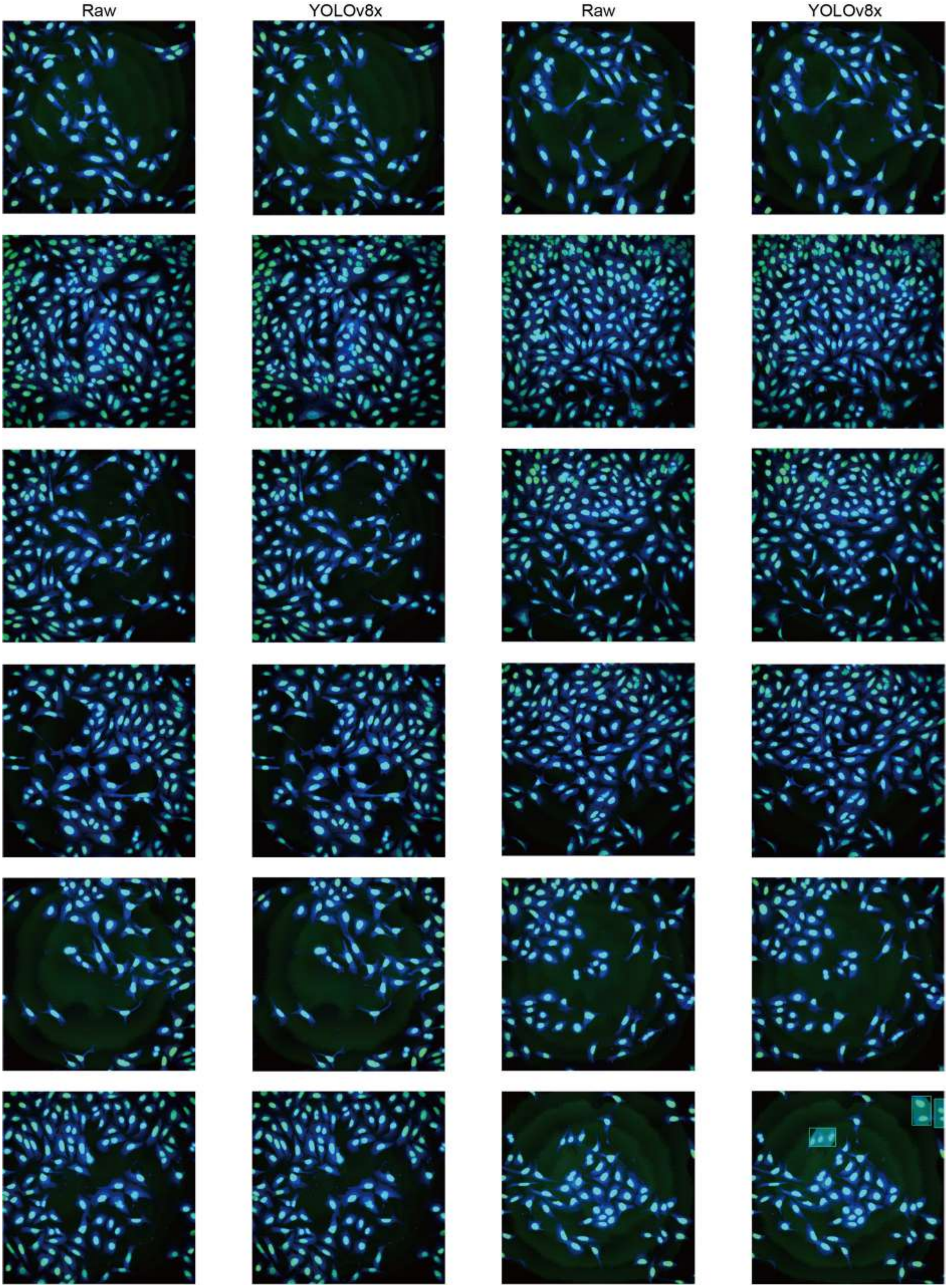

**Supplementary Figure 11. Segmentation results of the YOLOv8x model.** Cell segmentation results using the YOLOv8 model (YOLOv8x-seg.pt) on images of the ORF dataset for wells A01 to A03 in plate BR00117035. The first and third columns display raw images, and the second and fourth columns show segmentation results.

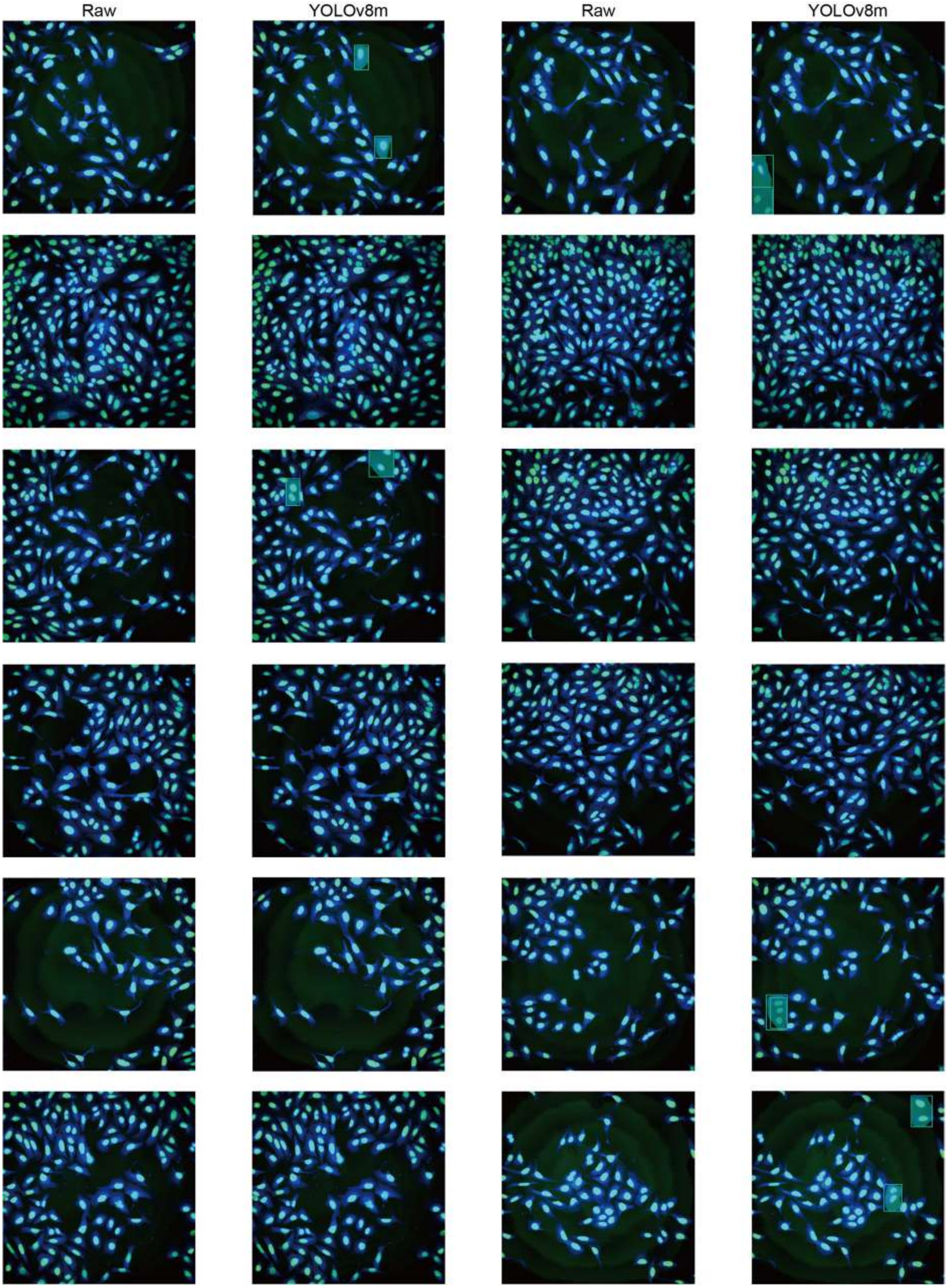

**Supplementary Figure 12. Segmentation results of the YOLOv8m model.** Cell segmentation results using the YOLOv8 model (YOLOv8m-seg.pt) on images of the ORF dataset for wells A01 to A03 in plate BR00117035. The first and third columns display raw images, and the second and fourth columns show segmentation results.

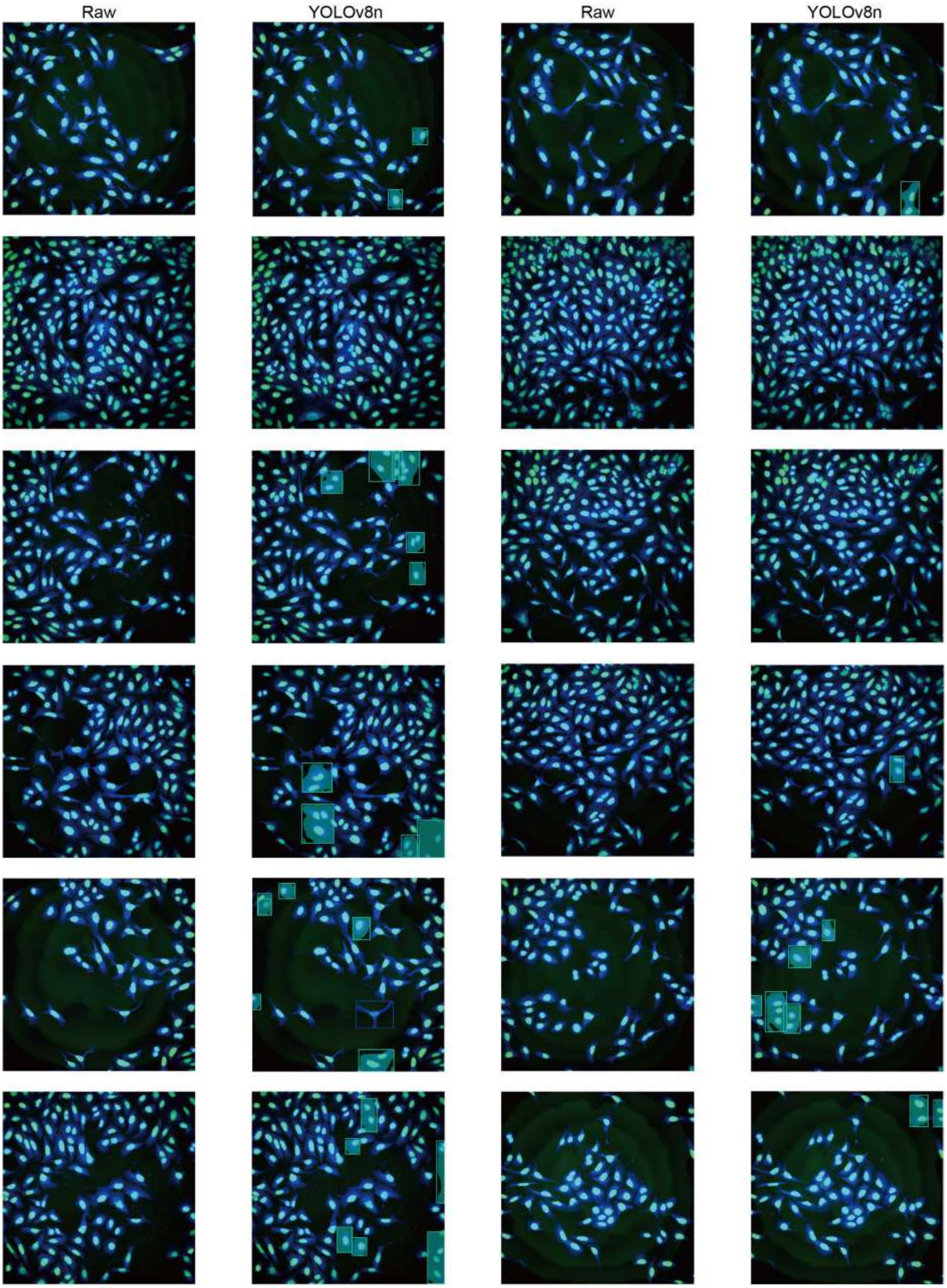

**Supplementary Figure 13. Segmentation results of the YOLOv8n model.** Cell segmentation results using the YOLOv8 model (YOLOv8n-seg.pt) on images of the ORF dataset for wells A01 to A03 in plate BR00117035. The first and third columns display raw images, and the second and fourth columns show segmentation results.

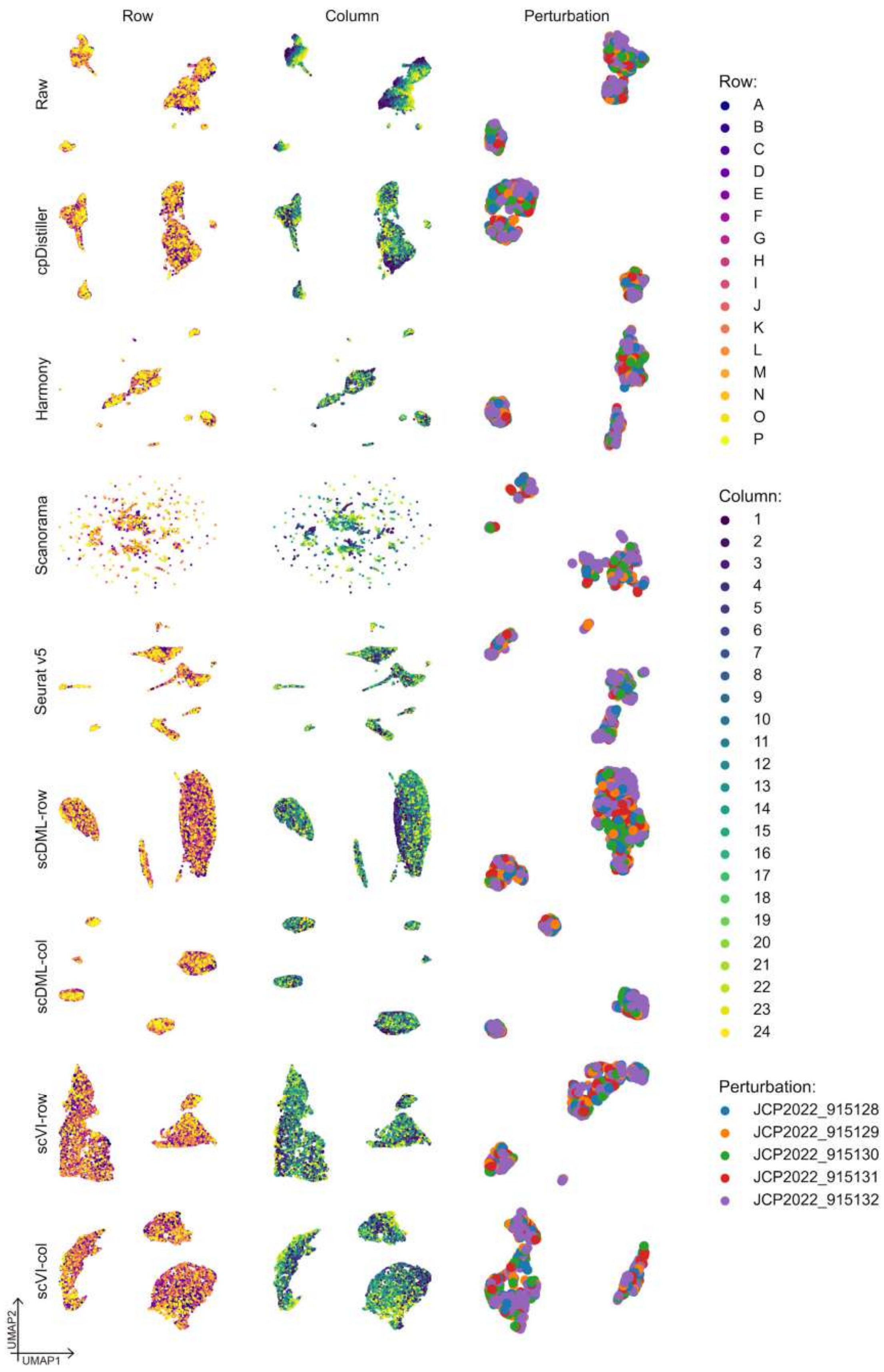

**Supplementary Figure 14. UMAP visualizations of embeddings obtained by different methods in Batch_1 of the ORF dataset.**

251

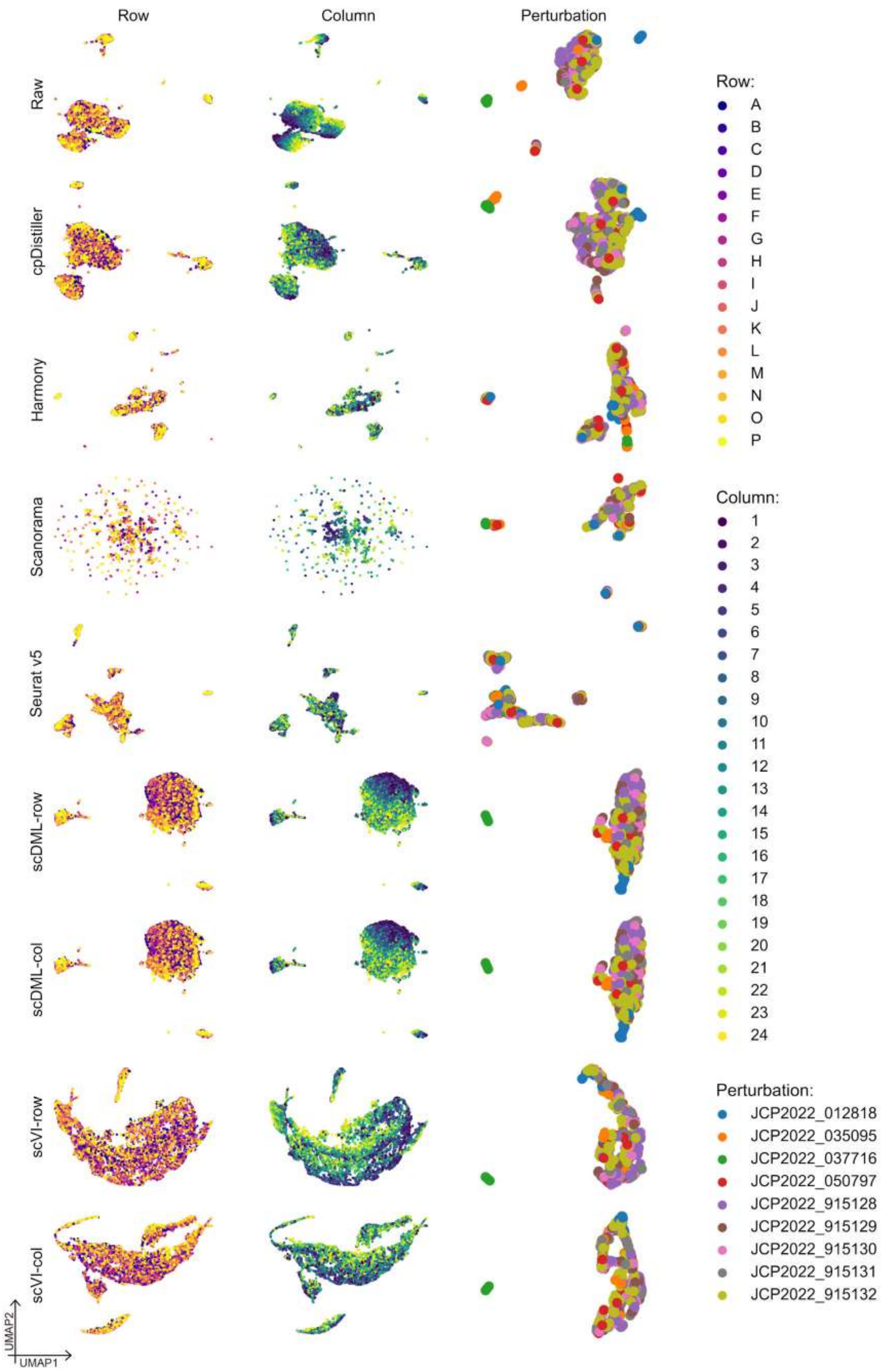

252 **Supplementary Figure 15. UMAP visualizations of embeddings obtained by different methods in Batch_2 of the ORF dataset.**

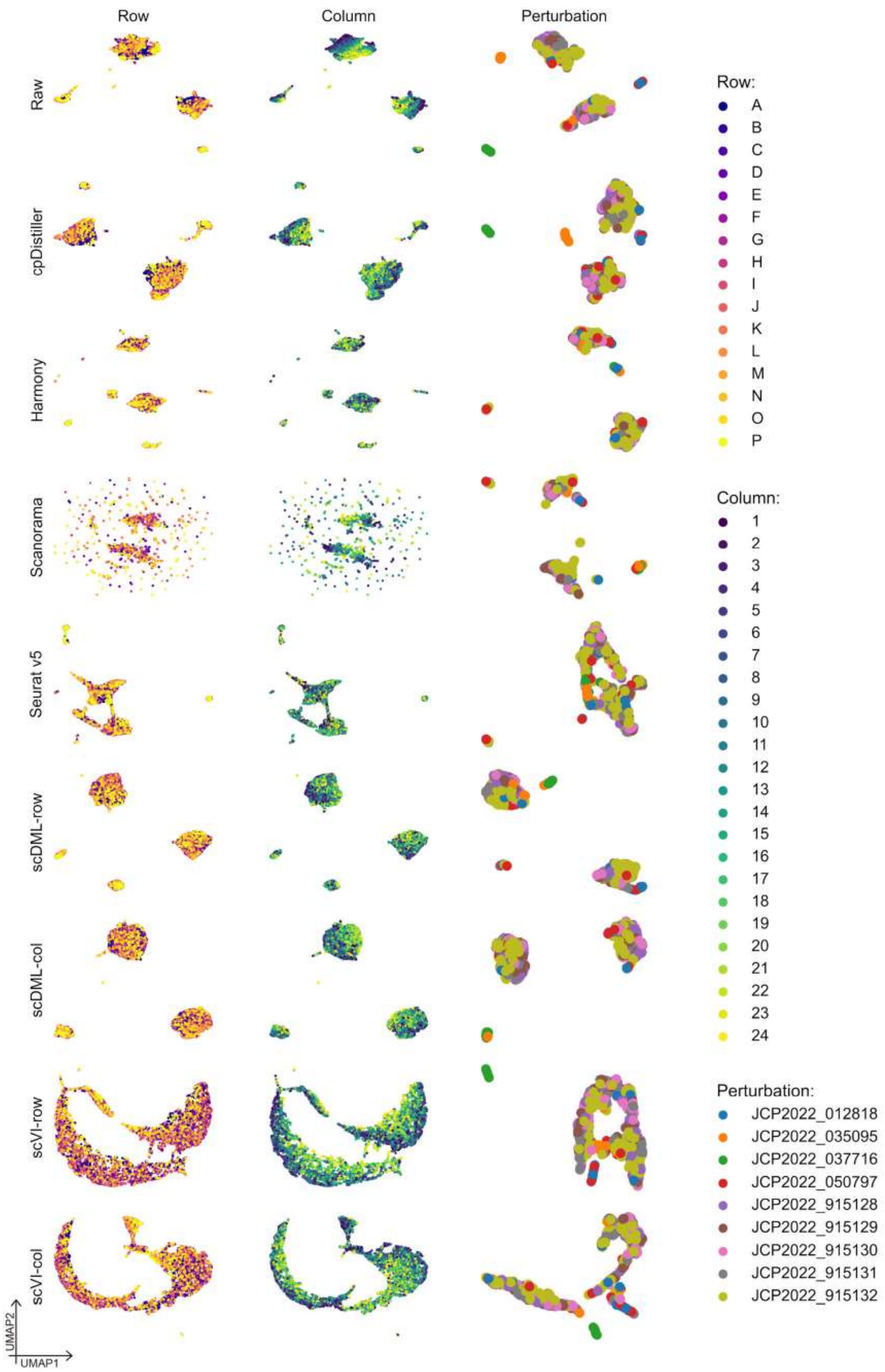

**Supplementary Figure 16. UMAP visualizations of embeddings obtained by different methods in Batch_3 of the ORF dataset.**

253

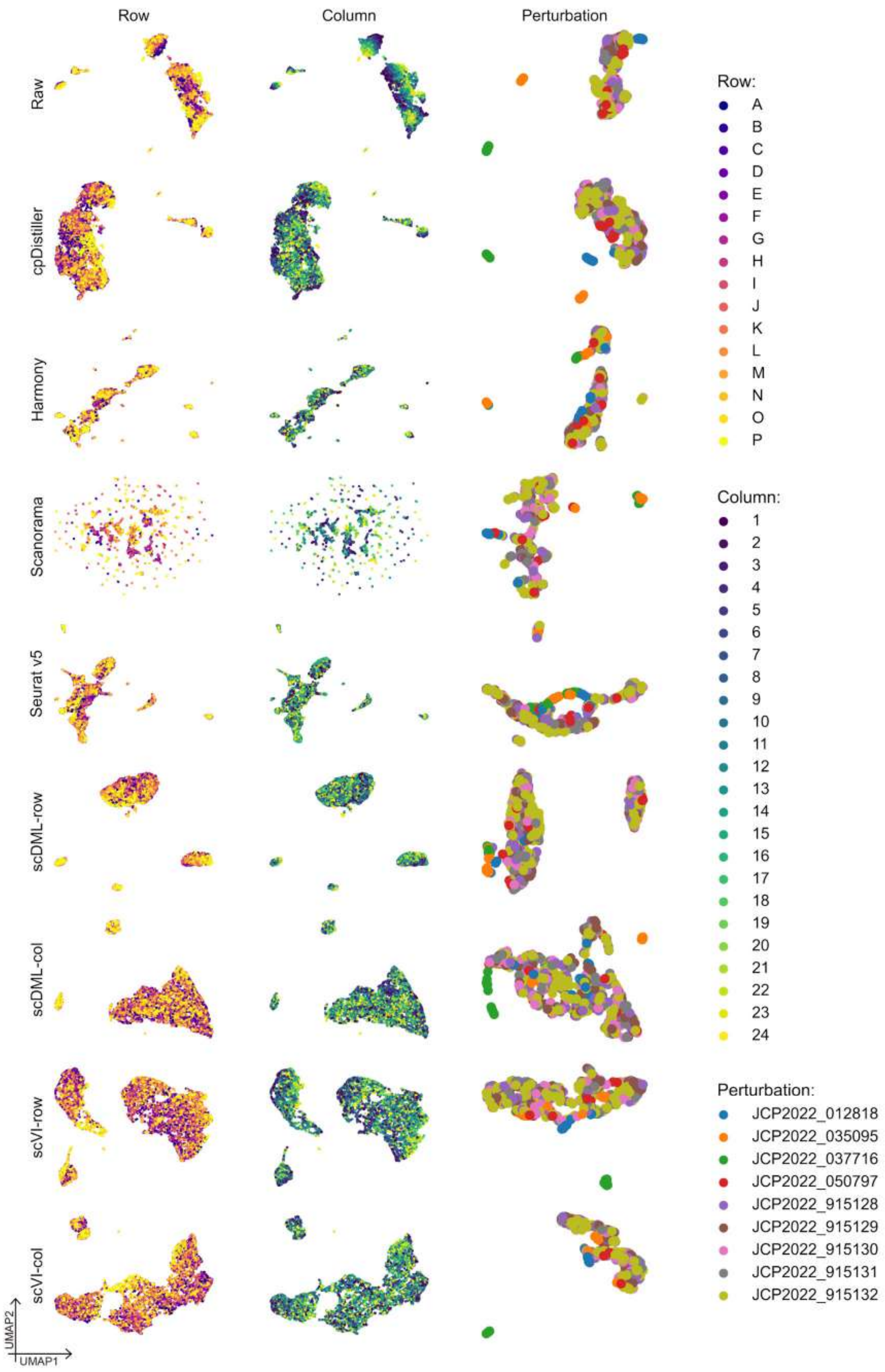

**Supplementary Figure 17. UMAP visualizations of embeddings obtained by different methods in Batch_4 of the ORF dataset**.

254

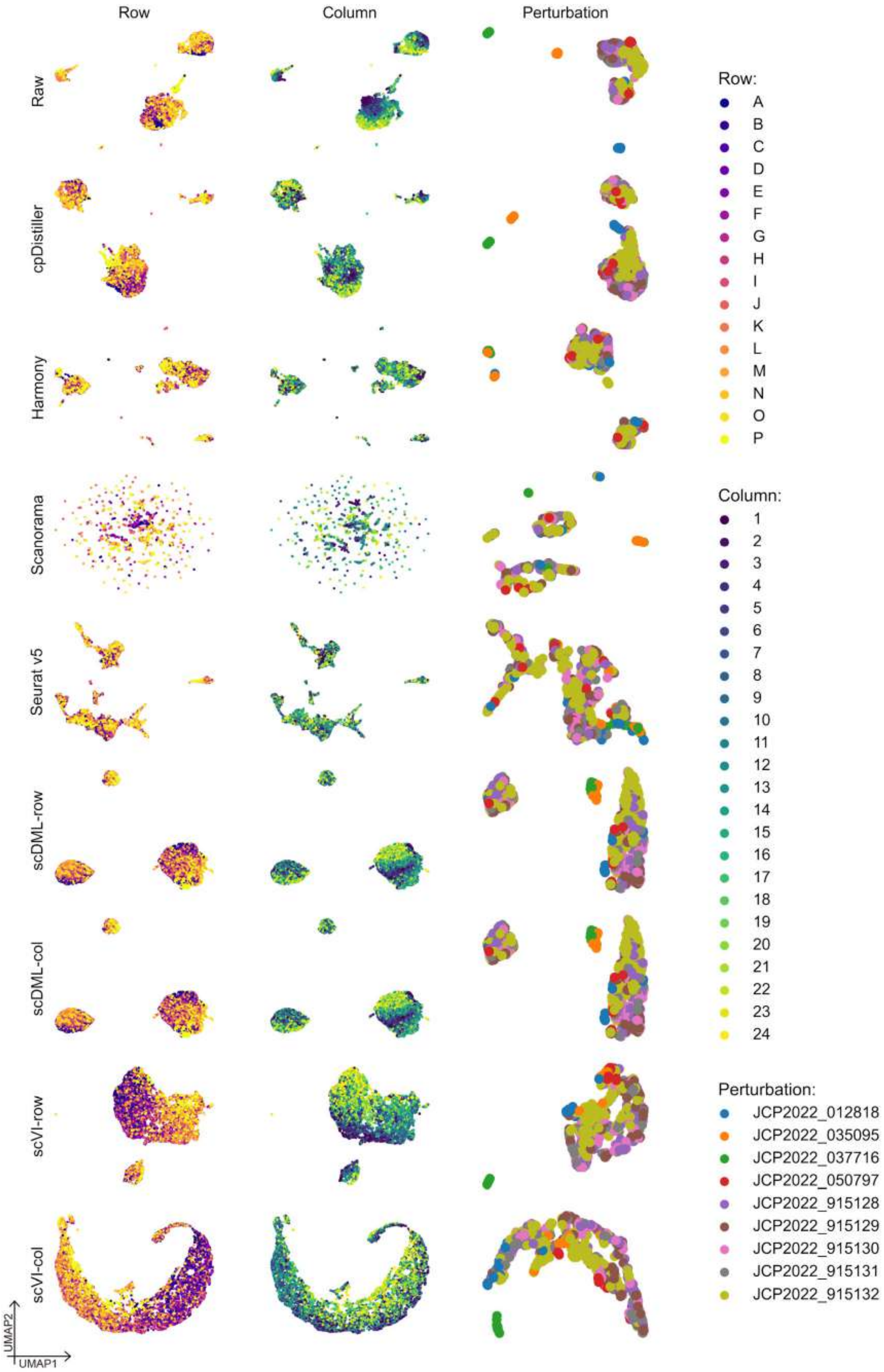

**Supplementary Figure 18. UMAP visualizations of embeddings obtained by different methods in Batch_5 of the ORF dataset.**

255

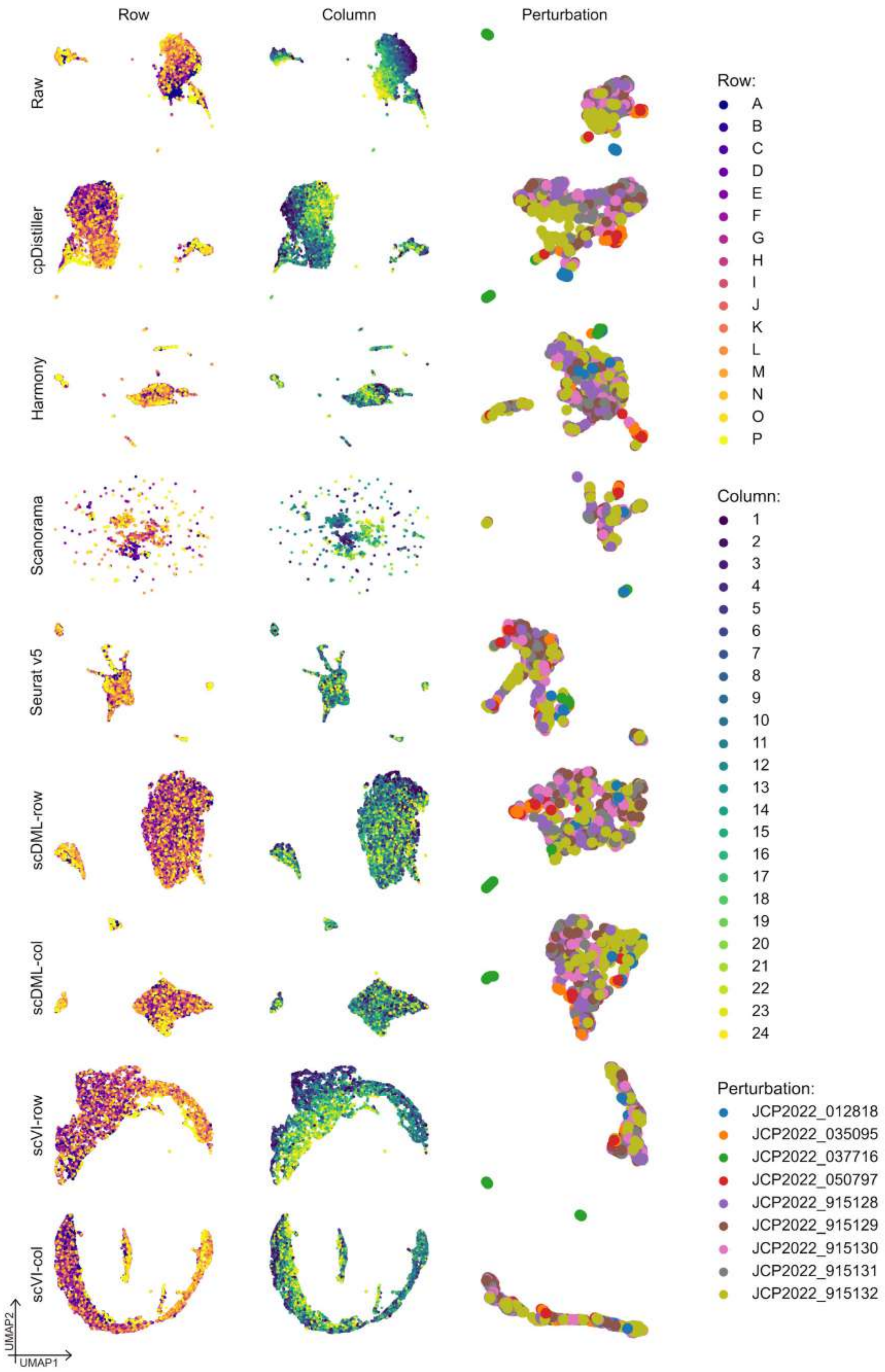

**Supplementary Figure 19. UMAP visualizations of embeddings obtained by different methods in Batch_6 of the ORF dataset.**

256

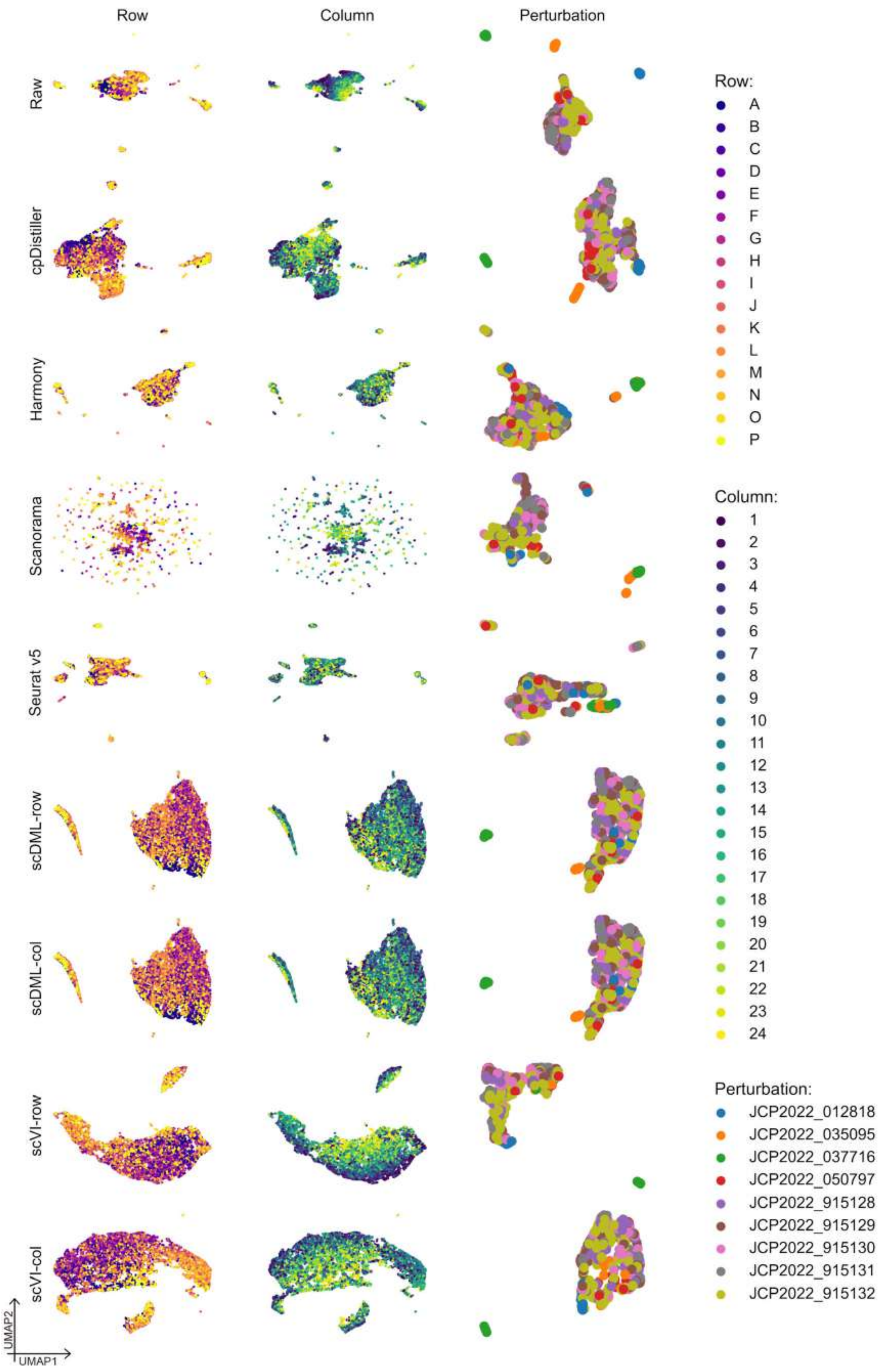

**Supplementary Figure 20. UMAP visualizations of embeddings obtained by different methods in Batch_7 of the ORF dataset.**

257

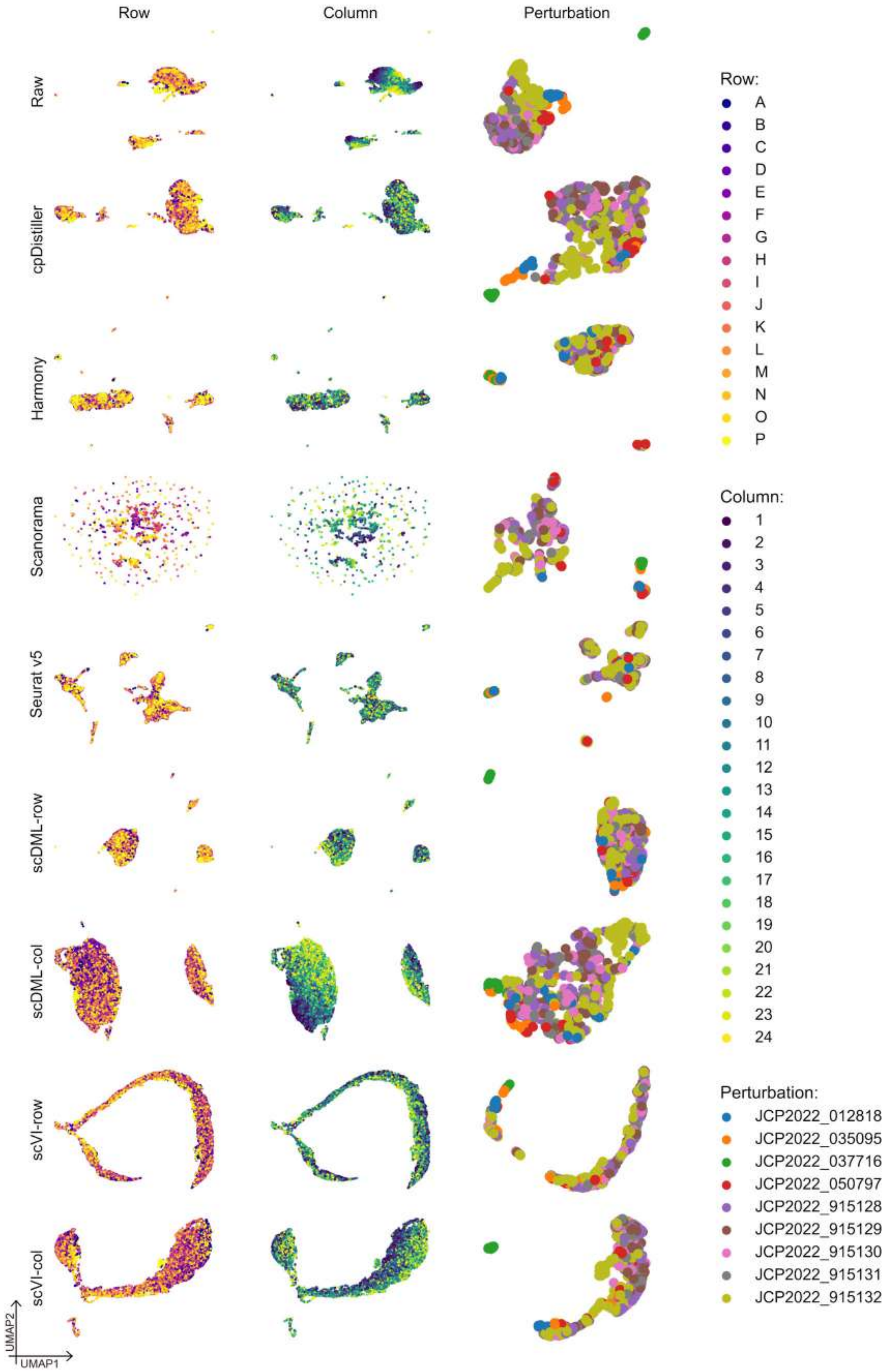

**Supplementary Figure 21. UMAP visualizations of embeddings obtained by different methods in Batch_8 of the ORF dataset.**

258

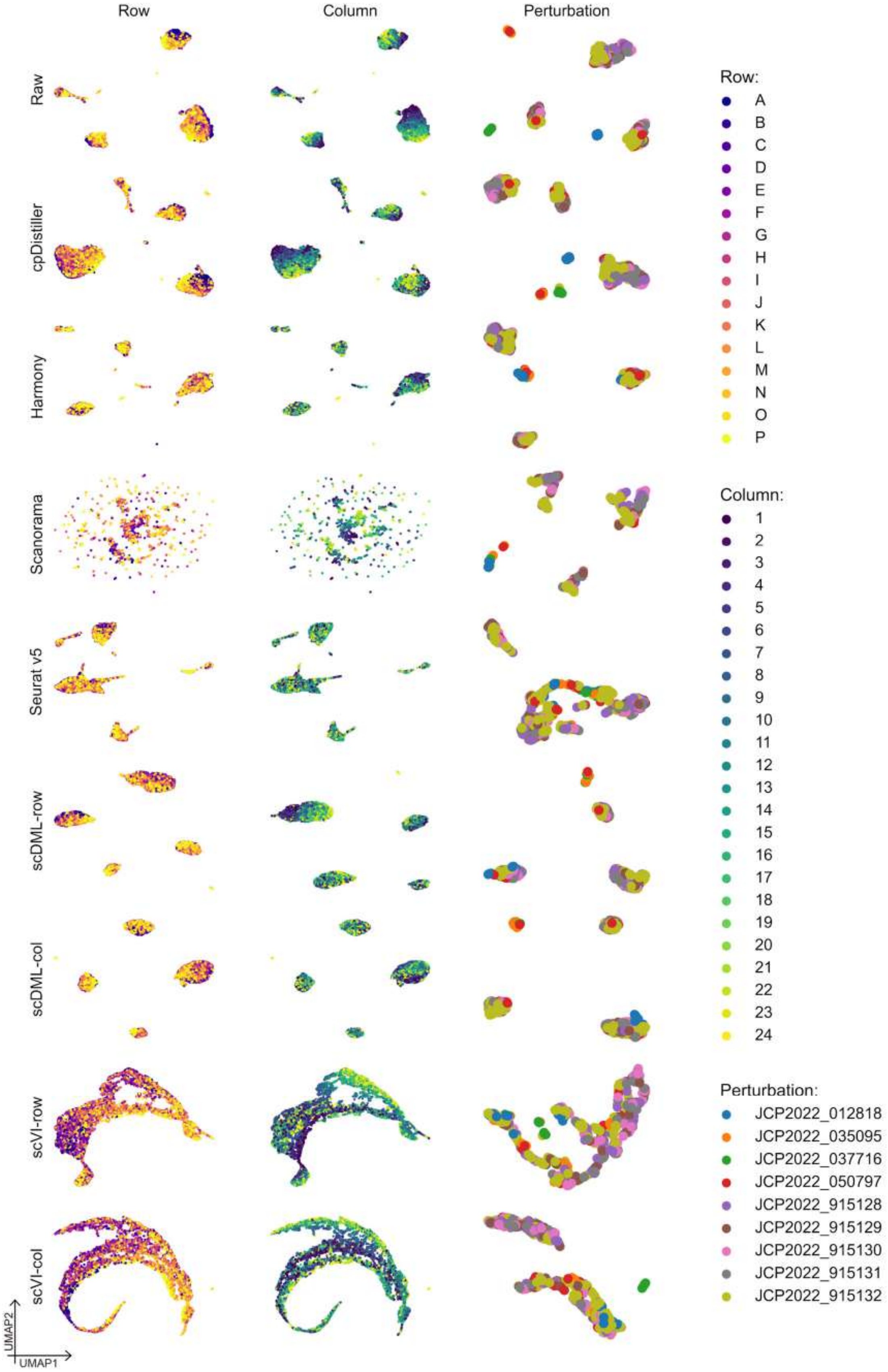

**Supplementary Figure 22. UMAP visualizations of embeddings obtained by different methods in Batch_9 of the ORF dataset.**

259

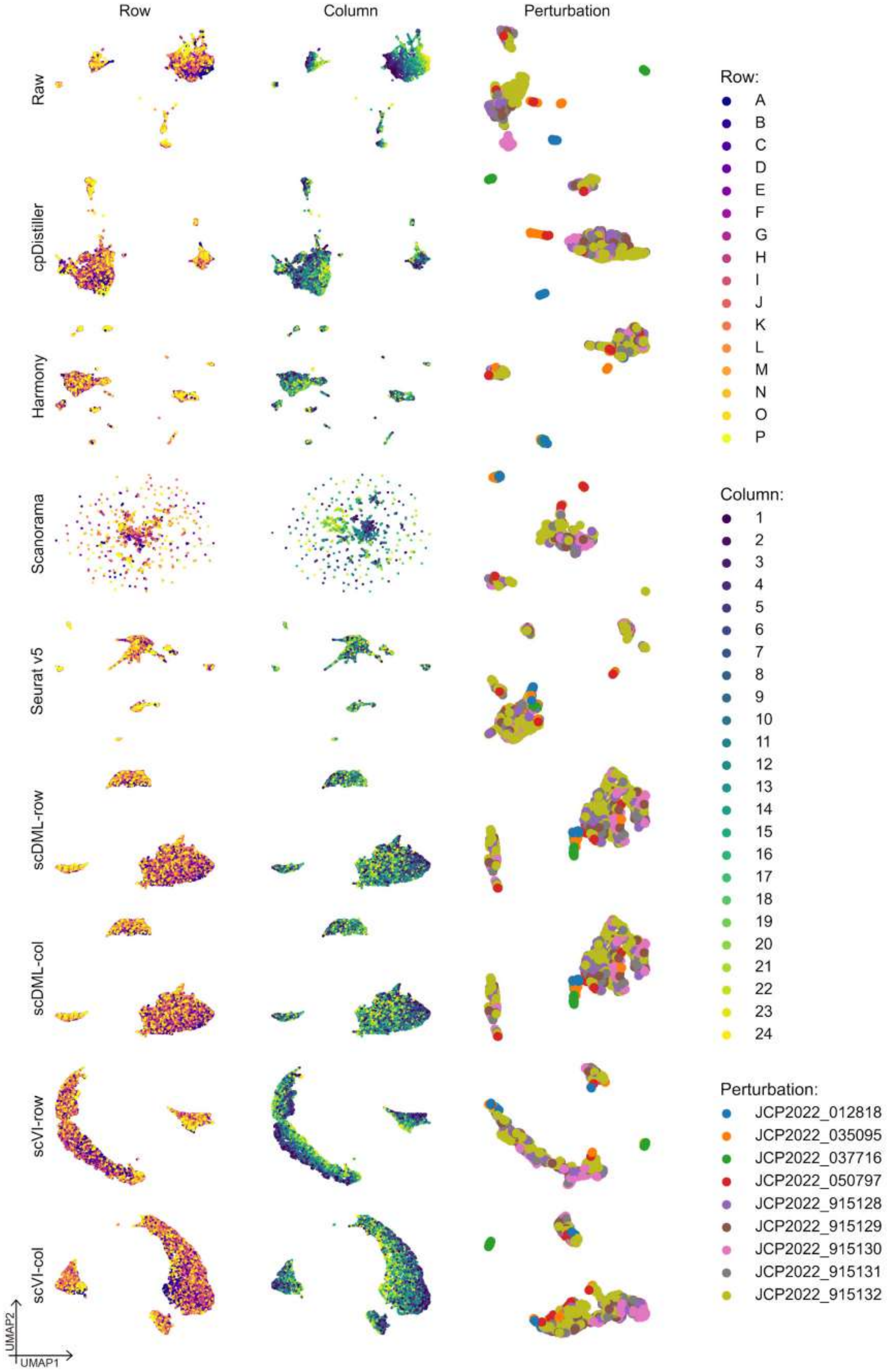

260 **Supplementary Figure 23. UMAP visualizations of embeddings obtained by different methods in Batch_10 of the ORF dataset.**

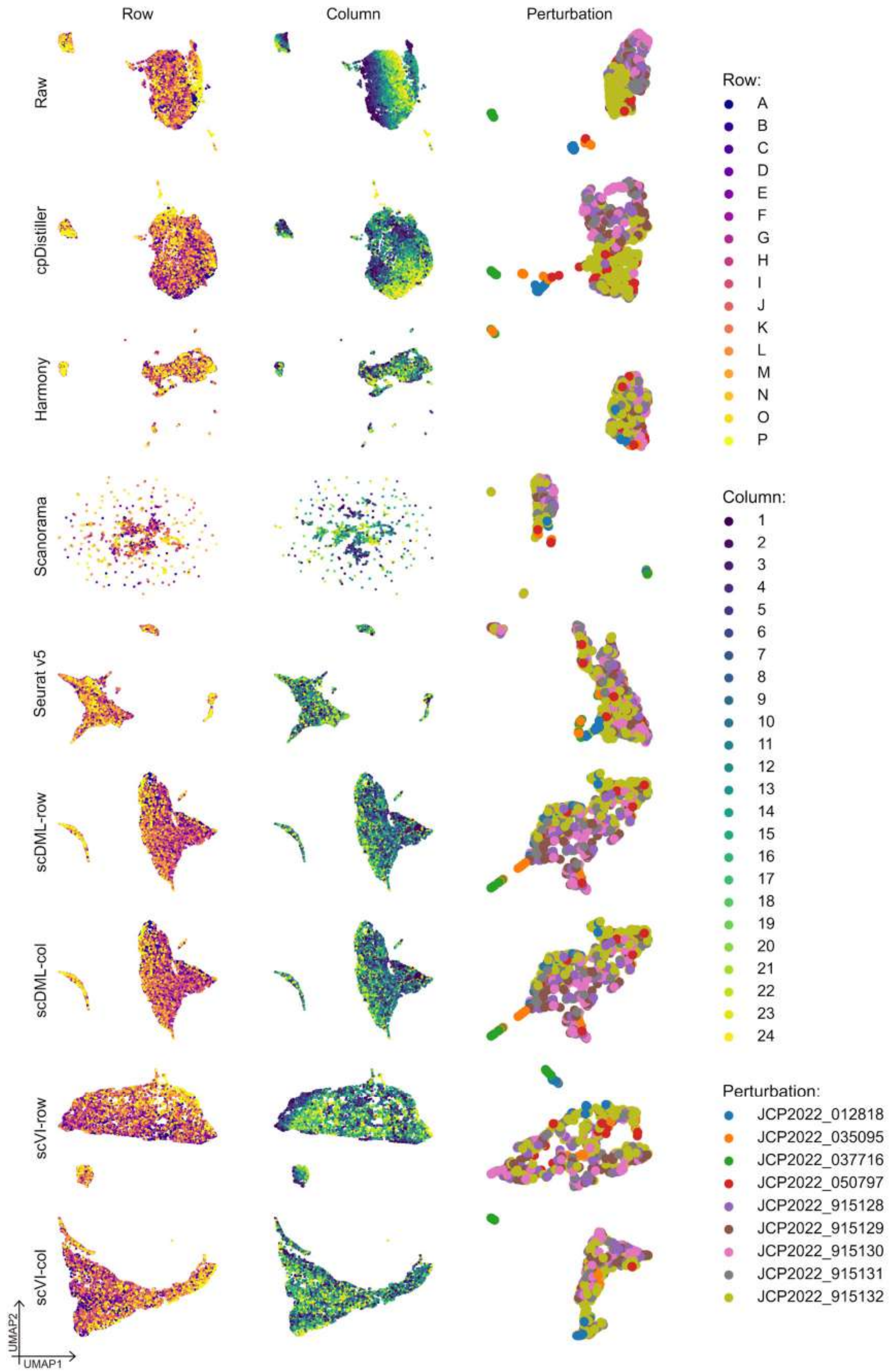

**Supplementary Figure 24. UMAP visualizations of embeddings obtained by different**
261 **methods in Batch_11 of the ORF dataset.**

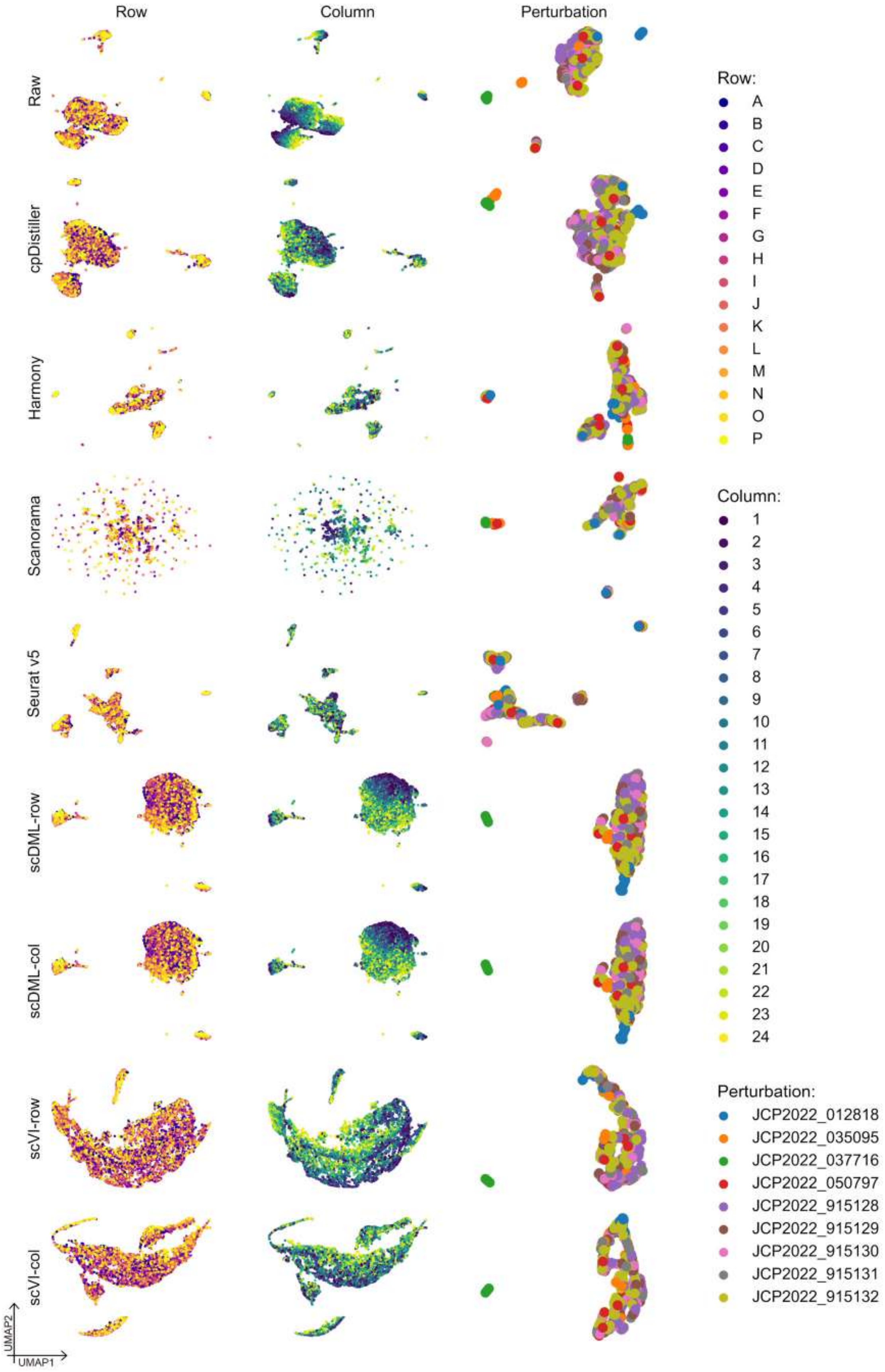

**Supplementary Figure 25. UMAP visualizations of embeddings obtained by different methods in Batch_13 of the ORF dataset.**

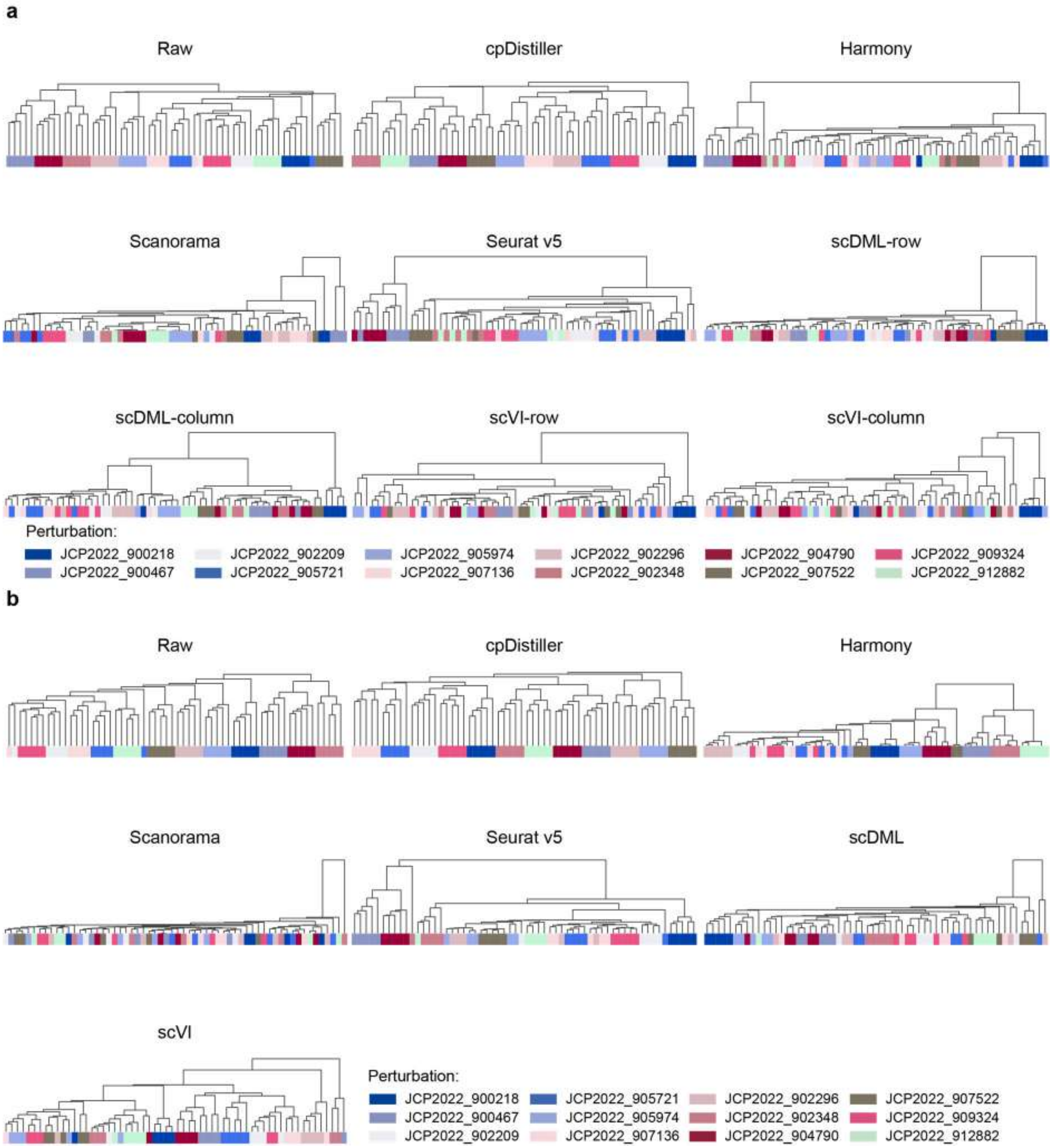

**Supplementary Figure 26. Dendrograms of hierarchical clustering for ORF perturbations using the embeddings learned by different methods. a,** Dendrograms are graphically rendered based on low-dimensional representations obtained by different methods, illustrating the clustering of ORF perturbations caused by 12 reagents in treatment, with correcting both row and column effects. **b,** Dendrograms are graphically rendered based on low-dimensional representations obtained by different methods, illustrating the clustering of ORF perturbations caused by 12 reagents in treatment, with correcting triple effects.

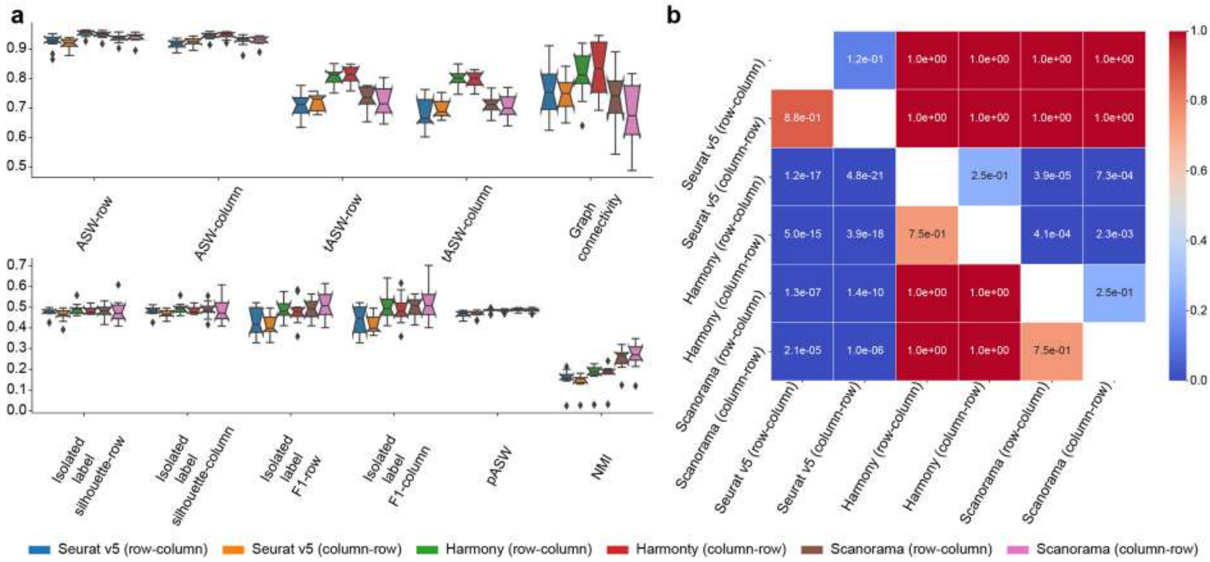

**Supplementary Figure 27. Impact of correction order on performance for different baseline methods a,** Box plots illustrate the impact of different correction orders on model performance. The top plot focuses on technical correction capabilities, while the bottom plot highlights biological preservation abilities. In the boxplots, center lines indicate the medians, box limits show upper and lower quartiles, whiskers represent 1.5× interquartile range, and notches reflect 95% confidence intervals via Gaussian-based asymptotic approximation. **b,** Heatmap shows *P*-values from one-sided paired Wilcoxon signed-rank tests. Each cell in the heatmap reflects the statistical significance of one method's (row) superiority over another over another (column), derived from 132 evaluations across 12 batches using 11 metrics.

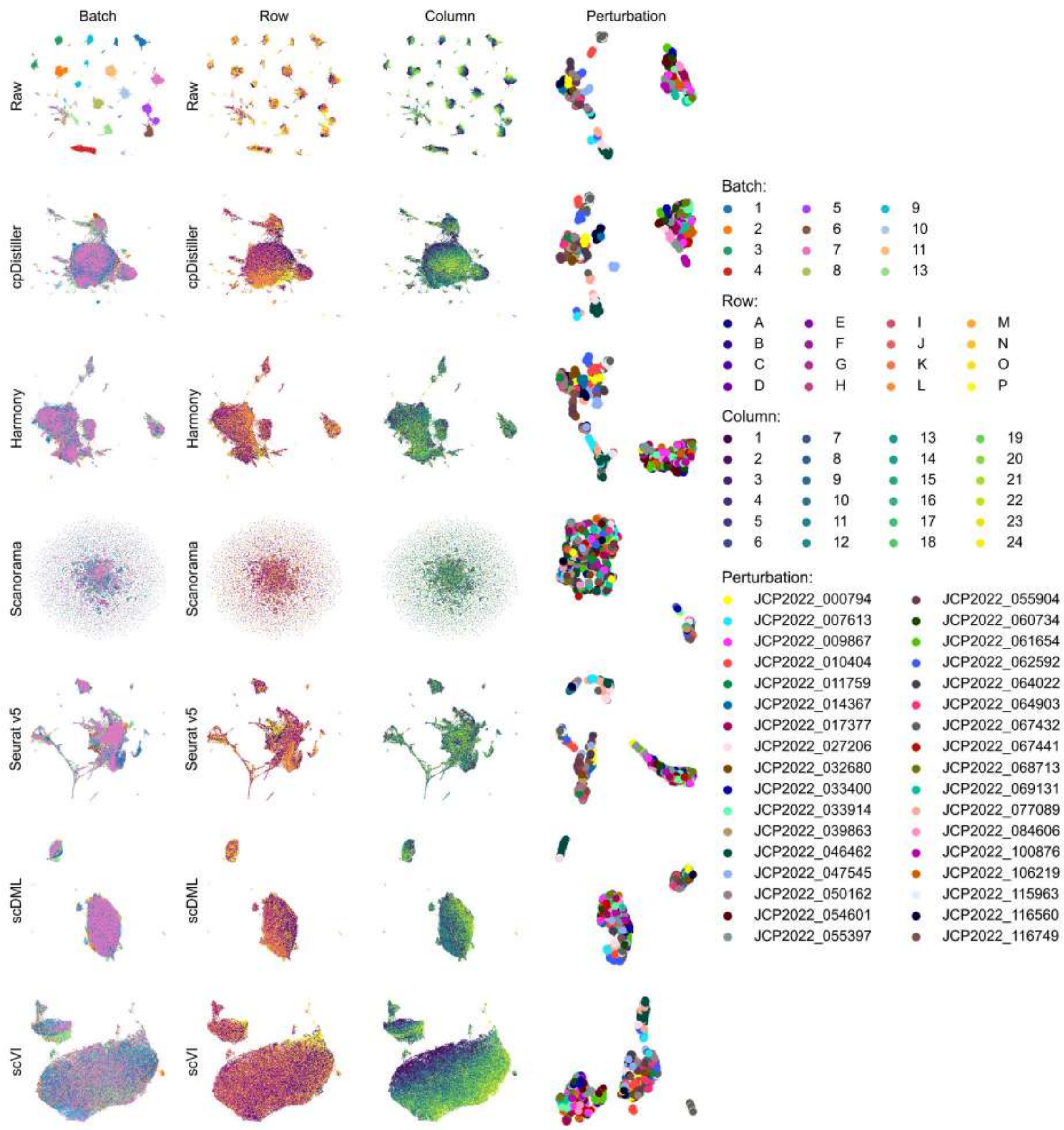

**Supplementary Figure 28. UMAP visualizations of embeddings obtained by different methods across all batches of the ORF dataset.**

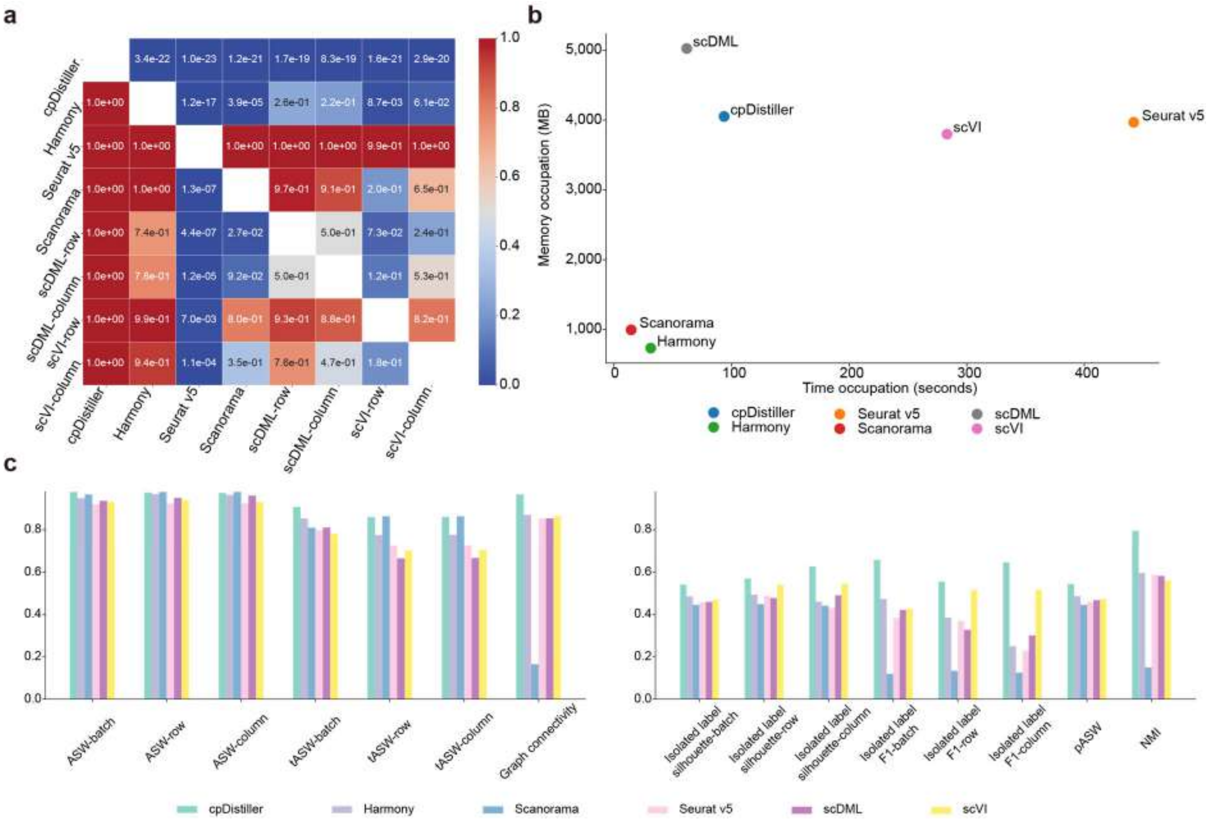

**Supplementary Figure 29. Benchmarks of different methods just using CellProfiler-based features. a,** Heatmap shows *P*-values from one-sided paired Wilcoxon signed-rank tests, with merely using CellProfiler-based features. Each cell in the heatmap reflects the statistical significance of one method's (row) superiority over another (column), derived from 132 evaluations across 12 batches using 11 metrics. **b,** Time and memory occupation of different methods for correcting well position effects in a single batch, with just using CellProfiler-based features. **c**, Bar chart illustrates the performance of different methods in removing triple effects and preserving biological variation, with only using CellProfiler-based features.

266

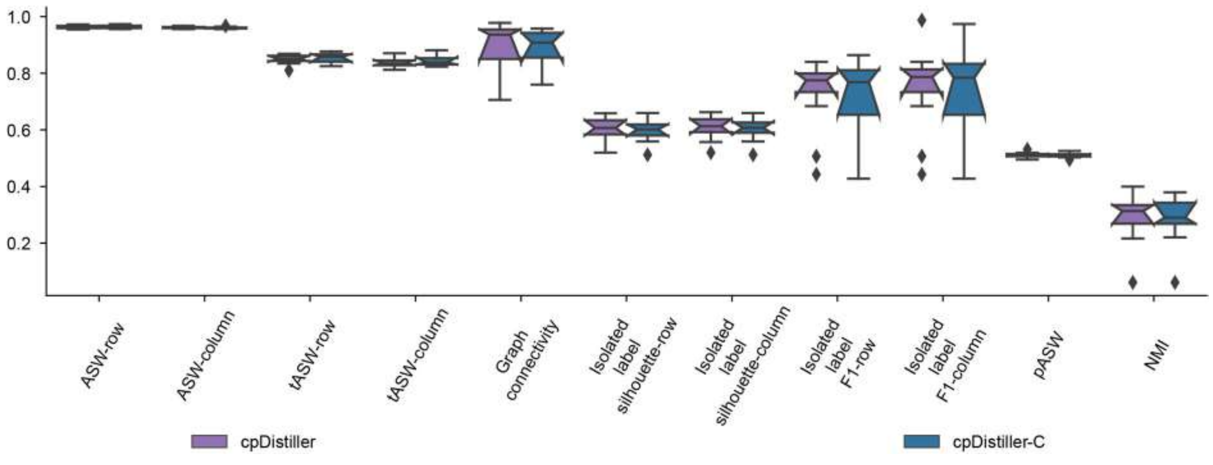

**Supplementary Figure 30. Ablation experiments on cpDistiller.** Quantitative evaluation of the performance of cpDistiller and cpDistiller-C on ORF profiles across 12 batches for correcting technical effects while preserving biological variation. In the boxplots, center lines indicate the medians, box limits show upper and lower quartiles, whiskers represent the 1.5× interquartile range, and notches reflect 95% confidence intervals via Gaussian-based asymptotic approximation.

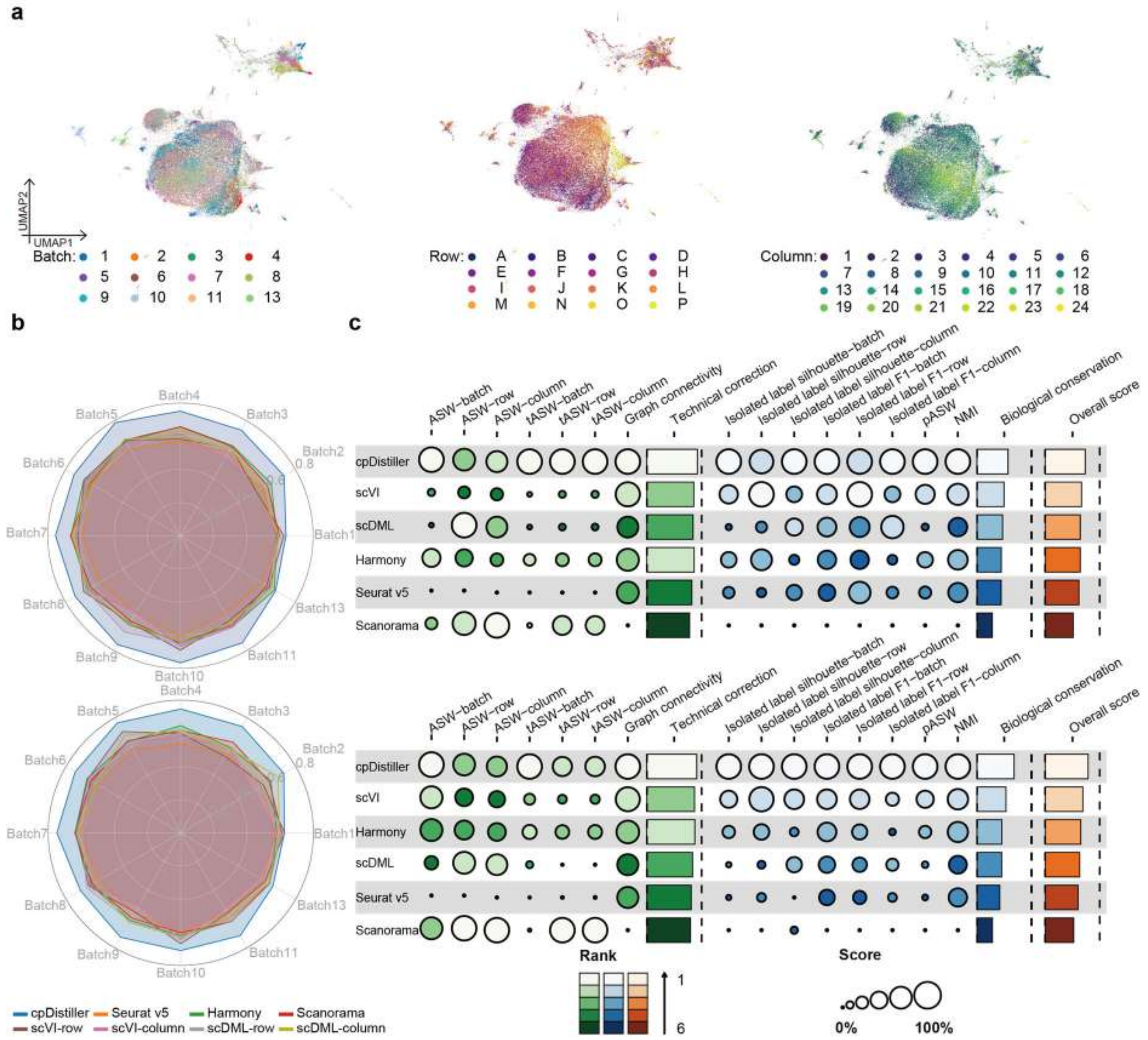

**Supplementary Figure 31. The extensive advantages of cpDistiller. a,** UMAP visualizations of embeddings obtained by cpDistiller trained on Batch1-Batch6, colored by batch, row and column, respectively. **b,c,** Top images illustrate results generated with the feature dimension of 4,752, whereas the bottom images correspond to results obtained using the feature dimension of 7,638. Radar plots show the average performance of different methods across 12 batches under two types of feature selection (**b**). Overview of benchmarking outcomes for different methods under two types of feature selection, focusing on two critical capabilities: removing triple effects and preserving biological variation (**c**). Technical correction and biological conservation scores refer to the average performance in these two aspects, whereas the overall score represents the aggregate performance across all metrics for different methods. Due to triple effects, which encompass batch, row and column effects, the metrics of ASW, tASW, isolated label scores yield three results, respectively.
